# ABCA7 Loss-of-Function Variants Impact Phosphatidylcholine Metabolism in the Human Brain

**DOI:** 10.1101/2023.09.05.556135

**Authors:** Djuna von Maydell, Shannon Wright, Ping-Chieh Pao, Colin Staab, Oisín King, Andrea Spitaleri, Julia Maeve Bonner, Liwang Liu, Chung Jong Yu, Ching-Chi Chiu, Daniel Leible, Aine Ni Scannail, Mingpei Li, Carles A. Boix, Hansruedi Mathys, Guillaume Leclerc, Gloria Suella Menchaca, Gwyneth Welch, Agnese Graziosi, Noelle Leary, George Samaan, Manolis Kellis, Li-Huei Tsai

## Abstract

Loss-of-function (LoF) variants in the lipid transporter ABCA7 significantly increase Alzheimer’s disease risk (odds ratio ≈ 2), yet the underlying pathogenic mechanisms and specific neural cell types affected remain unclear. To investigate this, we generated a single-nucleus RNA sequencing atlas of 36 human *postmortem* prefrontal cortex samples, including 12 carriers of ABCA7 LoF variants and 24 matched non-carriers. ABCA7 LoF variants were associated with transcriptional changes across all major neural cell types. Excitatory neurons, which expressed the highest levels of ABCA7, showed significant alterations in oxidative phosphorylation, lipid metabolism, DNA damage responses, and synaptic signaling pathways. ABCA7 LoF-associated transcriptional changes in neurons were similarly perturbed in carriers of the common AD missense variant ABCA7 p.Ala1527Gly (n = 240 controls, 135 carriers) predicted by molecular dynamic simulations to disrupt ABCA7 structure -, indicating that findings from our study may extend to large portions of the at-risk population. Human induced pluripotent stem cell (iPSC)-derived neurons carrying ABCA7 LoF variants closely recapitulated the transcriptional changes observed in human *postmortem* neurons. Biochemical experiments further demonstrated that ABCA7 LoF disrupts mitochondrial membrane potential via regulated uncoupling, increases oxidative stress, and alters phospholipid homeostasis in neurons, notably elevating saturated phosphatidylcholine levels. Supplementation with CDP-choline to enhance *de novo* phosphatidylcholine synthesis effectively reversed these transcriptional changes, restored mitochondrial uncoupling, and reduced oxidative stress. Additionally, CDP-choline normalized amyloid-*β* secretion and alleviated neuronal hyperexcitability in ABCA7 LoF neurons. This study provides a detailed transcriptomic profile of ABCA7 LoF-induced changes and highlights phosphatidylcholine metabolism as a key driver in ABCA7-induced risk. Our findings suggest a promising therapeutic approach that may benefit a large proportion of individuals at increased risk for Alzheimer’s disease.

## Introduction

Over 50 million people worldwide have dementia, with a large fraction of cases caused by Alzheimer’s disease [1]. Late-onset Alzheimer’s Disease (AD) affects individuals over the age of 65 and accounts for more than 95% of all AD cases [2]. Though AD is a multifactorial disorder, twin studies suggest a strong genetic component ( 70% heritability) [3] contributing to AD disease risk and progression. Large scale genome-wide association studies implicate multiple genes in AD etiology [4–10]. After APOE4, rare loss-of-function (LoF) mutations caused by premature termination codons (PTCs) in ATP-binding cassette transporter A7 (ABCA7), are among the strongest genetic factors for AD (odds ratio ≈ 2) [9, 11–15]. In addition to LoF variants, several common single nucleotide polymorphisms in ABCA7 depending on the population - moderately [9, 11–13, 16–18] to strongly [13] increase AD risk, suggesting that ABCA7 dysfunction may play a role in a significant proportion of AD cases. Despite the prevalence and potential impact of ABCA7 variants, the mechanism by which ABCA7 dysfunction increases AD risk remains poorly characterized.

ABCA7 is a member of the A subfamily of ABC transmembrane proteins [19] with high sequence homology to ABCA1, the primary lipid transporter responsible for cholesterol homeostasis and high-density lipoprotein genesis in the brain [20]. ABCA7 effluxes both cholesterol and phospholipids to APOA-I and APOE in *in vitro* studies [21–26] and has been shown to be a critical regulator of energy homeostasis, immune cell functions, and amyloid processing [27–32]. To date, study of ABCA7 LoF has been predominantly pursued in rodent knock-out models or in non-neural mammalian cell lines. These studies show that ABCA7 knock-out or missense variants cause increased amyloid processing and deposition [33–36], reduced plaque clearance by astrocytes and microglia [37, 38], and glial-mediated inflammatory responses [39, 40]. While these studies shed light on potential mechanisms of ABCA7 risk in AD, studies investigating the effects of ABCA7 LoF in human cells and tissue are severely lacking, with only a small number published to date [30, 36, 41, 42]. These human studies highlight a number of potential LoF-induced defects in human cells, including impacts on lipid metabolism and mitochondrial function [30]. However, comprehensive and unbiased profiling of multiple human neural cell types is needed to elucidate the mechanism by which ABCA7 LoF increases AD risk.

Single-nucleus RNA sequencing (snRNA-seq) of human neural tissue has identified cell type- specific transcriptional changes associated with AD risk variants in genes such as *APOE* and *TREM2* [43–47], providing insights into disease mechanisms and potential therapies. Here, we generated a cell type-specific transcriptomic atlas of ABCA7 LoF in the human prefrontal cortex (PFC). SnRNA-seq of *postmortem* brain tissue from ABCA7 LoF variant carriers and matched controls revealed widespread transcriptional alterations, particularly in excitatory neurons, which expressed the highest ABCA7 levels. Expression changes in these neurons indicated disruptions in lipid metabolism, mitochondrial respiration, DNA damage response, and synaptic function. Similar transcriptional changes were observed in neurons carrying the common missense variant p.Ala1527Gly, which was predicted to impair ABCA7 function based on structural simulations. This overlap indicates that p.Ala1527Gly may exert effects comparable to ABCA7 LoF, extending the relevance of our findings to a broader group of the at-risk population.

To complement our transcriptomic findings, we examined induced pluripotent stem cell (iPSC)-derived neurons harboring ABCA7 LoF variants. These neurons exhibited significant transcriptional overlap with human PFC neurons affected by ABCA7 LoF. Additionally, they demonstrated impaired uncoupled mitochondrial respiration, hyperpolarized mitochondrial membrane potential, elevated reactive oxygen species (ROS) levels, increased secretion of amyloid-*β* (A*β*), and hyperexcitability. Consistent with ABCA7’s known role in phospholipid transport, we also observed alterations in lipid composition, notably an increase in saturated phosphatidylcholine. Enhancing *de novo* phosphatidylcholine synthesis through CDP-choline supplementation effectively reversed these ABCA7 LoF-induced transcriptional changes and phenotypes. These findings link metabolic disruptions to AD pathology and suggest that neuronal ABCA7 may impact mitochondrial function through phosphatidylcholine imbalance, highlighting a potential mechanism by which ABCA7 variants increase AD risk.

## Results

### Single-nuclear transcriptomic profiling of human PFC from ABCA7 LoF-variant carriers

To investigate the cell type-specific impact of ABCA7 LoF variants in the human brain, we queried whole genome sequences of >1000 subjects from the Religious Order Study or the Rush Memory and Aging Project (collectively known as ROSMAP) for donors with Alzheimer’s disease diagnoses who are carriers of rare damaging variants in ABCA7 that result in a PTC. We identified 12 heterozygous carriers of ABCA7 LoF variants, including splice region variants (c.4416+2T>G and c.5570+5G>C), frameshift variants (p.Leu1403fs and p.Glu709fs), and nonsense ‘stop gained’ variants (p.Trp1245* and p.Trp1085*) (Figure 1A-C; Data S1). These variants have previously been associated with increased AD risk in genetic association studies (Table S1) [11, 14] and are presumed to induce risk via ABCA7 haploinsufficiency [48]. Analysis of published proteomic data for a subset of the 12 ABCA7 PTC-variant carriers and controls [49] (Table S6) confirmed that ABCA7 PTC-variant carriers indeed had lower ABCA7 protein levels in the human *postmortem* PFC compared to non-carriers (p=0.018; Figure 1D; Figure S1A).

**Figure 1:**
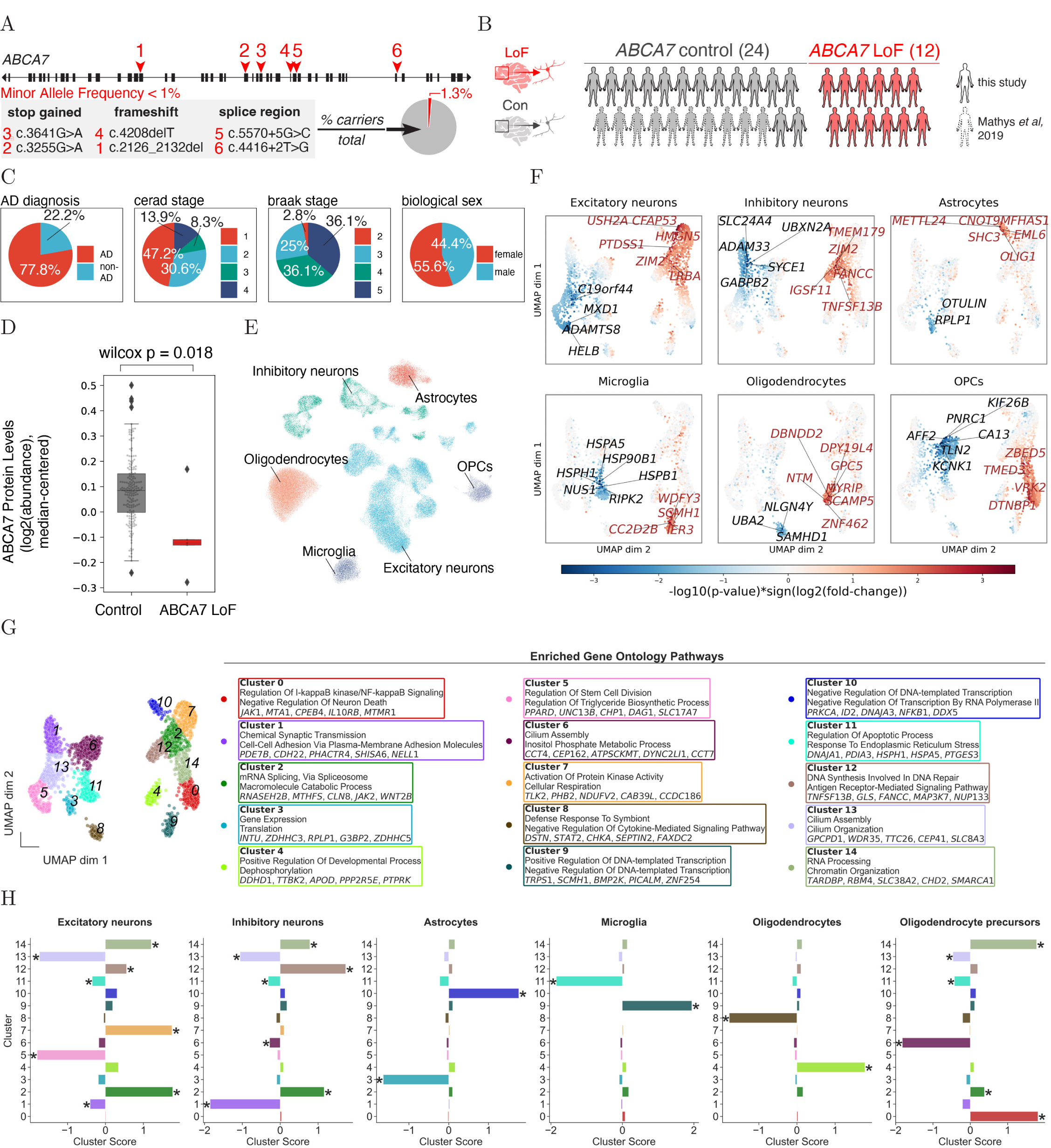
Single-nuclear RNA-sequencing Atlas of Human postmortem Prefrontal Cortex Reveals Cell Type-specific Gene Changes in ABCA7 LoF. **(A)** ABCA7 gene structure indicating variant locations studied here (average minor allele frequency <1%). Exons are black rectangles; introns, black lines. Pie chart indicates frequency of ABCA7 PTC-variant carriers in ROSMAP cohort. **(B)** Overview of human snRNA-seq cohort (created with BioRender.com). **(C)** Metadata summary of snRNA-seq cohort (*N* = 36 individuals). **(D)** ABCA7 protein abundance in postmortem prefrontal cortex from controls (*N* = 180) vs. ABCA7 LoF carriers (*N* = 5). Statistical comparison by Wilcoxon rank sum test. Boxes indicate quartiles; whiskers represent data within 1.5× interquartile range. **(E)** 2D UMAP projection of single-cell gene expression, colored by transcriptionally defined cell type. **(F)** 2D UMAP projection of ABCA7 LoF gene perturbation scores (*S* = − log_10_(*p*) × sign(log_2_(FC))). Red: *S >* 1.3, Blue: *S <* −1.3; point size reflects |*S*|. Up to top 10 genes labeled. **(G)** 2D UMAP projection colored by gene cluster assignment (Gaussian mixture model; see Methods). Top pathway enrichments per cluster shown (GO BP, hypergeometric enrichment, *p <* 0.01). Cell type-specific gene cluster scores (*SC* = mean(*S_i_*) for genes *i* in cluster *c*). * indicates permutation FDR-adjusted *p <* 0.01 and |*SC*| *>* 0.25.

We next selected 24 ABCA7 PTC non-carrier controls from the ROSMAP cohort that were matched to the ABCA7 LoF variant-carriers based on several potentially confounding variables, including Alzheimer’s disease (AD) pathology, age at death, *postmortem* intervals, sex, APOE genotype, and cognitive status (Figure 1C; Figure S1B,C; Data S2; Supplementary Text). We confirmed that none of the 36 selected subjects carried damaging variants in other known AD risk genes (*TREM2*, *SORL1*, *ATP8B4*, *ABCA1*, and *ADAM10* ) [14] and verified *ABCA7* genotypes in a subset of ABCA7 LoF carriers and matched controls using Sanger sequencing (Figure S1D).

For a subset of the selected samples, raw data (fastq files) for snRNAseq of the BA10 region of the prefrontal cortex (PFC) could be obtained from a previous study (10 non-carrier controls from [50]). For the remaining samples, fresh-frozen tissue samples from PFC BA10 were obtained for analysis. SnRNAseq was performed using the 10x Genomics Chromium platform. Accurate genotype assignments were confirmed by matching each single-cell library to its corresponding whole genome sequencing data (Figure S1E). Following extensive quality control measures—including detailed analysis and correction of batch effects (Figure S2; Data S3; Methods)—our final dataset consisted of 102,710 high-quality cells (Figure 1E), out of an initial total of 150,456 cells. This dataset encompassed diverse populations of inhibitory neurons (In, *SYT1* & *GAD1* +), excitatory neurons (Ex, *SYT1* & *NRGN* +), astrocytes (Ast, *AQP4* +), microglia (Mic, *CSF1R*+), oligodendrocytes (Oli, *MBP* & *PLP1* +), and oligodendrocyte precursor cells (OPCs, *VCAN* +) (Figure 1E; Figure S3A-E). A small putative vascular cell cluster did not meet our quality thresholds and was excluded from further analysis. Post-quality control, cell types were robustly represented across subjects (Figure S3F,G), and gene expression profiles showed high consistency within cell types (mean correlation 0.95) (Figure S3H,I).

### Cell type-specific perturbations in the presence of ABCA7 LoF

To investigate gene expression changes related to ABCA7 LoF across major cell types, we identified genes significantly perturbed (p<0.05, linear model; total genes = 2,389) in at least one of six major cell types (Ex, In, Ast, Mic, Oli, or OPC). We controlled for known and unknown covariates and considered only genes detected in >10% of cells within each specific cell type (Methods; Data S4). Next, we visualized these perturbed genes by projecting their high-dimensional perturbation scores (score = sign(log(FC)) × − log_10_(*p*-value) for each cell type) onto two dimensions, as shown in Figure 1F. Genes exhibiting similar perturbation patterns across cell types are positioned closer together in this two-dimensional visualization.

The two-dimensional visualization effectively captured the transcriptional landscape of ABCA7 LoF gene changes across all major cell types (Figure 1F; Figure S4A). To summarize this landscape in terms of biological pathways, we grouped genes into clusters based on their positions in the projection and analyzed each cluster for enrichment in biological pathways using the Gene Ontology Biological Process database (Figure 1G; Methods). This analysis identified several biological pathways correlated with ABCA7 LoF in the *postmortem* human PFC, including pathways related to cellular stress and apoptosis, synaptic function, DNA repair, and metabolism (Figure 1G; Data S5).

Decomposition of the ABCA7 LoF transcriptional signature revealed both shared and cell-specific gene perturbations across major PFC cell types (Figure 1G,H). Microglia exhibited significant downregulation of genes involved in cellular stress responses (*e.g.*, *HSPH1* ; cluster 11). A similar, though less pronounced, downregulation was observed in neurons and OPCs (FDR-adjusted *p <* 0.01, | score | *>* 0.25; Figure 1H). Microglia and astrocytes showed increased expression of transcriptional regulatory genes (clusters 9 and 10, respectively). OPCs and oligodendrocytes demonstrated alterations in inflammatory signaling pathways (*e.g.*, *IL10RB* in cluster 0 and *STAT2* in cluster 8; Figure 1H). Neurons displayed elevated expression of DNA repair genes (*e.g.*, *FANCC* ; cluster 12) and reduced expression of synaptic transmission genes (*e.g.*, *NLGN1*, *SHISA6* ; cluster 1). Excitatory neurons uniquely exhibited enhanced expression of genes involved in cellular respiration (*e.g.*, *NDUFV2* ; cluster 7) and reduced expression of genes related to triglyceride biosynthesis (*e.g.*, *PPARD*; cluster 5; Figure 1H). Overlapping differentially expressed genes across cell types are summarized in Figure S5A,B.

Together, these findings indicate that ABCA7 LoF variants may induce widespread, cell type-specific transcriptional changes in the human PFC. This single-cell atlas provides a rich resource for future studies aiming to elucidate the contributions of individual neural cell types to ABCA7 LoF-driven forms of AD risk. This resource will be made available for exploration via the UCSC Single Cell Browser and for further analysis via Synapse (accession ID: syn53461705).

### ABCA7 is expressed most highly in excitatory neurons

Our snRNAseq data suggest that excitatory neurons expressed the highest levels of ABCA7, compared to other major cell types in the brain (Figure S6A). ABCA7 transcripts were detected (count>0) in 30% of excitatory neurons and 15% of inhibitory neurons, while the detection rate was considerably lower (<10%) for microglia and astrocytes and an order of magnitude lower (<3%) for oligodendrocytes and OPCs (Figure S6A, B). We validated this expression pattern in an independent published dataset [51] (Table S6), where bulk RNA sequencing of NeuN- (glial) and NeuN+ (neuronal) cell populations derived from six human *postmortem* temporal cortex samples showed significantly higher ABCA7 levels in the neuronal population versus the glial cell population (p=0.021; Figure S6C). Several control genes, whose expression patterns in glial versus neuronal cells are well established (*ABCA1*, *APOE*, and *NEUROD1* ), had expected expression patterns that matched those in the snRNAseq data (Figure S6B,C). These results indicate that neurons, particularly excitatory neurons, are the primary ABCA7-expressing cell type in the aged human PFC. Given the relatively higher expression of ABCA7 in excitatory neurons and the evidence of transcriptional perturbations by ABCA7 LoF in this cell type, we focused our subsequent analysis specifically on excitatory neurons.

### ABCA7 LoF perturbations in excitatory neurons

As an alternative approach to the unsupervised clustering of gene perturbation scores among all cell types, we next used prior knowledge of biological pathway structure to perform an in-depth characterization of perturbed biological processes specifically in ABCA7 LoF excitatory neurons. To this end, we first estimated statistical overrepresentation of biological gene sets (WikiPathways, N pathways = 472) among up and down-regulated genes in ABCA7 LoF excitatory neurons vs controls (by GSEA; Methods). We observed a total of 34 pathways with evidence for transcriptional perturbation at p<0.05 in excitatory neurons (Data S6). Enrichments of these pathways were driven by 268 unique genes (“leading edge” genes [52]; Data S6).

To extract unique information from leading-edge genes and limit pathway redundancy, we next separated these genes and their associated pathway annotations into non-overlapping groups, formalized as a graph partitioning problem (Figure 2A; Figure S7; Methods; Supplementary Text). Establishing gene-pathway groupings of approximately equal size revealed eight biologically interpretable “clusters” associated with ABCA7 LoF in excitatory neurons (Figure 2A,B; Data S7). Predominantly, these gene clusters centered around two themes: (1) energy metabolism and homeostasis (PM.0, PM.1) and (2) DNA damage (PM.2, PM.3), cell stress (PM.4, PM.5), and synaptic dysfunction (PM.7) (Figure 2B). Cortical layer-specific analysis indicated that these perturbation patterns remained largely consistent across cortical layers, from deeper to superficial regions (Figure S8).

**Figure 2:**
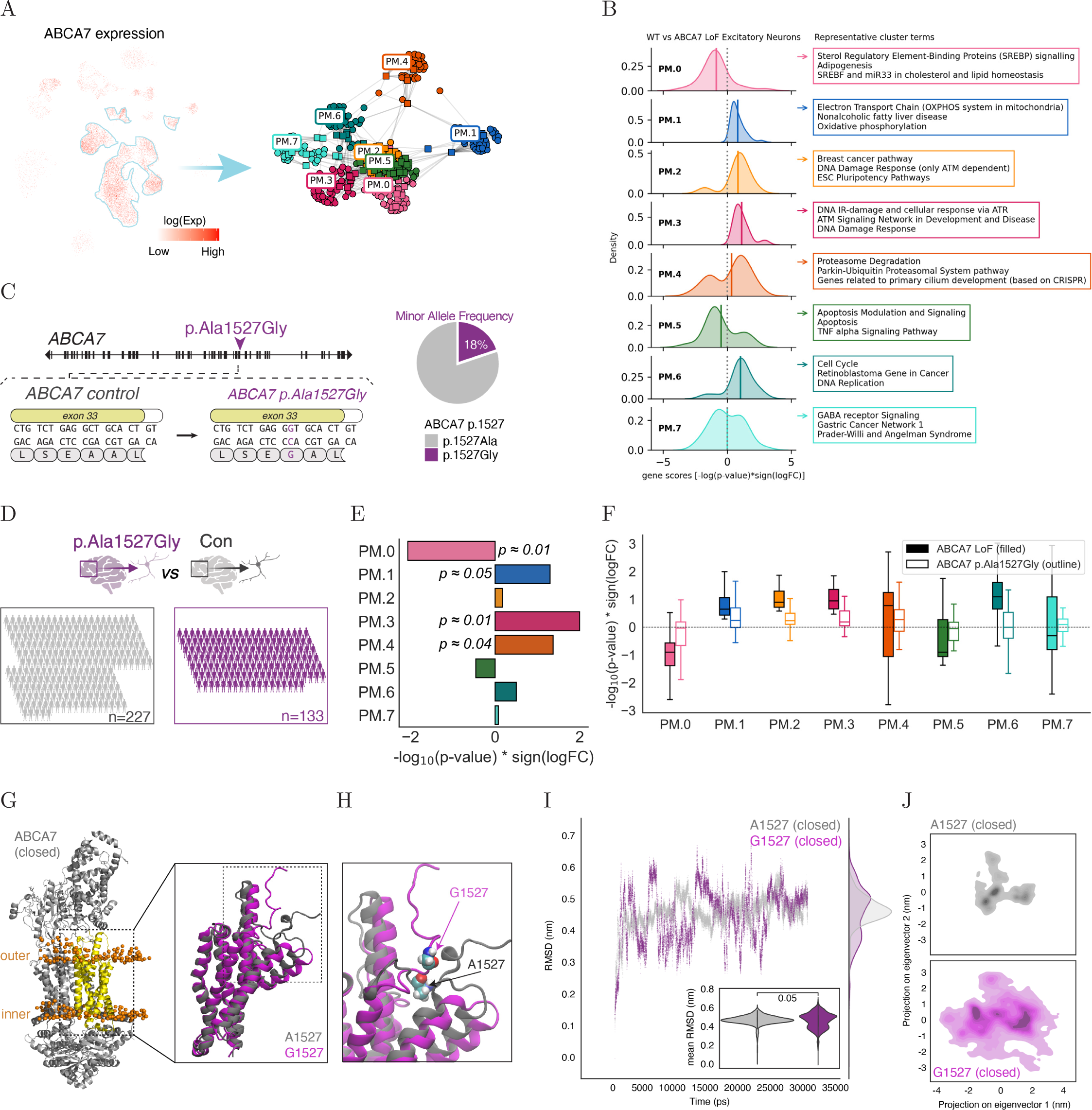
Transcriptional Perturbations in Excitatory Neurons in ABCA7 LoF and ABCA7 p.Ala1527Gly Variant Carriers. **(A)** (left) 2D UMAP projection of all cell types colored by log-transformed values of log-normalized ABCA7 expression (log(Exp)). (right) Kernighan-Lin (K/L) clustering of leading-edge genes from significantly perturbed pathways (*p <* 0.05) in ABCA7 LoF excitatory neurons. Colors indicate distinct K/L clusters (0–7). **(B)** Gaussian kernel density plots of gene perturbation scores (*S* = − log_10_(*p*) × sign(log_2_(FC))) per K/L cluster. Positive *S* indicates upregulation in ABCA7 LoF. Solid lines show distribution means. Representative pathways with highest intra-cluster connectivity annotated per cluster. **(C)** Schematic of ABCA7 gene highlighting the p.Ala1527Gly codon change (purple arrow). Minor allele frequency (MAF) shown at right. **(D)** Overview of snRNA-seq cohort comparing ABCA7 p.Ala1527Gly carriers (homozygous/heterozygous) to non-carrier controls (MAF ≈ 18%). **(E)** Perturbation (FGSEA scores) of ABCA7 LoF-associated gene clusters from (B) in excitatory neurons from p.Ala1527Gly carriers vs. controls. Top *p*-values (*p <* 0.1) indicated. Positive scores represent upregulation in carriers. **(F)** Distribution of gene perturbation scores (*S*) for each K/L cluster comparing ABCA7 p.Ala1527Gly (no fill) vs. LoF variants (solid fill). Positive *S* indicates upregulation. **(G)** Closed-conformation ABCA7 protein structure, highlighting domain (residues 1517–1756, yellow) used for molecular simulations. Lipid bilayer shown in orange. Expanded inset highlights Ala1527 (light grey) and Gly1527 (purple) residues. **(H)** Expanded inset from (G) with residues of interest indicated. **(I)** Root mean squared deviations (RMSD) of the closed-conformation ABCA7 domain (G) carrying Ala1527 (light grey) or Gly1527 (purple) during simulation, relative to reference closed conformation. Inset violin plot shows average *C_α_* atom positional fluctuations. Projection of *C_α_* positional fluctuations onto the first two principal components during simulation for Ala1527 (top, light grey) and Gly1527 (bottom, purple).

Clusters PM.0 and PM.1 were primarily defined by genes involved in cellular energetics, including genes related to lipid metabolism, mitochondrial function, and oxidative phosphorylation (OXPHOS) (Figure 2B). Cluster PM.0, characterized by transcriptional regulators of lipid homeostasis (*e.g., NR1H3*, *ACLY*, *PPARD*), exhibited evidence for down-regulation in ABCA7 LoF and featured pathways related to “SREBP Proteins” and “Adipogenesis” (Figure 2B; Data S7). Cluster PM.1 comprised multiple mitochondrial complex genes (*e.g., COX7A2*, *NDUFV2* ) responsible for ATP generation from carbohydrate and lipid catabolism and showed up-regulation in ABCA7 LoF (Figure 2B; Data S7). Clusters PM.2-6 were characterized by DNA damage and proteasomal, inflammatory, and apoptotic mediators. Clusters PM.2, PM.3, and PM.6 were up-regulated in ABCA7 LoF excitatory neurons and characterized by pathway terms such as “DNA Damage Response” (PM.2 & PM.3) and “DNA Replication” (PM.6) (Figure 2B; Data S7). They included up-regulated DNA damage/repair and proteasomal genes (*e.g., RECQL*, *TLK2*, *BARD1*, *RBL2*, *MSH6*, *PSMD5* ). Genes in clusters PM.4, linked to “Proteasome Degradation” and “ciliogenesis”; PM.5, associated with “Apoptosis” and “TNFalpha Signaling Pathway”; and PM.7, linked to “GABA receptor Signaling,” “Gastric Cancer Network 1,” and “Prader-Willi and Angelman Syndrome”, included both up- and down-regulated genes (Figure 2B; Data S7).

Together, these data suggest that ABCA7 LoF may disrupt energy metabolism in excitatory neurons and that these disruptions coincide with a state of increased cellular stress, characterized by genomic instability and neuronal dysfunction.

### ABCA7 LoF and common missense variants lead to overlapping neuronal perturbations

ABCA7 LoF variants substantially increase AD risk (Odds Ratio = 2.03) [11] but are rare and therefore only contribute to a small portion of AD cases [48]. To evaluate whether ABCA7 LoF transcriptomic effects in neurons generalize to more common, moderate-risk genetic variants in ABCA7, we examined the ROSMAP WGS cohort for carriers of the prevalent ABCA7 missense variant p.Ala1527Gly (rs3752246: Minor Allele Frequency ≈ 0.18; % carriers 1 allele ≈30%; Figure 2C). Although Gly1527 is listed as the reference allele, it represents the less common variant associated with increased AD risk (Odds Ratio = 1.15 [1.11-1.18]) [7, 14, 18]. We identified 133 individuals carrying at least one copy of the p.Ala1527Gly risk variant and 227 non-carriers (Figure 2D), all with available snRNAseq data from *postmortem* PFC [53]. We ensured that none of these 360 individuals were part of our earlier ABCA7 LoF snRNAseq cohort or carried ABCA7 LoF variants. Using this cohort, we investigated whether excitatory neurons from p.Ala1527Gly carriers exhibited evidence of transcriptomic perturbations in the ABCA7 LoF-associated clusters PM.0-7.

Remarkably, all clusters displayed directional trends in p.Ala1527Gly neurons consistent with the directionality observed in ABCA7 LoF neurons (Figure 2B,E,F), while controlling for pathology, age, sex, and other covariates (Methods). Notably, 4 out of 8 clusters exhibited substantial evidence of perturbation in p.Ala1527Gly variant carriers, with perturbation directions aligning with predictions for ABCA7 LoF (Figure 2E,F). Specifically, we observed an up-regulation in the DNA damage cluster PM.3 and the proteasomal cluster PM.4 in p.Ala1527Gly carriers compared to controls, suggesting a similar cell stress and genomic instability signature to ABCA7 LoF carriers (Figure 2E,F), and a borderline significant upregulation of the mitochondrial cluster PM.1, again consistent with ABCA7 LoF (Figure 2E,F). Finally, we observed significant perturbation to the lipid cluster PM.0, which was downregulated (Figure 2E,F) similar to our observations in ABCA7 LoF carriers.

Because missense variants often influence protein dynamics—and glycine substitutions typically introduce greater local flexibility than alanine—we next examined whether the convergent transcriptional signature associated with ABCA7 variants could be explained by structural changes in the protein. To directly investigate local structural consequences of the p.Ala1527Gly variant, we performed molecular dynamics simulations using newly available cryo-EM structures of ABCA7 in both the ATP-bound closed (Figure 2G,H) and ATP-unbound open (Figure S9A,B) conformations [54, 55]. Specifically, simulations were conducted on a 239-residue region of ABCA7 embedded within a lipid bilayer, comparing the Ala1527 and Gly1527 variants over a 300-ns timescale (Figure 2I; Figure S9C; Figure S10; Methods; Supplementary Text).

Our simulations revealed that the AD risk-associated Gly1527 variant increased local structural flexibility in the ATP-bound closed conformation, indicated by pronounced conformational fluctuations over time (Figure 2J). In contrast, the Ala1527 variant exhibited limited conformational fluctuations, suggesting minimal local structural flexibility in the closed state (Figure 2J). Both variants demonstrated stable conformational behavior in the ATP-unbound open state (Figure S9C-E). These results are further supported by analyses of *ϕ*/*ψ* dihedral angle distributions and secondary structure persistence, as detailed in the Supplementary Text (Figure S10).

Together, these data suggest that the Gly1527 variant may introduce increased local flexibility, potentially disrupting the stability of secondary structural elements specifically within the ATP-bound closed conformation. Given that this conformation is proposed to mediate lipid presentation to apolipoproteins [26, 54], the p.Ala1527Gly substitution may impact the efficiency of lipid extrusion, consistent with recent experimental findings from [26]. Combined with our transcriptomics analyses, these structural insights suggest that both rare, high-effect ABCA7 LoF variants and common, mild-effect variants may influence AD risk through similar ABCA7-dependent mechanisms, indicating that our in-depth studies of rare variants may generalize to broader at-risk populations.

### Deriving human neurons with ABCA7 LoF variants

To complement the correlative analyses in ABCA7 LoF human tissue, we next used CRISPRCas9 genome editing to generate two isogenic iPSC lines, each homozygous for a different ABCA7 LoF variant, from a parental line without ABCA7 variants (WT). The first LoF variant, ABCA7 p.Glu50fs*3, was generated by a single base-pair insertion in ABCA7 exon 3, resulting in a PTC early in the ABCA7 gene (Figure 3A; Figure S11A-C). The second LoF variant, ABCA7 p.Tyr622*, was generated by a single base-pair mutation in ABCA7 exon 15 (Figure 3A; Figure S11A-C). This PTC re-creates a variant previously observed in patients as associated with AD [11] and thus provides clinical context to ABCA7 dysfunction. Both variants are expected to generate severely truncated ABCA7 proteins or, due to nonsense-mediated mRNA decay, no ABCA7 protein at all. However, transcript rescue from nonsense-mediated decay and possible generation of mutated forms of ABCA7 through mechanisms such as exon skipping, which have previously been reported for multiple ABCA7 LoF variants [56], cannot be excluded.

**Figure 3:**
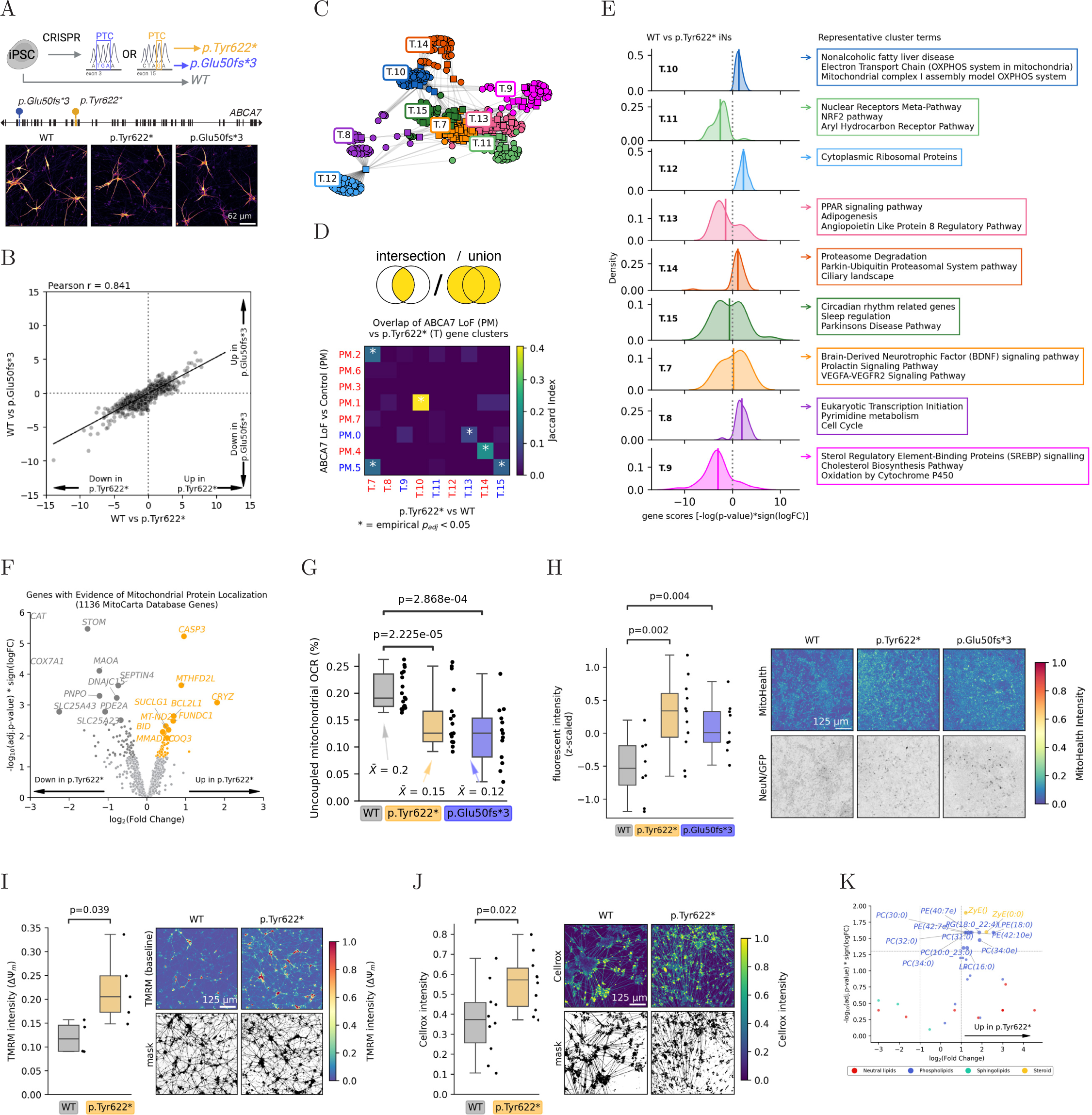
ABCA7 LoF Impacts Regulation of Mitochondrial Uncoupling in Neurons. **(A)** Schematic of iPSC-derived isogenic neuronal lines harboring ABCA7 loss-of-function (LoF) variants. Gene structure shows exons (black rectangles) and introns (black lines). CRISPR-Cas9 introduced premature termination codons in exon 3 (p.Glu50fs3, blue) or exon 15 (p.Tyr622*, orange). Confocal images show MAP2 staining in iNs differentiated for 4 weeks (genotypes indicated). **(B)** Correlation of gene perturbation scores (*S* = − log_10_(*p*)×sign(log_2_(FC))) by bulk mRNAseq comparing p.Glu50fs3 vs. WT and p.Tyr622* vs. WT iNs cultured for 4 weeks. **(C)** Kernighan-Lin (K/L) clustering of leading-edge genes from significantly perturbed pathways (Ben-jamini–Hochberg (BH) FDR-adjusted *p <* 0.05) in p.Tyr622* vs. WT iNs. Colors represent distinct K/L clusters. **(D)** Heatmap of Jaccard index overlap between K/L gene clusters from p.Tyr622* neurons and clusters identified in human postmortem excitatory neurons. Red text denotes clusters with average score *S* upregulated in ABCA7 LoF; blue text denotes clusters with average *S* downregulated in ABCA7 LoF. **(E)** Gaussian kernel density plots of gene perturbation scores (*S*) within each cluster. Positive *S* indicates upregulation in p.Tyr622*. Solid lines denote cluster means. Top enriched pathways with highest intra-cluster connectivity indicated. **(F)** Volcano plot of differential expression of genes with mitochondrial-localized protein products (MitoCarta) between p.Tyr622* and WT neurons. **(G)** Seahorse-measured mitochondrial uncoupled oxygen consumption rate (OCR) in WT and ABCA7 LoF and WT iNs cultured for 4 weeks. Each datapoint represents OCR from a single well. *N* = 18 (WT), 17 (p.Tyr622*), 13 (p.Glu50fs3) wells, across two differentiation batches. Statistical comparison by independent-sample *t*-test. **(H)** Mitochondrial membrane potential quantified via HCS MitoHealth dye fluorescence intensity in ABCA7 LoF iNs cultured for 4 weeks. Each datapoint represents average intensity per well (NeuN+ volumes averaged). Statistical comparison via linear mixed-effects model, accounting for well-of-origin random effects. *N* = 8 (WT), 11 (p.Tyr622*), 9 (p.Glu50fs3) wells; ≈ 3000 cells/condition, from three differentiation batches. Each NeuN/GFP image intensity was scaled relative to its maximum value, followed by gamma correction (*γ* = 0.5) for visualization. **(I)** Baseline mitochondrial membrane potential quantified by average TMRM fluorescence intensity per masked region (thresholded at 75th percentile) in ABCA7 LoF and WT iNs cultured for 4 weeks. Each datapoint represents average intensity per well. *N* = 4 (WT), 5 (p.Tyr622*) wells. Statistical comparison by independent-sample *t*-test. **(J)** Oxidative stress quantified by average CellROX fluorescence intensity per masked region (thresholded at 75th percentile) in p.Tyr622* and WT iNs cultured for 4 weeks. Each datapoint represents average intensity per well. *N* = 10 wells per genotype. Statistical comparison by independent-sample *t*-test. Volcano plot of differentially abundant lipid species between p.Tyr622* and WT iNs cultured for 4 weeks, colored by lipid class. Statistical comparisons by independent-sample *t*-tests followed by BH FDR adjustment. *N* = 10 wells (WT) and 8 wells (p.Tyr622*).

We differentiated isogenic iPSCs into neurons (iNs) via lentiviral delivery of a doxycycline-inducible NGN2 expression cassette as previously described [57] (Figure S12A). At 2 and 4 weeks post-NGN2 induction, cells expressed neuronal markers TUJ1 and MAP2 and exhibited robust neuronal processes as demonstrated by pan-axonal staining (Figure S12B,C). Both WT and ABCA7 LoF lines were capable of firing action potentials upon current injections (Figure S13A,B). Although the ABCA7 genotype did not alter resting membrane potential (Figure S13E), ABCA7 LoF iNs fired action potentials more readily and at lower current injection thresholds compared to WT iNs (Figure S13F,G), indicating a hyperexcitability phenotype. Collectively, these data confirm successful neuronal differentiation from iPSCs, robust electrophysiological activity, and recapitulation of Alzheimer’s disease-associated neuronal hyperexcitability.

### ABCA7 LoF iNs Recapitulate Excitatory Neuronal Transcriptional Signatures

To investigate whether transcriptional changes associated with ABCA7 LoF observed in *postmortem* human neurons are recapitulated in iNs, we performed bulk mRNA sequencing on ABCA7 WT, p.Glu50fs*3, and p.Tyr622* iNs (N=2, N=5, and N=5, respectively) after four weeks in culture (Data S8). Gene perturbation scores (defined as score = sign(log(FC)) × − log_10_(*p*-value)) showed a strong correlation between p.Glu50fs*3 vs. WT and p.Tyr622* vs. WT comparisons (Pearson correlation coefficient = 0.84; Figure 3B), indicating consistency in the transcriptional impact of ABCA7 variants.

We next conducted gene set enrichment analysis (GSEA) on the differentially expressed genes from these comparisons, identifying 15 significantly perturbed pathways in each comparison, p.Glu50fs*3 vs. WT and Y in p.Tyr622* vs. WT (FDR-adjusted p < 0.05; WikiPathways). These pathways were driven by 356 and 334 unique “leading edge” genes, respectively [52]. K/L partitioning of these leading edge genes identified 9 clusters for p.Tyr622* (Figure 3C) and 10 clusters for p.Glu50fs*3 (Figure S14A). Eight of nine p.Tyr622* T clusters and eight of ten p.Glu50fs*3 G clusters showed significant overlap (FDR-adjusted p < 0.05) (Figure S14B), indicating substantial concordance between the two ABCA7 variant lines.

We also observed that transcriptional signatures in ABCA7 LoF iNs closely aligned with those identified in *postmortem* excitatory neurons. Specifically, we found significant overlap in 5 out of 9 p.Tyr622*-associated clusters (Figure 3D) and in 7 out of 10 p.Glu50fs*3-associated clusters (Figure S14C) with the clusters identified in *postmortem* excitatory neurons, with the majority (4 out of 5 and 6 out of 7, respectively) showing concordant directional changes.

Due to the transcriptional similarity between the two LoF lines, our primary analysis focuses on the patient variant p.Tyr622*, with results for the p.Glu50fs*3 variant provided in supplementary materials (Figure S14). Consistent with findings from *postmortem* data, p.Tyr622* iNs exhibited downregulated clusters associated with lipid metabolism (T.9 and T.13) and upregulated clusters related to cell cycle regulation and proteasomal activity (T.8 and T.14) compared to WT iNs (Figure 3E). Notably, a mitochondrial cluster (T.10) demonstrated the most robust overlap with *postmortem* data (PM.1) and was consistently upregulated in both the p.Tyr622* and p.Glu50fs*3 lines, mirroring the findings in *postmortem* neurons (Figure 3D; Figure S14C). The probability of observing this degree of overlap by chance alone is very low (*p <* 5*x*10^−5^ in both cases, binomial test). Together, these data support a causal relationship between ABCA7 LoF variants and multiple transcriptional signatures observed in *postmortem* excitatory neurons, including mitochondrial, proteasomal, cell cycle, and lipid metabolism components.

### ABCA7 LoF impairs mitochondrial uncoupling in neurons

To further characterize mitochondrial alterations in ABCA7 LoF iNs, extending beyond the gene sets used for K/L cluster analysis, we examined the expression of 1,136 mitochondrial genes curated from the MitoCarta database in our bulk RNAseq data. Among the most significantly upregulated genes in p.Tyr622* versus WT iNs were genes encoding components of mitochondrial apoptosis pathways (e.g., *CASP3*, *BID*) and OXPHOS subunits (previously captured in clusters PM.1 and T.10) (Figure 3F; Table S4). Conversely, downregulated genes were significantly enriched (padj < 0.05) for key metabolic processes, including *β*-oxidation (*ACAD* and *CPT* genes), mitochondrial metabolite transport (*SLC25* genes), and oxidative stress detoxification (*CAT* ) (Figure 3F; Table S4). These MitoCarta mitochondrial gene expression profiles were highly correlated between p.Tyr622* and p.Glu50fs*3 relative to WT iNs (Figure S14E).

To directly assess mitochondrial function in ABCA7 LoF neurons, we measured the oxygen consumption rate (OCR) of WT and ABCA7 LoF iNs over time using the Seahorse metabolic flux assay (Figure S15A,B). The OCR-driven movement of protons across the inner mitochondrial membrane during OXPHOS builds and maintains the mitochondrial membrane potential (ΔΨm)(Figure S15C), and measuring OCR in the presence of mitochondrial inhibitors provides several functional readouts. Because OCR can be influenced by cell viability and mitochondrial abundance [58, 59], we only report internally normalized OCR ratios rather than absolute values [60] for WT, ABCA7 p.Glu50fs*3, and ABCA7 p.Tyr622* iNs. To assess the spare respiratory capacity, we normalized the OCR measured following pharmacological collapse of the proton gradient to the basal OCR, with higher values indicating more spare respiratory capacity [60] (Figure S15D). We then quantified the proportion of basal oxygen consumption that can be attributed to rebuilding the membrane potential lost due to proton leakage through the membrane (i.e., uncoupled mitochondrial OCR) rather than due to ATP synthesis [60] (Figure S15E).

While spare respiratory capacity was comparable between WT and ABCA7 LoF iNs (Figure S15F), ABCA7 LoF iNs showed significantly reduced uncoupled mitochondrial respiration (Figure 3G). Uncoupled mitochondrial oxygen consumption rates in WT iNs (≈ 20%; Figure 3G) align with previously reported values for neurons and other cell types [61–63], indicating that ABCA7 LoF iNs exhibit abnormally low mitochondrial uncoupling. Consistent with this observation, expression levels of UCP2 a member of the mitochondrial uncoupling protein family expressed in the brain [64] were reduced in ABCA7 LoF iNs (Figure S15G).

Because decreased mitochondrial uncoupling often correlates with elevated mitochondrial membrane potential (ΔΨm) [65, 66], we next assessed ΔΨm in NeuN-positive soma using the fixable MitoHealth dye, which accumulates in mitochondria proportionally to membrane potential. We observed significantly increased MitoHealth fluorescence in both p.Tyr622* and p.Glu50fs* iNs compared to WT per NeuN surface (Figure 3H). To further confirm these findings, we measured ΔΨm in soma and neuronal processes using the fluorescent cation tetramethylrhodamine methyl ester (TMRM) in non-quenching mode. TMRM accumulation was higher in p.Tyr622* iNs relative to WT (Figure 3I), and the specificity of this TMRM signal was validated by showing drastically reduced TMRM signal intensity after depolarization of the ΔΨm with the uncoupler FCCP (Figure S15H). Together, these results indicate that ABCA7 LoF iNs exhibit elevated ΔΨm.

Regulated mitochondrial uncoupling serves as a mechanism to control mitochondrial membrane potential and mitigate reactive oxygen species (ROS) generation [65, 67]. To assess whether ABCA7 LoF iNs exhibited elevated ROS levels, we incubated p.Tyr622* iNs with CellROX dye, a fluorescent indicator of oxidative stress. We observed significantly increased CellROX fluorescence in p.Tyr622* iNs compared to WT iNs (Figure 3J), indicating elevated ROS accumulation in ABCA7 LoF iNs. Together, these data suggest that ABCA7 LoF variants decrease mitochondrial uncoupling and increase oxidative stress in neurons.

### ABCA7 LoF induces phosphatidylcholine imbalance in neurons

Since ABCA7 functions as a lipid transporter, we examined the lipidome of WT and ABCA7 LoF iNs using LC-MS (Data S9). Comparing lipidomic profiles between WT and p.Glu50fs*3 iNs revealed significant alterations across multiple lipid classes, including neutral lipids, phospholipids, sphingolipids, and steroids (Figure S16A,B). Among these, triglycerides (TGs)—particularly species enriched in long-chain, predominantly polyunsaturated fatty acids—were frequently altered, showing significant upregulation in p.Glu50fs*3 iNs (Figure S16B,C).

In line with ABCA7’s established preference for phospholipids [26, 68, 69], several phospholipid species also exhibited notable differences (Figure S16B). Phosphatidylcholines (PCs), which are essential structural components of biological membranes and potential ABCA7 substrates [26, 54], were most prominently affected; the majority (≈64% of perturbed PC species) showed increased abundance in p.Glu50fs3 iNs (Figure S16B). Further analysis based on fatty acid saturation—an important factor influencing membrane fluidity—revealed significant enrichment of saturated PCs among the upregulated species (hypergeometric p=0.026) (Figure S16D). In contrast, polyunsaturated fatty acid-containing (PUFA) PCs showed mixed directionality, with several highly unsaturated species showing decreased abundance (*e.g.*, PC(44:7) and PC(38:7)) (Figure S16E,F).

To determine whether neutral lipid and PC imbalances were conserved in p.Tyr622* iNs, we performed targeted lipidomic analysis in positive ionization mode. Consistent with p.Glu50fs*3 iNs, upregulated lipids in p.Tyr622* iNs were significantly enriched for saturated PCs (hypergeometric p=0.044) (Figure 3K; Figure S16G,H). However, PUFA PCs and longchain triglycerides were not reliably detected in this LC-MS run (Figure S16I,J), leaving it unclear whether p.Tyr622* iNs exhibit the same changes in PUFA PC or long-chain triglycerides as p.Glu50fs*3 iNs.

*De novo* PC synthesis occurs via the Kennedy pathway, and subsequent remodeling of the fatty acyl chains is catalyzed by LPCAT enzymes through the Lands cycle, with LPCAT3 specifically introducing PUFA chains into PCs [70–72]. LPCAT3 expression was reduced in p.Tyr622* and p.Glu50fs*3 iNs compared to WT (Figure S16K,L), aligning with increased levels of saturated PCs in these cells. Overall, our data indicate accumulation of neutral lipids in ABCA7 LoF iNs, including long-chain polyunsaturated triglycerides and sterol lipids (zymosteryl), and reveal imbalances in PC composition, with higher saturated species.

### Treatment with CDP-choline reverses impacts of ABCA7 LoF in neurons

Previous work indicated that exogenous choline supplementation normalized phospholipid saturation levels in yeast and ameliorated APOE4-related lipid phenotypes [73, 74]. We therefore next examined whether CDP-choline treatment could similarly mitigate ABCA7 LoF-induced phenotypes in iNs.

Targeted LC-MS analysis confirmed that CDP-choline treatment increased its concentration in the media from undetectable to detectable levels (Figure S17A). Additionally, both CDP and choline specifically accumulated in media conditioned by p.Tyr622* cells after treatment (Figure S17A), indicating extracellular hydrolysis of CDP-choline. While intracellular CDP and CDP-choline could not reliably be detected in this experiment, intracellular choline levels significantly increased after treatment (Figure S17B) and expression levels of choline transporters were significantly upregulated (Figure S17C). This suggests that choline was successfully taken up by p.Tyr622* iNs upon CDP-choline treatment.

We anticipated that higher intracellular choline availability would lead to increased phospholipid synthesis. Lipidomic analysis indeed revealed elevated levels of phospholipids, particularly choline-containing phospholipids (PC and lysophosphatidylcholines (LPC)) and sphingolipids (sphingomyelins (SM)), alongside a reduction in a single TG species, with other neutral lipid species showing a similar downward trend (Figure 4A). Consistent with these changes, we observed increased expression of *PCYT1B*, the enzyme responsible for the rate-limiting step in PC synthesis through the Kennedy pathway (Figure S17C). Additionally, expression of several LPCAT genes, including LPCAT3, was elevated after treatment (Figure S17D), coinciding with observed increases in both saturated and unsaturated PC species (Figure 4A; Figure S17E). These findings suggest that CDP-choline treatment promotes synthesis and remodeling of choline-containing lipids.

**Figure 4:**
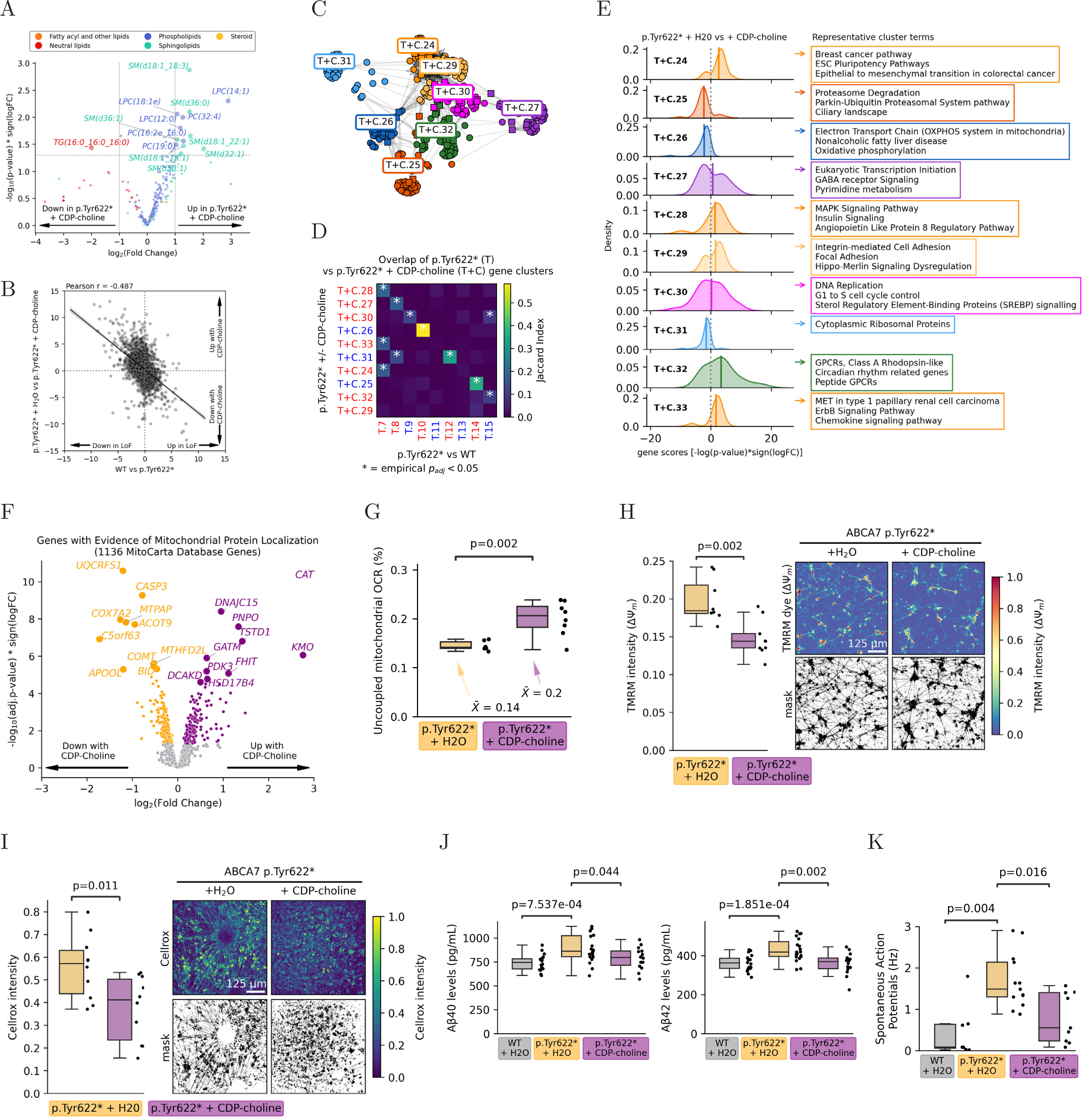
CDP-choline Treatment Rescues ABCA7 LoF-Induced Disruptions in Neurons. **(A)** Volcano plot of differentially abundant lipid species in p.Tyr622* iNs cultured for 4 weeks (treated with or without 100 *µ*M CDP-choline during the final 2 weeks), colored by lipid class. Statistical comparisons by independent-sample *t*-tests. *N* = 5 wells per genotype. **(B)** Correlation of gene perturbation scores (*S* = − log_10_(*p*) × sign(log_2_(FC))) comparing p.Tyr622* vs. WT and p.Tyr622* ± CDP-choline iNs. **(C)** Kernighan-Lin (K/L) clustering of leading-edge genes from significantly perturbed pathways (BH FDR-adjusted *p <* 0.05) comparing p.Tyr622* ± CDP-choline iNs. Colors represent distinct K/L gene clusters, matched to p.Tyr622* vs. WT cluster colors based on Jaccard analysis in (D). **(D)** Heatmap of Jaccard index overlap between K/L clusters from p.Tyr622* vs. WT and p.Tyr622* ± CDP-choline iNs. **(E)** (left) Gaussian kernel density plots of gene perturbation scores (*S*, positive values indicate upregulation with CDP-choline treatment) for each cluster. Solid lines denote cluster means. (right) Representative pathways annotating the most genes per cluster. **(F)** Volcano plot of differential expression of genes with mitochondrial-localized protein products (MitoCarta) for p.Tyr622* ± CDP-choline iNs. **(G)** Mitochondrial uncoupling quantified by Seahorse assay (proportion of basal oxygen consumption due to proton leak) in p.Tyr622* ± CDP-choline iNs cultured for 4 weeks (treated with or without 100 *µ*M CDP-choline during the final 2 weeks). Each datapoint represents OCR from a single well. Statistical comparisons via independent-sample *t*-tests. *N* = 6 (vehicle), 8 (CDP-choline) wells. **(H)** Average TMRM fluorescence intensity per mask (thresholded at 75th percentile) in p.Tyr622* ± CDP-choline iNs cultured for 4 weeks (treated with or without 100 *µ*M CDP-choline during the final 2 weeks), under baseline and FCCP-treated conditions. *N* = 8 wells in each condition. **(I)** Average CellROX fluorescence intensity per mask (thresholded at 75th percentile) in p.Tyr622* ± CDP-choline iNs cultured for 4 weeks (treated with or without 100 *µ*M CDP-choline during the final 2 weeks). Each datapoint represents average intensity per well. *N* = 10 wells in each condition. **(J)** Quantification of secreted A*β* levels from media of cortical organoids derived from WT or p.Tyr622* iPSCs (cultured for 182 days), treated with or without 1 mM CDP-choline for 4 weeks. Each datapoint represents A*β* levels measured for a single cortical organoid. *N* = 20 (WT), 19 (p.Tyr622*), and 14 (p.Tyr622* + 1 mM CDP-choline) organoids. Spontaneous action potentials recorded from dissociated cortical organoids derived from p.Tyr622* iPSCs (cultured for 150 days, followed by 2 weeks treatment post-dissociation), treated with or without 100 *µ*M CDP-choline. Each datapoint represents an individual cell. *N* = 7 (WT), 13 (p.Tyr622*), and 9 (p.Tyr622* + 100 *µ*M CDP-choline) cells.

Next, we characterized changes induced by CDP-choline treatment using LC-MS-based metabolomics and bulk RNAseq. While most of the metabolites increasing or decreasing after treatment could not be annotated, a principal component analysis of the overall metabolite changes indicated that CDP-choline treatment reversed the separation of WT and pTyr622* iN along the axis of the first principal component (PC1; Figure S17F). Transcriptionally, CDP-choline treatment also induced significant changes, clearly distinguishing treated from untreated samples (Figure S17G). The transcriptional signature of CDP-choline treatment negatively correlated with that of p.Tyr622* (Figure 4B), suggesting partial restoration toward the WT state. Performing K/L cluster analysis on the p.Tyr622* vs CDP-choline treated p.Tyr622* samples (Figure 4C), we observed significant overlap in 7 of the 9 clusters identified in the p.Tyr622* vs WT comparison (Figure 4D), with 5 of these clusters showing reversed directional changes following treatment (Figure 4E).

Specifically, clusters related to proteasomal and ribosomal functions (T+C.25 and T+C.31)—previously upregulated in p.Tyr622* (see T.14 and T.12)—were downregulated following CDP-choline treatment (Figure 4D). Most notably, mitochondrial cluster T+C.26—which strongly overlapped with cluster T.10, the cluster most consistent with *postmortem* PM.1—was also reversed after treatment (Figure 4E). Further analysis using the MitoCarta database confirmed a significant reversal in expression of genes encoding mitochondrial proteins, including reduced expression of apoptosis-related genes (*BID*, *CASP3* ; Figure 4F), restoration (upregulation) of the mitochondrial metabolic signature (Table S5), and increased expression of regulators of mitochondrial fusion (*MFN2*, *OPA1* ), a process which enables high metabolic capacity, dissipation of mitochondrial membrane potential, and mitochondrial biogenesis [75]. Overall, ABCA7 LoF-related changes to expression of MitoCarta genes were significantly reversed following CDP-choline treatment (Figure S17H).

To determine whether CDP-choline treatment could restore mitochondrial uncoupling to WT levels, we repeated the Seahorse assay on p.Tyr622* iNs with and without CDPcholine treatment (Figure S17I,J). CDP-choline treatment significantly increased uncoupled respiration in p.Tyr622* iNs, restoring it to WT levels (Figure 4G), with no significant change in spare respiratory capacity (Figure S17K). Consistent with this finding, both TMRM staining (Figure 4H) and MitoHealth fluorescence per NeuN-positive surface (Figure S17L) confirmed a decrease in the mitochondrial membrane potential (ΔΨm) in treated cells. Additionally, CDP-choline treatment significantly decreased CellROX fluorescence (Figure 4I), indicating a reduction in oxidative stress.

### CDP-Choline Ameliorates AD-Associated Phenotypes in Cortical Organoids

Next, we tested whether CDP-choline treatment could improve key AD-associated phenotypes, since previous studies have linked ABCA7 dysfunction to altered amyloid-*β* (A*β*) processing [33–36, 76]. Indeed, p.Tyr622* iNs secreted significantly higher levels of A*β*40 and showed a trending increase in A*β*42 secretion into the media, as measured by enzyme-linked immunosorbent assay (ELISA), although absolute levels remained relatively low (Figure S18A). To study CDP-choline’s effects in a model with stronger pathology, we differentiated p.Tyr622* and WT lines into cortical organoids matured for ≈ 6 months (Figure S18B), a stage at which we observed robust A*β* secretion (approximately two- to four-fold higher levels of A*β*40 and A*β*42 compared to iNs)(Figure S18C). Treatment with 1 mM CDP-choline for four weeks reduced A*β*40 and A*β*42 secretion from p.Tyr622* organoids to WT levels (Figure 4K). This effect was not observed at lower concentrations or shorter treatment durations (Figure S18C). Additionally, treatment of dissociated cortical organoids with 100 *µ*M CDP-choline for two weeks significantly reduced neuronal hyperexcitability in p.Tyr622* organoids, as shown by electrophysiology (Figure 4K).

## Discussion

Loss-of-function (LoF) mutations in the lipid transporter ABCA7 are among the strongest genetic risk factors for late-onset AD. Here, we generated a transcriptional atlas of ABCA7 LoF effects across all major brain cell types in the human prefrontal cortex. Our dataset showed the highest levels of ABCA7 expression in excitatory neurons and strong evidence that ABCA7 LoF led to transcriptional perturbation in pathways related to lipid biosynthesis, mitochondrial respiration, and cellular stress, including up-regulation of DNA damage pathways, and changes to inflammatory and synaptic genes. Using iPSC-derived isogenic neuronal lines (iN) with and without ABCA7 LoF variants, we show that ABCA7 LoF leads to decreased mitochondrial uncoupling, elevated mitochondrial membrane potential, and increased reactive oxygen species (ROS). Consistent with ABCA7’s role as a phospholipid transporter, ABCA7 LoF iNs exhibited significant imbalances in phosphatidylcholine composition, characterized by increased saturated PCs and a decrease to several highly polyunsaturated (PUFA) PCs. Similar changes in phospholipid saturation were recently observed in neuronal models of ALS/FTD, highlighting the broader significance of phospholipid saturation in neurodegenerative conditions [77]. Treatment of ABCA7 LoF iN with CDP-choline increased phosphatidylcholine synthesis, upregulated expression of phosphatidylcholine remodeling enzymes, and rescued mitochondrial uncoupling, mitochondrial membrane potential, and oxidative stress. In addition, CDP-choline supplementation mitigated hyperexcitability and amyloid-*β* secretion in ABCA7 LoF neurons. Together, our data indicate that the observed effects of ABCA7 LoF on neurons may be at least partially mediated by imbalances in phosphatidylcholine metabolism.

While the precise mechanism linking ABCA7 LoF to phosphatidylcholine imbalance remains unclear, disrupted ABCA7 floppase activity—responsible for phospholipid flipping across membrane leaflets—likely impacts membrane fluidity and curvature [78, 79], important determinants of numerous cellular functions [80, 81]. Changes in membrane composition may also broadly affect lipid metabolism by altering the activity of transcriptional regulators controlling lipid biosynthesis and remodeling genes (including LPCATs), which are responsive to shifts in membrane properties [82, 83]. Consistent with our observations and previous reports, CDP-choline supplementation supports *de novo* synthesis of phosphatidylcholine species containing both saturated and polyunsaturated fatty acids [84]. Thus, CDP-choline may help restore phosphatidylcholine balance in ABCA7 LoF neurons by supporting the synthesis and remodeling of diverse phosphatidylcholine species. Given that phosphatidylcholine species are ubiquitous components of biological membranes—including abundant lipids within mitochondrial membranes [85]—imbalances in their fatty acyl chain composition could broadly impact cellular functions [71, 86], including mitochondrial activity. Indeed, alterations in phospholipid saturation and composition have been linked to changes in mitochondrial dynamics, cristae morphology, bioenergetics, and membrane potential [85, 87]. However, additional studies are needed to clarify the precise mechanisms by which phosphatidylcholine imbalances influence mitochondrial function and uncoupling dynamics.

Mitochondrial dysfunction, including impaired mitochondrial uncoupling, is increasingly recognized as critical in aging, AD, and other neurodegenerative diseases. Although mitochondrial uncoupling was recently linked to frontotemporal dementia, its specific role in AD remains poorly investigated [88–91]. Neurons maintain high mitochondrial OXPHOS to meet their significant energy demands [92, 93]. Mitochondrial uncoupling, actively regulated by mitochondrial proteins [94], supports mitochondrial health by modulating mitochondrial membrane potential, reducing reactive oxygen species [90, 95], and promoting mitochondrial biogenesis [96–98]. Impaired mitochondrial uncoupling, as observed in ABCA7 LoF neurons, can elevate oxidative stress, impair synaptic and calcium signaling, and contribute to neurodegeneration [96–98]. Increased oxidative stress also triggers DNA damage and inflammatory responses [99–102], which are elevated in the presence of ABCA7 LoF based on transcriptomic signatures.

In line with our findings linking phosphatidylcholine imbalances to mitochondrial impairments in ABCA7 LoF neurons, a recent study in ABCA7 deficient neurospheroids independently revealed a link between phosphatidylglycerol deficiency and mitochondrial function [30], further highlighting the importance of lipid-centric therapeutic interventions for ABCA7 LoF. Here, we offer a therapeutic strategy to reverse these dysfunctions including ABCA7 LoF-induced AD pathology and neuronal hyperexcitability -through CDP-choline treatment, a readily available and safe dietary supplement [103–105]. Recent work from our lab implicates phosphatidylcholine and fatty acyl saturation imbalances in APOE4 dysfunction [74], and in cognitive resilience to AD pathology [106], suggesting that phosphatidylcholine disruptions may be central to AD risk in large fractions of the population. Indeed, our work suggests that the common missense variant p.Ala1527Gly likely has convergent effects with ABCA7 LoF. Genetic interactions with other risk factors, including APOE4, may exacerbate otherwise subtle ABCA7 dysfunction, and contribute to risk in a significant subset of AD cases [107–110]. As such, our study supports a growing body of literature, including recent studies on APOE4 [111, 112] that implicates lipid disruptions in the etiology of AD, and pinpoints additional genotypes that may benefit from interventions on phosphatidylcholine metabolism.

## Acknowledgments

We thank the individuals who donated postmortem brain samples, and their families, for enabling this research; Y. Zhou, E. McNamara and T. Garvey for administrative support and animal care; U. Geigenmüller for reviewing and editing the manuscript and for helpful discussions; Rebecca Pinals for helpful discussions and input on the manuscript; J. Davila-Velderrain and R. Firenze for providing input on the manuscript; the MIT SuperCloud and Lincoln Laboratory Supercomputing Center for providing HPC and consultation resources that have contributed to the research results reported within this paper; the Harvard Center for Mass Spectrometry (HCMS) for lipidomic and metabolomic sample runs and Charles Vidoudez (HCMS) for consultation on the lipidomic data analysis; the MIT BioMicro Center for bulk and single-nucleus RNA-sequencing runs and Stuart Levine and his Team for consultations on data processing; ROSMAP resources can be requested at https://www.radc.rush.edu. Graphic illustrations were generated using BioRender under agreement # X25SQ61W7.

## Author contributions

DVM and L-HT designed the study, with L-HT supervising the overall project and acquiring funding alongside MK. Experimental work included snRNA-seq experiments performed by DVM, JMB, HR, and LL; Seahorse assays conducted by SEW; and differentiation and maintenance of iPSC lines, induced neurons, and cortical organoids by SEW, CS, P-CP, and OK. SEW and P-CP prepared samples for LC-MS analysis and performed amyloid ELISA experiments and neuronal marker staining. P-CP conducted TMRM and CellRox assays and SEW performed MitoHealth assays. P-CP, CS, and OK prepared bulk RNA samples. LL conducted electrophysiological recordings and analysis, and CJY generated the p.Tyr622* cell line. AS performed molecular dynamic simulations, analyses, and visualizations and DVM performed formal data analysis and visualization, with contributions from C-AB, GL, and AS. Experimental and technical support was provided by A-NS, ML, G-SM, GW, AG, NL, and GS. C-CC and DL helped with revision experiments. DVM, SEW, and L-HT wrote the first draft of the manuscript. DVM and L-HT wrote the revised draft of the manuscript.

## Ethics Statement

The study protocol involving the use of human stem cells was approved by the Coriell Institutional Review Board (Coriell IRB) in compliance with DHHS regulations (45 CFR Part 46). The initial cell lines were obtained from the Coriell Institute, which ensured that informed consent was received from all donors. Donors were informed that their tissue donations would be used for the creation of cell lines intended for educational and research purposes, and that all biological materials would be anonymized. More information can be found here. For postmortem human brain samples, informed consent was obtained from each subject, and the Religious Orders Study and Rush Memory and Aging Project were approved by an Institutional Review Board (IRB) of Rush University Medical Center.

## Data availability

All postmortem human data can be accessed through the Synapse AD Knowledge Portal (syn53461705), which also includes associated ROSMAP metadata. These data are subject to controlled access in compliance with human privacy regulations. To obtain the data, a data use agreement (DUA) must be completed. This requirement ensures the anonymity of ROSMAP study participants. A DUA can be established with either the Rush University Medical Center (RUMC) or SAGE, the organization that manages Synapse. The necessary forms are available for download on their respective websites. All iPSC-related data are accessible through links provided in our code repositories. For a complete list of data availability and download links, please refer to the code repositories listed below. Additionally, relevant processed datasets are available in the supplementary files of this manuscript.

## Code availability

All code used in this study is available on GitHub (https://github.com/djunamay/ABCA7lof2).

## Supplementary Materials

### Materials and Methods

#### Experimental Methods on human *postmortem* brain tissue

**Isolation of nuclei from frozen postmortem brain tissue.** For batch #1: The protocol for the isolation of nuclei from frozen postmortem brain tissue (region BA10) was adapted for smaller sample volumes from a previous study [50]. All procedures were carried out on ice or at 4°C. In brief, postmortem brain tissue was homogenized in 700 µl Homogenization Buffer (320 mM sucrose, 5 mM CaCl2, 3 mM Mg(CH3COO)2, 10 mM Tris HCl pH 7.8, 0.1 mM EDTA pH 8.0, 0.1% IGEPAL CA-630, 1 mM *β*-mercaptoethanol, and 0.4 U µl-1 recombinant RNase inhibitor (Clontech)) using a Wheaton Dounce tissue grinder (15 strokes with the loose pestle). Homogenized tissue was filtered through a 40-µm cell strainer, mixed with an equal volume of Working Solution, which is prepared by mixing Diluent (30mM CaCl2, 18mM Mg(CH3COO)2, 60mM Tris pH 7.8, 0.6mM EDTA, 6mM *β*-mercaptoethanol) with Optiprep density gradient solution (Sigma-Aldrich D1556-250ML) in a 1:5 ratio. The sample mix was then loaded on top of an Optiprep density gradient consisting of 750 µl 30% OptiPrep solution (1.5:1 ratio of Working Solution:Homogenization Buffer) on top of 300 µl 40% OptiPrep solution (4:1 ratio of Working Solution:Homogenization Buffer). The nuclei were separated by centrifugation (5 min, 10,000 g, 4 °C). Approximately 100µl of nuclei were collected from the 30%/40% interface and washed twice with 1 ml of PBS containing 0.04% BSA, centrifuging 300g for 3 min (4 °C) in between, then resuspended in 100µl PBS containing 0.04% BSA. The nuclei were counted on C-Chip disposable hemocytometer and diluted to 1000 nuclei per µl in PBS containing 0.04% BSA.

For batch #2: These samples (fresh postmortem brain; PFC BA10) were prepared as part of and according to a previous study [50].

Informed consent and Anatomical Gift Act consent were obtained from each participant. The Religious Orders Study and Rush Memory and Aging Project were approved by the Institutional Review Board (IRB) of Rush University Medical Center. All participants signed a repository consent, allowing their data and biospecimens to be shared.

**Droplet-based snRNA-seq.** For batch #1: cDNA libraries were generated using the Chromium Single Cell 3 Reagent Kits v3 following the manufacturer’s protocol (10x Genomics). Libraries were sequenced on the NovaSeq 6000 S2 platform (paired-end, 28 + 91 bp, with an 8-nucleotide index). Samples were distributed across two lanes and sequenced twice on separate flow cells to enhance sequencing depth.

For batch #2: Libraries were prepared using Chromium Single Cell 3 Reagent Kits v2 and sequenced with the NextSeq 500/550 High Output v2 kits (150 cycles), as described in our previously published study [50].

Raw sequencing reads from all samples were processed jointly for alignment and gene counting.

#### Culture and generation of human isogenic iPSCs

A control parental line was derived from a 75-year-old female (AG09173) with an APOE3/3 genotype by the Picower Institute for Learning and Memory iPSC Facility as first described [113]. Two ABCA7 LoF isogenic lines were derived from parental AG09173. ABCA7 p.Glu50fs*3, generated by Synthego (www.synthego.com), contains a premature termination codon in exon 3 (Figure S11A), which to our knowledge has not been discovered in patients, but is functionally analogous to patient loss-of-function mutations.

ABCA7 p.Tyr622* contains a patient-derived mutation (Y622*) [114] and was generated in house by CRISPR-Cas9 genome editing. The CRISPR/Cas9-ABCA7Y622* sgRNA plasmid was prepared followed by the published protocol [115]. In brief, a sgRNA sequence within 10 nucleotides from the target site was designed using the CRISPR/Cas9 Design Tool (http://crispr.mit.edu). The oligomer pairs (forward: 5’-CACCGCCCCTACAGCCACCCGGGCG-3’ and reverse: 5’AAACCGCCCGGGTGGCTGTAGGGGC-3’) were annealed and cloned into pSpCas9-2A-GFP (PX458) plasmid (Addgene #48138). Plasmid DNA was submitted for Sanger sequencing to confirm correct ABCA7 sgRNA sequence (Figure S11B).

AG09173 iPSCs were dissociated with Accutase (Thermo Fisher Scientific) supplemented with 10 M ROCK inhibitor (Tocris) for electroporation using Amaxa and Human Stem Cell Nucleofector Kit I (Lonza). 5x106 cells were resuspended in 100 l of reaction buffer supplemented with 7.5 g of CRISPR/Cas9-ABCA7 sgRNA plasmid and 15 g of single-strand oligodeoxynucleotide (ssODN) template (5’-GGTGCGCGCCCCCAGGCCAATCCAGGAGCTGCACCCTAAGCTCCCGTTGCCTCTCACAGCTGGGAGACATCCTCCCCTAGAGCCACCCGGGCGTCGTCTTCCTGTTCTTGGCAGCCTTCGCGGTGGCCACGGTGACCCAGAGCTTCCTGCTCAGCGCCTTCTTCTCCCGCGCCAACCTGG-3’). This reaction mixture was nucleofected with program A-23, resuspended with media supplemented with 10 M ROCK inhibitor and seeded on MEF plates. Two days after electroporation, cells were dissociated and filtered through Falcon polystyrene test tubes (Corning #352235), transferred to Falcon polypropylene test tubes (Corning #352063) and sorted by BD FACS Aria IIU in FACS Facility at the Whitehead Institute and seeded as single cells in media supplemented with 1X Penicillin-Streptomycin (P/S, Gemini Bio-products) and 10 M ROCK inhibitor. After sufficient colony growth, each colony was transferred in part to a 12-well plate while the remainder was collected and used to extract genomic DNA (Qiagen DNeasy Blood Tissue Kit, Cat. No. 69504) and screen for the Y622* mutation by sanger sequencing.

All lines used were confirmed to have normal karyotypes before use and periodically reviewed (Cell Line Genetics) (Figure S10C). All human iPSCs were maintained at 37°C and 5% CO2, in feeder-free conditions in mTeSR-1 medium (Cat #85850; STEMCELL Technologies) on Matrigel-coated plates (Cat # 354277; Corning; hESC-Qualified Matrix). iPSCs were passaged at 60–80% confluence using ReLeSR (Cat# 05872; STEMCELL Technologies) and reseeded between 1:6 and 1:24 (depending on desired density) onto Matrigel-coated plates.

#### Experimental Methods on iNs

**rTTA and NGN2 Virus production.** HEK293T cells (ATCC, Cat#CRL-3216) were maintained in DMEM/F-12, GlutaMAX (ThermoFisher, Cat#10565018), 10% fetal bovine serum (GeminiBio, SKU#100-106), 1% MEM Non-essential amino acids (Sigma, Cat#M7145), 1% sodium pyruvate (ThermoFisher, Cat#11360070), and 1% Penicillin-Streptomycin (GeminiBio, SKU#400-109). Cells were passaged for maintenance with TrypLE (ThermoFisher, Cat#12605010) at 70-80% confluence and reseeded 1:10 in 10 cm tissue culture plates.

For transfection, HEK293T cells were seeded at 5x106 cells per 10 cm plate. Transfection mixtures containing the components required for 3rd generation lentiviral production (per 10 cm dish: 10 µg EF1a-rtTA-Hygro (Addgene #66810) or pLV-TetO-hNGN2-eGFP-Puro (Addgene #79823), 5 µg pMDL g/pRRE, 2.5 µg pRSV-Rev, 2.5 µg MD2.G, and 48 µL polyethyleneimine (1 mg/mL) in 600 uL OptiMEM (Fisher, Cat#51-985-034)). Mixtures were inverted 10X and incubated at RT for 20 min, then added dropwise to the dish. Transfection media was removed 16h later and replaced with 10 mL fresh media. Three days after transfection, media was collected and centrifuged at 3000 xg for 5 min at 4°C to pellet any contaminating cells. Supernatant was transferred to sterile Millex glass ultracentrifuge tubes and centrifuged at 25,000 rpm for 2 hours using a SW32Ti rotor in a Beckman Optima L-90K Ultracentrifuge. The pellets were resuspended in 1 mL PBS per 10 cm plate, and stored at -80°C until use.

**Lentivirus-mediated NGN2 induction in iPSCs and drug treatments.** iPSCs were dissociated into single cell suspension with Cell Dissociation Buffer (Life Technologies, Cat#13151-014), centrifuged at 300 xg for 5 min, and resuspended in mTeSR1 media with Rock inhibitor (Rockout; Abcam, ab285418). Single-cell suspension was plated in a 6-well plate coated with Matrigel for an optimized seeding density of 50-60% confluence 24 hours after plating. One day after plating, cells were co-transduced with 80 µL pLV-TetO-hNGN2eGFP-Puro and 80 µL EF1a-rtTA-Hygro added in 1 mL fresh media per well and incubated overnight at 37°C. NGN2 expression was then induced 24 hours later with addition of 2 mL fresh media supplemented with doxycycline (DOX, 1 µg/mL, final concentration) and Rock inhibitor. Puromycin (1 µg/mL) selection of non-NGN2 expressing cells was performed with media change 24 hours after induction, with continued DOX supplementation. After 24 hours of puromycin selection, immature neurons were re-plated onto PDL/laminin coated plates at 1x106 cells/well on 6-well plates or 5x104 cells/well on 96-well plates. Neurons were maintained in BrainPhys Neuronal Media (STEMCELL Technologies, Cat#05793) with Neurocult SM1 Neuronal Supplement (STEMCELL Technologies, Cat#05711), N2supplement-A (STEMCELL Technologies, Cat#07152), laminin (1ug/mL), and DOX (1 µg/mL) with half media changes every 3-4 days. Neuronal cultures were maintained for 28 days before experimentation.

iPSC-derived neurons were treated with cytidine 5’-diphosphocholine (CDP-choline; Millipore Sigma 30290) to final concentrations of 100 µM beginning at day 14 and repeated with each media change until 28 days matured. Choice of treatment concentration and duration was based on a previous study by our lab [74].

**Electrophysiological recordings.** Cells were placed in a recording chamber and perfused with oxygenated artificial cerebrospinal fluid (ACSF) contains (in mM) 125 NaCl, 2.5 KCl, 1.2 NaH2PO4•H2O, 2.4 CaCl2•2H2O, 1.2 MgCl2•6H2O, 26 NaHCO3 and 11 D-Glucose at a constant rate of 2 mL/min at 32°C. Cells were visualized using infrared differential interference contrast (IR-DIC) imaging on an Olympus BX-50WI microscope.

Recordings were performed using Axon Multiclamp 700B and Clampex 11.2 (Molecular Devices). Action potentials were generated by injecting various steps of currents using current clamp configuration. Whole-cell currents were recorded from a holding potential of -80 mV by stepping to various voltages using voltage clamp configuration. Cell-attached recording configution was used to measure spontaneous firing activity. Signals were filtered at 1 kHz using the amplifier’s four-pole, low-pass Bessel filter, digitized at 10 kHz with a Digidata 1550B interface (Molecular Devices). Pipette solution contained (in mM) 120 K gluconate, 5 KCl, 2 MgCl2•6H2O, 10 HEPES, 4 ATP, 0.2 GTP. pClamp 11.2 (Molecular Devices) and GraphPad Prism 10 software suites were used for data acquisition and analysis. Data are presented as means ± standard errors of means (SEM).

**A***β* **Enzyme linked immunosorbent assays on iNs.** Media was collected from 4 week old iNs and flash frozen. ELISAs were performed on thawed media according to manufacturer’s instructions to measure A40 (ThermoFisher Scientific, KHB3481) and A42 (ThermoFisher Scientific, KHB3441) respectively.

**Mitochondrial Health cell stain.** HCS Mitochondrial Health kit (ThermoFisher, Cat#H10295) were used on live cells according to manufacturer’s protocols. In brief, 50 uL of media containing 1.5 µL MitoHealth dye was added to each well and incubated on live cells for 30 min at 37°C. Next, neurons were fixed in 4% paraformaldehyde/4% sucrose in PBS at 4°C for 15 min at room temperature, washed 3X with PBS, then permeabilized with 0.1% Triton-X in PBS for 5 min at room temperature. Cells were blocked in 2% Bovine Serum Albumin (BSA, Fisher Bioreagents, BP9703) in PBS for 20 min at room temperature, then incubated in primary antibodies diluted in blocking solution (NeuN; 1:500) overnight at 4°C. Cells were washed 3X for 5 min with PBS, then incubated in secondary antibodies diluted in blocking solution (1:1000) for 2 hours at room temperature. Cells were washed 3X for 5 min with PBS, then incubated for 10 min with 1:2000 Hoechst 33342 (Invitrogen, H3570). Cells were washed 1X with PBS, and wells flooded with PBS for imaging.

Confocal images were acquired on a Zeiss LSM900. Acquisition settings were kept constant within each imaging batch (where conditions of interest were uniformly distributed across plates). The minimum and maximum z-plane was manually determined for each culture well, to accommodate differences in culture thickness. Cultures were imaged at 1 µm intervals along the z-axis.

**Live imaging of TMRM.** Stock solutions were initially prepared at concentrations of 10 µM TMRM (ThermoFisher, I34361), 100 µM FCCP (Cayman Chemical, 15218), and 100 µM oligomycin from Streptomyces diastatochromogenes (Sigma, O4876-25MG), achieving final working concentrations of 0.1 µM TMRM, 1 µM FCCP, and 1 µM oligomycin. For imaging, 3 µL of the 10 µM TMRM stock solution was added to each well of iNs cultured in 300 µL of media on a 96-well plate. Cells were incubated with TMRM dye for 30 minutes at 37°C. Live-cell imaging was then performed using a Zeiss LSM 900 confocal microscope equipped with ZEN software. After initial TMRM images were acquired, 3 µL of the 100 µM FCCP stock was added to each well, and images were captured immediately. Subsequently, 3 µL of the 100 µM oligomycin stock was added, followed by immediate image acquisition. Confocal images were acquired as single optical sections (single-plane imaging) on a Zeiss LSM900 microscope.

**Live imaging of CellROX.** iNs were maintained in 300 µL of media per well in a 96-well plate. A stock solution of CellROX Orange Reagent (ThermoFisher, C10443) was prepared at a concentration of 500 µM. For staining, 3 µL of the 500 µM CellROX dye was added directly to each well containing iNs in 300 µL media. Cells were incubated with the dye for 30 minutes at 37°C. After incubation, live-cell images were acquired as single optical sections (single-plane imaging) on a Zeiss LSM900 microscope.

**Seahorse Metabolic Assays.** iPSCs were differentiated as described above directly on 96-well Agilent Seahorse XFe96/XF Pro cell culture microplates and matured for 28 days before assaying on a Seahorse XFe96 Analyzer. Seahorse XF Cell Mito Stress Test and Oxidation Stress Tests were performed according to manufacturer protocol with the following final drug concentration: Oligomycin, 2.5 µM; FCCP, 1 µM; Rotenone/Antimycin, 0.5 µM.

**MAP2 and NeuN staining of iNs.** iNs were plated on coverslips with coating. Cells were fixed in 4% formaldehyde for 10min at room temperature followed by PBS wash once. Cells were next permeabilized and blocked in PBS containing 0.2% TritonX-100 and 10% bovine serum albumin for 1 hour at room temperature, then incubated with primary antibody (MAP2 1:1000; NeuN 1:1000; diluted in blocking solution) at 4 °C overnight. Primary antibody was visualized using the appropriate secondary antibody conjugated to Alexa Fluor 594, or Alexa Fluor 647 (1:500, Thermo Fisher Scientific). Nuclei were visualized with Hoechst 33342 (1:1000, Thermo Fisher Scientific). Coverslips were then mounted on the glass slides with fluoromount g. Confocal images were acquired as single optical sections (single-plane imaging) on a Zeiss LSM900 microscope.

**RNA Extraction, Library Preparation, and Sequencing.** Total RNA was extracted from iNs using the RNeasy Mini Kit (Qiagen), and RNA quality was assessed using a 5300 Fragment Analyzer (Agilent). Only samples with an RNA Quality Number (RQN) greater than 9.5 were selected for library preparation. Full-length cDNA was synthesized using the SMART-seq v4 kit (Takara Bio), and sequencing-ready libraries were subsequently prepared using the Nextera XT DNA Library Preparation Kit (Illumina). Libraries were sequenced as 75 bp + 75 bp paired-end reads with dual 8-nucleotide indexes on an Element AVITI sequencing platform (Element Biosciences) at the MIT BioMicro Center.

#### LC-MS Experiments on iNs

**Biphasic Extraction.** iPSC-derived neurons were washed once with cold PBS (Fisher; Cat#MT21040CM) and lifted off plate with a cell scraper in 1 mL cold PBS. Cells were centrifuged at 2000 xg for 5 min. PBS was removed, and cells were resuspended in 2 mL cold methanol for biphasic extraction. Chloroform (Sigma 1.02444) (4 mL; cold) was added to each vial, and mixed by vortexing for 1 min. Water (Sigma WX0001) (2 mL; cold) was added to each vial, and mixed by vortexing for 1 min. Vials were placed in 50 mL conical tubes and centrifuged for 10 min at 3000 rcf for phase separation. The lower, chloroform phase was collected (3 mL from each sample) and transferred to new vials. In instances where samples were prepared by the Harvard Center for Mass Spectrometry, cell pellets were provided in 500 µL of methanol, vortexed, and transferred to 8 mL glass vials. Each sample received an additional 1.5 mL of methanol and 4 mL of chloroform, followed by vortexing and incubation for 10 min in an ultrasound bath. Next, 2 mL of water was added, and samples were again vortexed. Phase separation was achieved by centrifugation at 800 rcf for 10 min at 4°C. The resulting upper aqueous phases were transferred into new glass vials designated for metabolomics analysis, while the lower chloroform phases were transferred separately for lipidomics analysis. At least one blank sample (containing no cells) was prepared alongside each biphasic extraction experiment and processed identically through the LC-MS analysis pipeline.

**Cell pellet sample preparation for LC-MS Lipidomics.** Subsequent sample preparation for lipidomics was performed by the Harvard Center for Mass Spectrometry. Samples were dried under nitrogen flow until approximately 1 mL remained, transferred into microcentrifuge tubes, and completely evaporated to dryness. Dried samples were resuspended in chloroform, with volumes scaled according to biomass (cell count) using a minimum of 60 µL, then split into two equal aliquots for positive and negative ionization mode analyses. For experiments using only positive ionization mode, samples were resuspended in a smaller biomass-scaled volume (minimum 20–25 µL) without splitting. Following resuspension, samples were centrifuged (maximum speed for 10 min or 18,000 rcf for 20 min at 4°C), and supernatants were transferred into microinserts for LC-MS analysis.

**Cell pellet sample preparation for LC-MS Metabolomics.** Subsequent sample preparation for metabolomics was performed by the Harvard Center for Mass Spectrometry. Samples were dried under nitrogen flow until approximately 1 mL remained, transferred into microcentrifuge tubes, and evaporated completely to dryness. Dried samples were resuspended in 50% acetonitrile in water, using volumes scaled according to provided biomass (minimum 20 µL). Following centrifugation at maximum speed for 10 min, a consistent volume (either 12 µL or 15 µL, depending on the batch) of supernatant from each sample was transferred into microinserts. The remaining supernatants from each batch were pooled separately to create batch-specific quality control (QC) samples.

**Media preparation for LC-MS Metabolomics.** Media samples (100 µL each) were transferred into microcentrifuge tubes containing 1 mL of methanol and incubated at -20°C for 2 hours. Following incubation, samples were centrifuged at 18,000 rcf for 20 min at -9°C, and supernatants were transferred into new tubes and evaporated to dryness under nitrogen flow. The dried samples were resuspended in 50 µL of 30% acetonitrile in water containing 2 mM medronic acid, centrifuged again at 18,000 rcf for 20 min at 4°C, and the resulting supernatants were transferred into glass microinserts for LC-MS analysis.

**LC-MS Lipidomics.** LC-MS lipidomics was performed by the Harvard Center for Mass Spectrometry. LC-MS analyses were modified from [116] and were performed on an Orbitrap Exactive plus MS (Thermo Scientific) in line with an Ultimate 3000 LC (Thermo Scientific). Each sample was analyzed in positive and negative modes, in top 5 automatic data-dependent MS/MS mode. Column hardware consisted of a Biobond C4 column (4.6 x 50 mm, 5 m, Dikma Technologies). Flow rate was set to 100 µL min-1 for 5 min with 0% mobile phase B (MB), then switched to 400 µL min-1 for 50 min, with a linear gradient of MB from 20% to 100%. The column was then washed at 500 µL min-1 for 8 min at 100% MB before being re-equilibrated for 7 min at 0% MB and 500 µL min-1. For positive mode runs, buffers consisted for mobile phase A (MA) of 5mM ammonium formate, 0.1 % formic acid and 5% methanol in water, and for MB of 5 mM ammonium formate, 0.1% formic acid, 5% water, 35% methanol in isopropanol. For negative runs, buffers consisted for MA of 0.03% ammonium hydroxide, 5% methanol in water, and for MB of 0.03% ammonium hydroxide, 5% water, 35% methanol in isopropanol. Lipids were identified and their signal integrated using the Lipidsearch © software (version 4.2.27, Mitsui Knowledge Industry, University of Tokyo). Integrations and peak quality were curated manually before exporting.

**LC-MS Metabolomics.** LC-MS metabolomics was performed by the Harvard Center for Mass Spectrometry. Samples were analyzed by LC-MS on a Vanquish LC coupled to an ID-X MS (Thermofisher Scientific). Five µL of sample was injected on a ZIC-pHILIC peek-coated column (150 mm x 2.1 mm, 5 micron particles, maintained at 40 °C, SigmaAldrich). Buffer A was 20 mM Ammonium Carbonate, 0.1% Ammonium hydroxide in water and Buffer B was Acetonitrile 97% in water. The LC program was as follows: starting at 93% B, to 40% B in 19 min, then to 0% B in 9 min, maintained at 0% B for 5 min, then back to 93% B in 3 min and re-equilibrated at 93% B for 9 min. The flow rate was maintained at 0.15 mL min-1, except for the first 30 seconds where the flow rate was uniformly ramped from 0.05 to 0.15 mL min-1. Data was acquired on the ID-X in switching polarities at 120000 resolution, with an AGC target of 1e5, and a m/z range of 65 to 1000. MS1 data is acquired in switching polarities for all samples. MS2 and MS3 data was acquired on the pool sample using the AquirX DeepScan function, with 5 reinjections, separately in positive and negative ion mode. A mixture for standards of interest was prepared and analyzed immediately following the sample runs for targeted metabolite analysis.

### Experimental Methods on Dorsal Cortical Organoids

**Cortical organoid generation** Dorsal cortical organoid were generated as previously described [117]. In brief, iPSC were cultured until 80-90% confluence, before dissociation and preparing a single cell suspension at 1*x*10^5^ cells/mL in mTesr supplemented with 10 M Rock inhibitor. cortical organoids were then generated by distributing in 100uL / well in PrimeSurface® 96 Slit-well Plates (S-Bio, MS9096SZ). After 48-72 hours, differentiation was induced on days 0 - 5 via daily media change in Neural Induction Media comprised of DMEMf/12 (Life Technologies, cat. no. 11330-032), KnockOut serum replacement (Life Technologies, 10828-028), GlutaMAX, 2-Mercaptoethanol, Penicillin-Streptomycin, and SMAD inhibitors SB-431542 and dorsomorphin. From days 6-16, media was switched to Neural Differentiation medium composed of Neurobasal A medium, B27 supplement, GlutaMAX, Penicillin–streptomycin, Human recombinant EGF (20 ng/ml), Human recombinant FGF2 (20 ng/ml). Daily media change was performed until day 16, followed by every other day until day 25. From day 25 onwards, EGF and FGF was removed and replaced with 20ng/mL Human recombinant brain-derived neurotrophic factor (BDNF; 450-02) and 20 ng/mL Human recombinant neurotrophin 3 (NT3; PeproTech, cat. no. 450-03). After day 45, media was changed two times per week.

**A Enzyme linked immunosorbent assays on cortical organoids.** Culture media was collected from cortical organoids at days 176-182, following 3-4 week treatments with 500 *µ*M or 1 mM CDP-choline. Samples were kept on ice and immediately analyzed using ELISA to quantify levels of A*β*40 and A*β*42. ELISAs were performed according to the manufacturer’s protocols for A*β*40 (ThermoFisher Scientific, KHB3481) and A*β*42 (ThermoFisher Scientific, KHB3441). For the A*β*42 assay, 50 µL of undiluted media was used, whereas media samples were diluted (1:6.67 or 1:10) for the A*β*40 assay. The cortical organoid age (5-6 months) was selected based on empirical evidence showing the emergence of a robust amyloid phenotype at this developmental stage. CDP-choline concentrations were increased from 100 µM to 500 µM and 1 mM to compensate for reduced perfusion in cortical organoids compared to 2D iN cultures.

**Electrophysiological recordings on cortical organoids** To produce 2D cultures from cortical organoids, day 150 cortical organoids were washed in PBS, before transferring to 1.5 mL eppendorf tubes containing Accutase (Stem Cell Technologies, 07920) and placing in a 37C water bath for 40 minutes, with gentle agitation using a P1000 pipette every 5-10 minutes. Dissociated organoids were then plated on #1 glass coverslips (Fisher Scientific, 50-194-4702) with PDL, laminin, and matrigel coating as previously described. 2D cultures were maintained with or without 100 *µ*M CDP-choline treatment for two weeks before electrophysiological recordings. Electrophysiological recordings were performed as described for the iNs. This treatment duration and concentration were selected to align with conditions in 2D iNs experiments. The cortical organoid age was chosen based on empirical observations indicating the emergence of a robust amyloid phenotype at the developmental stage of 5-6 months.

Outliers in spontaneous action potentials were detected using the interquartile range (IQR) method. Specifically, data points falling below Q1 - 2 × IQR or above Q3 + 2 × IQR were identified as outliers. This method resulted in the removal of two data points exhibiting unusually high spontaneous action potentials: one in condition p.Tyr622* (value = 9.38) and one in condition p.Tyr622*+Choline (value = 6.15). Additionally, cells recording zero spontaneous potentials were excluded, as these were presumed to represent glial cells rather than neuronal cells.

**Immunostaining of cortical organoids.** 20 *µ*m cortical organoid sections were fixed in 4% formaldehyde for 10min at room temperature, and cortical organoids were then transferred to 30% sucrose/PBS for dehydration at 4C for 2 to 3 days. cortical organoids were next embedded in optimal cutting temperature compound (OCT compound) and sliced as 20-mm sections using a Cryostat microtome (Leica). For immunostaining, sections were washed with PBS once, followed by permeabilized and blocked in PBS containing 0.2% TritonX-100 and 10% bovine serum albumin for 1 hour at room temperature, then incubated with primary antibody (MAP2 1:1000; NeuN 1:1000, diluted in blocking solution) at 4 °C overnight. Primary antibody was visualized using the appropriate secondary antibody conjugated to Alexa Fluor 488, or Alexa Fluor 594 (1:500, Thermo Fisher Scientific). Nuclei were visualized with Hoechst 33342 (1:1000, Thermo Fisher Scientific). All images were captured using a Zeiss LSM 900 confocal microscope and the ZEN software.

**Image Processing**

**Quantification of fixed z-stack imaging.** Image data were acquired at 8 or 16 bits, with voxel sizes of 1 µm x 0.62 µm x 0.62 µm. Image files extracted from the confocal Zeiss microscope (.czi format) were loaded into Python using the ‘aicsimageio‘ package and converted to floating-point format in the range [0, 1]. Confocal image acquisition settings were kept consistent within each imaging batch.

A pre-trained model (”cyto2” from Cellpose [118]) was applied to segment NeuN+ cell bodies per image. For images sampled at 0.62 µm along the xy-plane, segmentation on the NeuN channel in 3D produced the best results. For images sampled at 0.31 µm along the xy-plane, segmentation on the NeuN and Hoechst channels in 2D (xy), with subsequent stitching along the z-axis, produced the best results. Specific segmentation settings were determined for each imaging experiment. Segmentation quality was assessed manually, blinded to condition, and images with low-quality segmentations were discarded.

The model outputs per-voxel probabilities representing the Bernoulli probability that a given voxel lies within a cell (any cell) and per-voxel masks—recovered from flow vectors and from the pixel probabilities output by the model—representing regions of interest (cells). We leveraged these per-voxel probabilities to compute the expected fluorescence intensity *E*(*I_t_*) for our target channel *t* in each cell *c*. This is calculated as *E*(*I_t_*) = *a* · *b*, where *a* is a 1-dimensional vector containing the measured intensities for channel *t* across all *n* voxels annotated as part of the region of interest for cell *c*, and *b* is a 1-dimensional vector of the same length *n*. Each element *b_i_* in *b* represents the normalized probability that the corresponding voxel *i* belongs to cell *c*, calculated as:

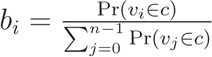

This normalization ensures that the probabilities sum to 1, providing a weighted contribution of each voxel to the total expected fluorescence intensity for the cell.

A linear mixed-effects model was fit (using ‘mixedlm()‘ from the ‘statsmodels‘ package) to cell-level average fluorescence intensities, with treatment or genotype as a fixed effect and well-of-origin as a random effect; formalized as follows:

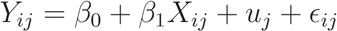

where: *Y_ij_*is the observed fluorescence intensity for cell *i* in well *j*, *β*_0_ is the intercept, - *β*_1_ is the coefficient for the fixed effect (treatment or genotype), - *X_ij_* is the fixed effect predictor (treatment or genotype) for cell *i* in well *j*, - *u_j_* is the random effect for well *j*, assumed to be normally distributed with mean 0 and variance *σ*^2^, - *ɛ_ij_* is the residual error for cell *i* in well *j*, assumed to be normally distributed with mean 0 and variance *σ*^2^.

Where indicated, measurements were combined over multiple differentiation batches (independent staining and imaging experiments). To this end, an equal number of cells from each experimental condition were sampled uniformly per batch, fluorescent values were z-scaled within that batch, and then combined. Indicator vectors for well-of-origin and batch-of-origin were included in the model. Before applying this linear transformation, per-cell per-image clipping was determined to be low (< 0.1%) and the response function of the confocal microscope was assumed to be linear.

For each condition, representative images were chosen from a single batch as the images closest to the mean fluorescence intensity for each condition. Voxels not belonging to a cell (i.e., not used in quantification) were masked prior to mean-projection for visualization.

**Quantification of live single-plane imaging** Single-plane live imaging data were binarized using a threshold set at the 75th intensity percentile for each channel of interest (TMRM or CellROX), and mean fluorescence intensities within these masked regions were calculated following previously established methodology ( [119]). The standardized intensity threshold (75th percentile) was empirically selected to reliably identify high-fluorescence areas across experimental conditions.

**Quantification of live single-plane imaging with FCCP** For time-course experiments involving FCCP treatment, images acquired before and after treatment were aligned using Fourier-based image registration. Spatial shifts between time points were estimated by phase cross-correlation and corrected using Fourier transformations. Alignment accuracy was manually verified by visual inspection. A binary mask defining regions of high fluorescence intensity was generated from the 75th percentile threshold of the initial (baseline) TMRM image. This baseline-derived mask was consistently applied across all subsequent time points to quantify mean fluorescence intensities within these regions.

### Oxygen Consumption Rate Data Analysis

The oxygen consumption rate (OCR) of cells was determined over time using a Seahorse XF Analyzer. Prior to analysis, OCR curves were visually inspected in a blinded manner to exclude wells that did not respond to drug injections. To calculate per-well total oxygen consumption for a given experimental period (e.g., under basal conditions prior to injections of uncouplers), integrals between specific experimental time points were computed from the OCR curve. The following measurements were made:

1. Basal respiration was computed as the total oxygen consumption prior to oligomycin injection.
2. Proton leak was computed as the total oxygen consumed after oligomycin injection and prior to FCCP injection.
3. Maximal respiration was computed as the total oxygen consumption after FCCP and prior to Rotenone + Antimycin injection.
4. Relative uncoupling was computed as the fraction of basal respiration attributed to proton leak.
5. Spare respiratory capacity was determined as the ratio of maximal respiration to basal respiration.

#### LC-MS Data Analysis

**LC-MS Lipidomics Data Analysis.** Lipids were identified, and their signals integrated using the Lipidsearch © software (version 4.2.27, Mitsui Knowledge Industry, University of Tokyo). Integrations and peak quality were curated manually. Peak areas were first background-corrected by subtracting three times the median peak area measured in blank samples; negative values resulting from this correction were set to zero. Statistical comparisons between different cell lines were performed using Welch’s t-test (scipy.stats.ttest_ind, equal_var=False). For comparisons involving treatment conditions within the same cell line, Student’s t-test (scipy.stats.ttest_ind, equal_var=True) was used, assuming equal variance due to identical genetic backgrounds.

**LC-MS Metabolomics Data Analysis.** Data were analyzed using Compound Discoverer 3.2 (Thermo Fisher Scientific). Metabolite identification was based either on MS2/MS3 spectral matching against a local mzVault library and corresponding retention times from pure standards (Level 1), or spectral matching using mzCloud (Level 2). Each metabolite identification was manually inspected. Peak areas were first background-corrected by subtracting three times the median peak area measured in blank samples; negative values resulting from this correction were set to zero. Median-centered peak areas were scaled (StandardScaler() from scikit-learn) prior to principal component analysis (PCA). The Harvard Center for Mass Spectrometry identified three samples with notably low overall metabolite intensities, which were subsequently excluded from downstream analyses.

**Targeted LC-MS Metabolite Analysis in Media Samples.** Peak areas from targeted metabolite analysis of media samples were compared for CDP, CDP-choline, and choline. To ensure accurate detection, solvent blanks were analyzed: CDP and CDP-choline were not detected in these blanks, while choline was detected at levels several orders of magnitude lower than in media samples.

#### Bulk mRNA Sequencing of iNs

**Bulk mRNA Sequencing Data Processing.** Sequencing data were processed using the BMC/BCC pipeline version 1.8 (updated 06/06/2023). Further details of the pipeline are available at: https://openwetware.org/wiki/BioMicroCenter:Software#BMC-BCC_Pipeline by the MIT BioMicro Center. Fastq files were subsequently processed in-house using STAR and featureCounts. Human reference transcriptome (GRCh38.p14, GENCODE release 47) and annotation files were downloaded from GENCODE and indexed using STAR (version recommended parameters). Sequencing reads were trimmed using Trim Galore with Nextera- specific settings and a stringency of 3, requiring a minimum overlap of 3 bases with adapter sequences for trimming. Trimmed reads were then mapped to the indexed human genome using STAR, with paired-end reads processed concurrently. Gene-level counts were generated using featureCounts with settings optimized for paired-end data: counting only read pairs with both ends aligned (-B), excluding pairs mapping to different chromosomes or strands (-C), and counting fragments rather than individual reads (-p). Counts were summarized at the exon level, and grouped by gene identifier.

**Differential gene expression analysis.** Differential gene expression analysis was performed using edgeR and limma-voom. Counts were filtered to retain protein-coding genes expressed at a minimum of 1 count-per-million (CPM) in at least one sample. Normalization factors were calculated, and linear modeling was conducted using limma’s voom method, followed by empirical Bayes moderation (eBayes). Statistical comparisons were performed using contrasts tailored to experimental conditions (treatment and genotype) within batches, and results were summarized as log-fold changes with associated p-values.

**Gene set enrichment analysis.** Gene set enrichment analysis was conducted using Fast Gene Set Enrichment Analysis (FGSEA). Differentially expressed genes were ranked by a score calculated as the sign of log-fold change multiplied by the negative log-transformed p-value. Pre-defined gene sets were evaluated with FGSEA using 10,000 permutations, and significant pathways were identified based on adjusted p-values < 0.05. Leading-edge genes from these significant pathways were partitioned along with their associated pathways as described in “Gene-pathway clustering using Kernighan-Lin heuristic.”

**Gene-Pathway K/L Cluster Similarity Analysis.** Jaccard indices were computed to assess similarity between gene-pathway clusters. Empirical p-values for the overlaps were obtained by comparing observed overlaps against 1000 random permutations, and p-values were adjusted using the Benjamini-Hochberg method to control the false discovery rate.

#### Single-cell Transcriptomic Data Processing

**Variant calling and ROSMAP subject selection.** A total of 36 individuals were selected from the ROSMAP cohort, a longitudinal cohort study of aging and dementia in elderly nuns, priests, and brothers. Processed whole genome sequencing (WGS) variant call files for all ROSMAP samples, where available (*N* = 1249 sequencing samples), were downloaded from Synapse (syn11724057). Variant call data were downloaded for chromosomes harboring SORL1, TREM2, ABCA7, ATP8B4, ABCA1, and ADAM10 (see Github Repository). When more than one WGS sample existed for a given subject, the sample with the higher Genomic Quality Score was chosen. Only samples that did not have sex mismatches and were consistent with previous array-based genotype data were considered (see syn12178037). Only variants that passed quality control (‘FILTER*_P_ ASS*^′^)*wereconsidered*.

Potential PTC (protein-truncating) variants in each of the aforementioned genes were flagged based on the following criteria: the variant had to be either a splice, missense, frameshift, nonsense, or premature start variant and be annotated as ‘LOF’ (loss of function). For ABCA7, this filtering captured known ABCA7 LoF risk variants from the literature, except for c.5570+5G>C, which was manually added to the filtered variants. Also see syn10901595 (https://www.synapse.org/#!Synapse:syn10901595) for information on WGS library preparation, quality control, variant annotations, and impact predictions. Annotated ABCA7 PTC variants are shown in Data S1.

The WGS data was used to identify 12 subjects who did not carry a known PTC variant in one of the aforementioned genes, other than in ABCA7, and for whom fresh-frozen postmortem tissue was available for request from Rush University (termed ‘LoF’ samples). We also selected 24 individuals who do not carry a single ABCA7 PTC mutation or PTC variants in one of the aforementioned genes (termed ‘control’ samples). Control samples were matched on age, sex, and pathology.

**Read Counting Alignment.** Library demultiplexing was performed using the BMC/BCC pipelines BioMicroCenter Software. Fast-q reads were aligned to the human genome GRCh38 and counted using the ‘cellranger count()‘ function from Cell Ranger version 6.1.2 (10x Genomics). Introns were included in counting to allow for the detection of unspliced transcripts, and the expected number of cells was set to 5000. Otherwise, Cell Ranger (v.6.1.2) default parameters were used. Counts across individual samples were then aggregated using a custom aggregation script (see GitHub Repository), resulting in a total of 150,456 cells.

**Sample Swap Analysis.** Sample swap analysis was performed using a previously established pipeline (MVV; QTLtools_1.1) [120], which compares allelic concordance between genomic and transcriptomic sequencing data. As input, we used the BAM files generated in the cellranger counting step and the chromosome 19 (the chromosome harboring ABCA7) variant call files (VCF). When comparing the concordance of BAM and VCF data for homozygous and heterozygous sites, the expected WGS sample appeared as a clear outlier (more consistent along both dimensions than any of the other 1249 WGS ROSMAP samples) for all single cell samples (Figure S1).

**Cell filtering metrics.** Prior to cell type annotation, we performed a series of quality control steps on the aggregated counts matrix. First, we filtered cells based on *N_g_*, the number of genes for each cell where counts *>* 0, and kept cells for which 500 *< N_g_ <* 10000. Next, we removed all cells with a high fraction of counts from mitochondrial-encoded genes. Mitochondrial fraction (*M_f_* ) is a commonly used per-cell metric to measure compromised nuclear integrity, with high fractions indicating low-quality nuclei, where *C_mt_* is the total counts of mitochondrially-encoded genes for a cell, *C_t_* is the total count of all genes for the same cell, and 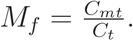.

We fit a Gaussian mixture model (GMM) using sklearn’s GaussianMixture implementation to ^′^ = log_10_(*M_f_*+*ɛ*), where *ɛ* is a small value added to *M_f_*to avoid taking the logarithm of zero.

A GMM models the data as independently sampled from a mixture of *k* Gaussian probability densities parameterized by a mean *µ_k_*, a variance *σ_k_*, and a mixture weight *π_k_* indicating the proportion of the data derived from each component. The following log-likelihood function was maximized: 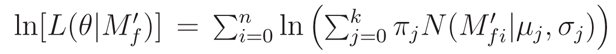 The model with *k* = 5 components had the lowest Bayesian information criterion (BIC) score (GridSearchCV from sklearn.preprocessing), where: BIC = −2 ln(*L*) + *k* ln(*n*) where *L* is the maximized log-likelihood of the model, and *n* is the number of observations (cells). Finally, each cell *i* is assigned a component in *k* according to: argmax*_k_* ^è^*π_k_N* (*M* ^′^ |*µ_k_, σ_k_*)^l^ Cells assigned to the component *k* with the highest mean *M* ^′^ scores were presumed to constitute a population of low-quality cells and were removed from further analysis. This initial filtering removed approximately 20,000 cells.

Considering all remaining cells in marker-gene expression space, where marker genes include only known cell type-specific genes for the major human PFC cell types, including astrocytes (159 markers), excitatory neurons (113 markers), inhibitory neurons (83 markers), microglia (97 markers), oligodendrocytes (179 markers), OPCs (143 markers), and vascular cells (124 markers) (Reference 1; Table S6), normalized to total library size *NC_m_* = *Cm* where *NC_m_* and *C_m_* are respectively the normalized and unnormalized count values for a given marker gene *m* and *C_t_* is the total counts of all genes for the same cell. Next, we performed a memory-efficient implementation of singular value decomposition (IncrementalPCA from sklearn.decomposition) to transform cells from the marker-gene space (mean-centered and unit-variance) into a lower dimensional space (top 50 principal components sorted by variance). Visually, the cells formed a number of Gaussian-like clusters when projected onto the first two principal components. Under the assumption that each Gaussian cluster represented a different cell type in the brain, we again fit a GMM, as described above, except this time parameterized by a covariance matrix Σ*_k_* instead of variance *σ_k_*, to the projected data. The model with *k* = 10 and full covariance had the lowest BIC score. Each resulting cell cluster was enriched for a subset of major cell type markers in the brain, indicating clusters of astrocytes, microglia, OPCs, oligodendrocytes, excitatory neurons, inhibitory neurons, and a heterogeneous cluster of vascular cells.

To remove cells that were not well-explained by the GMM and likely represent low-quality cells, we next computed the per-cell log-probability given the model *L_i_* = ln[*P* (*x_i_*|*θ*)], using Sklearn’s GaussianMixture ’score_samples’ function, and removed cells with *L_i_ <* −100. We also removed two Gaussian clusters whose probability distributions constituted clear outliers compared to remaining clusters. The excluded cells had lower *C_t_* and higher *M_f_* compared to those that passed the log-likelihood filter, suggesting that the removed cells were indeed of low quality. As expected, when examining the data visually projected onto the first two principal components, this filtering removed many of the cells that were not visibly associated with a main Gaussian cluster. Together, this filtering removed an additional approximately 12,000 cells, leaving a total of 118,668 cells.

**Gene filtering metrics.** For the remaining downstream analysis we only considered genes that were both nuclear-encoded and protein-coding, which constituted a total of 19384 genes, based on annotation of ensembl GRCh38p12.

**Cell type annotations.** To remove variance explained by sequencing batch and individual- of-origin, we first applied the Python implementation of the Harmony algorithm [121] with individual-of-origin as an indicator vector to the low-dimensional embedding of cells (first 50 principal components) remaining after the initial rounds of quality control described above. Next, we computed a neighborhood graph on the Harmony-corrected values in the PC embedding space, as implemented in the Scanpy Python package [122], using default parameters. Finally, we applied the Leiden graph-clustering algorithm to cluster this neighborhood graph of cells, using the Scanpy implementation of the Leiden algorithm [123].

We used the Scanpy ’rank_genes_groups’ function to compute top marker genes per Leiden cluster. Briefly, we assigned a major cell type label *c* to each Leiden cluster, where *c* ∈ {’Ex’, ’In’, ’Ast’, ’Mic’, ’Oli’, ’Opc’, ’Vascular’}, by computing the average cell-type-specific marker gene enrichment per Leiden cluster. Specifically, this is a vector of cell type signatures *S*_c_ for each Leiden cluster, where 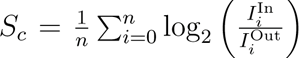 where *n* is the total number of marker genes assigned to a cell type in *c* and *I*^In^ and *I*^Out^ indicate average gene expression values for a gene *i* for cells inside or outside a specific Leiden cluster, respectively. Then each Leiden cluster is assigned a label *c* by argmax*_c_*(*S_c_*). Finally, we sub-clustered cells from each major cell type using the Leiden clustering algorithm and examined distributions of mitochondrial fractions *M_f_* and total counts *C_t_* among subclusters *s* of the same cell type. Clusters were removed if *M_f_ >* (2 · std(*M_f_* ) + *M_f_* ) or if |*C_t_* | *>* (2 · std(*C_t_*) + *C_t_*), where *M_f_* and *C_t_* are respectively the mean *M_f_* and *C_t_* for all cells in a given Leiden cluster *s* and *M_f_* and *C_t_* are respectively the means of those values across all Leiden clusters, considering only clusters with the same cell type annotation *c*, because variance in *M_f_* and *C_t_* across major cell types can be biologically explained. Manual inspection of the removed clusters revealed that they tended to have fewer cells and low individual-level representations, and were not well-connected in the graph.

**Individual-level filtering.** After all rounds of quality control as described above, we noted a subset of individuals (*N* = 6) with very few cells (*<* 500). These subjects were removed from further analysis, resulting in 24 control individuals and 12 ABCA7 LoF individuals. None of these individuals carried ABCA7 PTC variants, and removing them did not substantially alter the distribution of clinical variables across genotypes.

**Differential gene expression.** Summed (pseudo-bulked) gene expression values were computed by matrix multiplication *XI*, where *X* is the gene x cell counts matrix and *I* is a cell x individual binary matrix indicating the individual-of-origin for each cell, resulting in 36 gene expression vectors for each of the six major cell types. For each cell type, only genes with a nonzero detection rate *>* 0.10 were considered for differential expression. Summed counts were normalized using the edgeR TMM method. The residual mean-variance trend not explained by the multivariate linear model (formalized below) was removed using Limma-Voom. Unknown sources of variance were captured in the model using surrogate variable analysis (SVA). Limma’s lmFit, eBayes, and topTable functions were then used to estimate differential gene expression statistics, as reported in Data S4. The following model was fit for each cell type: *G_i_* = *β*_0_ ×ABCA7LoF+*β*_1_ ×msex +*β*_2_ ×nft +*β*_3_ ×amyloid+*β*_4_ ×age_death+ *β*_5_ × PMI + *β*_6_ × batch + *β*_7_ × APOE4 + *β*_8_ × SV0 *G_i_* refers to a vector of expression profiles of size 1 × 36 for a gene *i* in a given cell type. ABCA7LoF is a binary variable, encoding the presence of an ABCA7 variant predicted to cause loss of function (see Data S2). See Supplementary Text for descriptions of the remaining variables included in the model. SV0 refers to the first surrogate variable estimated from the data. The exact number of surrogate variables per cell type to include as additive terms in the model was estimated using the num.sv() function in R.

**Gene-pathway projections.** For each cell type, we computed a set of gene-wise scores quantifying the direction and statistical significance of gene expression changes (computed as part of the differential gene expression analysis) associated with ABCA7 LoF: *S* = sign(log_2_ FC) × − log_10_(p-value) where log_2_ FC *>* 0 indicates up-regulation in ABCA7 LoF vs control. Top differentially expressed genes per cell type (|*S*| *>* 1.3) were projected from 6-dimensional score space, where each dimension captures ABCA7 LoF perturbation scores in one of the major cell types (Ex, In, Ast, Mic, Oli, OPC), into two dimensions, using the UMAP algorithm (using the ‘umap‘ Python package). Gene scores that were not detected in >10% of cells in a given cell type were set to 0.

We performed a grid search for Gaussian mixture parameters (parameter 1: number of components; parameter 2: covariance type) on the embedded cells (using the Python ‘sklearn‘ package) to assign genes to clusters in the 2D space. We proceeded with the model with the lowest BIC score, which had 15 components and a tied covariance matrix.

Each cluster was assigned representative pathway names by testing genes in that cluster for enrichment with Gene Ontology Biological Process pathways (Table S6) against the background of all genes in the embedding space, by hypergeometric enrichment (using the Python package ‘gseapy‘). Pathways with an enrichment p-value < 0.01 were considered for cluster annotation.

Per-cell-type perturbation scores (*S_c_*) for each cluster were computed as the average gene score *S* (for a given cell type) for all genes in that cluster. The statistical significance of each cell type-specific cluster score was assessed by permuting cluster assignments (100,000 permutations).

**Gene-set enrichment.** Genes were rank-ordered based on their scores *S* (see description in Gene-Pathway Projections). An R implementation of gene set enrichment analysis (GSEA) [124] (fast gene set enrichment analysis, fGSEA) was run with 10,000 permutations to estimate the statistical overrepresentation of gene sets in the WikiPathways databases (Table S6) within high-scoring (|*S*|), differentially expressed genes. Gene sets with a minimum size of 5 and a maximum size of 1000 were considered.

**Gene-pathway clustering using Kernighan-Lin heuristic.** To reduce the solution’s computational search space, we reformulated the gene-pathway association problem as a bipartite graph *G* constructed from all the genes in the Leading Edge subset (LE) and their associated pathways. LE was defined as the set of 268 genes driving the enrichment signal for pathways that passed a significance threshold of *p <* 0.05 (fGSEA) in Con vs. ABCA7 LoF excitatory neurons. *G* was constructed from an *n* × *m* unweighted adjacency matrix, where *n* represented the number of LE genes and *m* represented the number of pathways associated with four or more LE genes, as specified in the WikiPathways database.

We chose to group gene-pathways into clusters of approximately equal size, making this a graph partitioning problem. We found that removing this constraint made the grouping results highly susceptible to outliers (Supplementary Text; Figure S7C). Of the three graph partitioning algorithms tried, METIS and the Kernighan-Lin (K/L) algorithms had the lowest loss (Supplementary Text; Figure S7B). Both METIS and K/L achieved very comparable losses (within 1.8% of each other, after 5.0 × 10^4^ random initiations) and produced almost identical solutions (Rand index=0.98, after 5.0 × 10^4^ random initiations) (Supplementary Text; Figure S7B, D-F). We proceeded with the K/L algorithm for gene-pathway groupings as we found this algorithm to perform consistently better than METIS across a wider range of graph sizes (not shown).

The K/L algorithm was implemented in Python (see GitHub Repository) based on its original paper [125] and run with parameters set as *C* = 0, *KL*_*modif ied* = *True*, *random*_*labels* = *True*, *unweighted* = *True*, and *K* = 50 to partition *G* into 8 groups. We performed 5.0 × 10^4^ random initiations on *G* and report the partitioning with the lowest loss among all initiations.

Gene-pathway graph layouts were computed using the ‘networkx‘ Python package with the spring layout algorithm, using 10,000 iterations. Layouts were visualized using the ‘matplotlib‘ ‘pyplot‘ package in Python.

Representative pathways for each cluster were inferred from the graph by averaging the ABCA7 LoF perturbation scores *S* for all genes in the cluster of interest sharing an edge with the pathway in question. Scores for pathways with intra-cluster degrees ≥ 5 were reported in the figures. Manually picked subsets of genes with the largest scores (|*S*| *>* 1) were reported in the figures. All gene statistics are reported in Data S4, and cluster assignments are reported in Data S7.

**Excitatory neuronal layer annotation.** Excitatory neurons were annotated by cortical layer using previously published marker gene sets [126] (Table S6). The normalized gene expression matrix (post-qualtiy control described above) for excitatory neurons was filtered to include only layer-specific marker genes and cells expressing at least 15% of these genes. Dimensionality was reduced using iterative principal component analysis (iPCA), and batch effects from individual subjects were corrected using Harmony. A neighborhood graph was constructed based on these corrected components, followed by Leiden clustering to identify neuronal clusters. Clusters were annotated by calculating the average log-fold change of layer-specific marker genes. Clusters with significant enrichment (average log-fold change > 0.1) were labeled by cortical layer, while ambiguous clusters were removed from further analysis. Layer 5 and 6 annotations were combined into a single ’L5/6’ category. These annotations were confirmed using marker genes from an independent study [127] (Table S6). Per-layer differentially expressed genes were computed as described in the section “Differential Gene Expression,” followed by gene set enrichment analysis of ABCA7 LoF-associated K/L clusters in excitatory neurons as described in the section “Gene Set Enrichment.”

**ABCA7 p.Ala1527Gly variant calling and gene-pathway clustering comparisons.** We followed the same steps indicated in the section “Variant Calling and ROSMAP Subject Selection” to identify subjects who carried the p.Ala1527Gly variant and had snRNAseq performed on the PFC as part of a previous study. Differentially expressed genes were computed as described in the section “Differential Gene Expression,” followed by gene set enrichment analysis of ABCA7 LoF-associated K/L clusters in excitatory neurons as described in the section “Gene Set Enrichment.”

#### Molecular Dynamics Simulations

The initial structure of ABCA7 was obtained from the Protein Data Bank (PDB) in unbound- open and bound-closed conformations, PDB IDs 8EE6 and 8EOP, respectively. These experimentally solved structures harbor the G1527 variant. The A1527 structure was generated by mutating the glycine residue to alanine using pymol software.

The ABCA7 domain between residues 1517 and 1756 was embedded in a dipalmitoylphosphatidylcholine (DPPC) membrane using the CHARMM-GUI web server and oriented according to the Orientations of Proteins in Membranes (OPM) database. Four different simu- lations were performed using GROMACS 2022.3, as reported in Table S3. The CHARMM36M force field was used for all simulations.

The protein-membrane system was solvated in a cubic box with a minimum distance of 1.0 nm between the protein and the box edge, using the TIP3P water model. Energy minimization was performed using the steepest descent algorithm with a maximum force threshold of 1000 kJ/mol/nm to relieve any steric clashes or bad contacts. The system was equilibrated in six phases, each 125 ps long, to equilibrate volume (NVT) and pressure (NPT). The production run, 300 ns long, was performed in the NPT ensemble at 323 K using a v-rescale thermostat and 1 bar using the Parrinello-Rahman barostat. A 2 fs time step with h-bonds constraints was used with periodic boundary conditions applied in all directions. Long-range electrostatics were handled using the Particle Mesh Ewald (PME) method with a cutoff of 1.0 nm for non-bonded interactions.

RMSD was calculated to monitor the conformational stability of a given structure over the course of the simulation by comparing the position of *C_α_* at time *t* under simulation to its reference position (in 8EOP or 8EE6).The *ϕ* and *ψ* dihedral angles were calculated using the *gmx rama* tool, followed by post-processing. Secondary structure analysis was performed using *gmx dssp -hmode dssp*, with subsequent post-processing using custom Python scripts.

Principal Component Analysis (PCA) was conducted to identify the major conformational changes during the simulation. The analysis involved the following steps:

1. A covariance matrix of the *C_α_* atom positional fluctuations was constructed using the ‘gmx covar‘ tool.
2. The covariance matrix was diagonalized to obtain the eigenvalues and eigenvectors, representing the principal components (PCs).
3. The trajectory was projected onto the first two principal components (PC1 and PC2) using the ‘gmx anaeig‘ tool to visualize the dominant motions.
4. A kernel density estimate (KDE) plot implemented in seaborn python3 was used to visualize the first two eigenvectors for each simulation, corresponding to 45%, 40%, 40% and 33% of variance in 8EOP-G1527, 8EE6-A1527, 8EE6-G1527 and 8EOP-A1527 respectively.

Visualization of the trajectories was carried out using VMD software.

### Supplementary Text

#### Description of variables according to the Rush Alzheimer’s Disease Center Code- book

1. **age_death**. Age at death
2. **amyloid**. Overall amyloid level - Mean of 8 brain regions. Amyloid beta protein identified by molecularly-specific immunohistochemistry and quantified by image analysis. Value is percent area of cortex occupied by amyloid beta. Mean of amyloid beta score in 8 regions (4 or more regions are needed to calculate). The 8 regions are hippocampus, entorhinal cortex, midfrontal cortex, inferior temporal gyrus, angular gyrus, calcarine cortex, anterior cingulate cortex, superior frontal cortex.
3. **braaksc**. Braak stage. Semi quantitative measure of neurofibrillary tangles. Braak Stage is a semi quantitative measure of severity of neurofibrillary tangle (NFT) pathology. Bielschowsky silver stain was used to visualize NFTs in the frontal, temporal, parietal, entorhinal cortex, and the hippocampus. Braak stages were based upon the distribution and severity of NFT pathology: Braak stages I and II indicate NFTs confined mainly to the entorhinal region of the brain; Braak stages III and IV indicate involvement of limbic regions such as the hippocampus; Braak stages V and VI indicate moderate to severe neocortical involvement.
4. **ceradsc**. CERAD score. Semiquantitative measure of neuritic plaques.CERAD score is a semiquantitative measure of neuritic plaques. A neuropathologic diagnosis was made of no AD (value 4), possible AD (value 3), probable AD (value 2), or definite AD (value 1) based on semiquantitative estimates of neuritic plaque density as recommended by the Consortium to Establish a Registry for Alzheimer’s Disease (CERAD), modified to be implemented without adjustment for age and clinical diagnosis. A CERAD neuropathologic diagnosis of AD required moderate (probable AD) or frequent neuritic plaques (definite AD) in one or more neocortical regions. Diagnosis includes algorithm and neuropathologist’s opinion, blinded to age and all clinical data. Value 1: definite AD, Value 2: probable AD, Value 3: possible AD, Value 4: no AD.
5. **cogdx**. Final consensus cognitive diagnosis. Clinical consensus diagnosis of cognitive status at time of death. At the time of death, all available clinical data were reviewed by a neurologist with expertise in dementia, and a summary diagnostic opinion was rendered regarding the most likely clinical diagnosis at the time of death. Summary diagnoses were made blinded to all postmortem data. Case conferences including one or more neurologists and a neuropsychologist were used for consensus on selected cases. Value 1: NCI, No cognitive impairment (No impaired domains), Value 2: MCI, Mild cognitive impairment (One impaired domain) and NO other cause of CI, Value 3: MCI, Mild cognitive impairment (One impaired domain) AND another cause of CI, Value 4: AD, Alzheimer’s disease and NO other cause of CI (NINCDS PROB AD), Value 5: AD, Alzheimer’s disease AND another cause of CI (NINCDS POSS AD), Value 6: Other dementia. Other primary cause of dementia
6. **msex**. Sex. Self-reported sex, with “1” indicating male sex. 1 = Male 0 = Female
7. **nft**. Neurofibrillary tangle burden. Neurofibrillary tangle summary based on 5 regions. Neurofibrillary tangle burden is determined by microscopic examination of silver-stained slides from 5 regions: midfrontal cortex, midtemporal cortex, inferior parietal cortex, entorhinal cortex, and hippocampus. The count of each region is scaled by dividing by the corresponding standard deviation. The 5 scaled regional measures are then averaged to obtain a summary measure for neurofibrillary tangle burden.
8. **pmi**. postmortem interval. Time interval in hours from time of death to autopsy. postmortem interval (PMI) refers to the interval between death and tissue preservation in hours.

#### Choosing a partitioning heuristic for gene-pathway grouping

**Methods.** The heatmap in Figure S7A highlights how frequently pathways within a pathway database, such as WikiPathways, share gene members. On average, every pathway shown in Figure S7A shares at least one gene with approximately 40% of the other pathways, highlighting that there is redundancy in this matrix that could be summarized in simpler terms.

To summarize redundant gene-pathway information into a limited number of non-redundant gene-pathway groups, we reformulated the gene-pathway association problem as a bipartite graph *G*constructed from all the genes in the Leading Edge subset (LE) and their associated pathways. LE was defined as the set of 268 genes driving the enrichment signal for pathways that passed a significance threshold of *p <* 0.05 (fGSEA) in Con vs. ABCA7 LoF excitatory neurons. *G* was constructed from an *n* × *m* unweighted adjacency matrix, where *n* represented the number of LE genes and *m* the number of pathways associated with four or more LE genes, as specified in the WikiPathways database.

Graph partitioning involves segmenting the vertices of a graph into equal-sized partitions, optimizing for the minimal number of interconnecting edges (i.e., “total cut size”). We tested three prominent graph partitioning techniques, as outlined by Elsner (1997) [128], to approximate optimal partitioning. These methods include:

1. **Recursive Spectral Bisection**: Implemented in Python using the numpy linear algebra package, this method was executed for log_2_(*N* ) iterations, yielding *N* = 8 partitions. A detailed description of the algorithm can be found in Elsner (1997) [128].
2. **Multilevel Graph Partitioning**: Leveraging the METIS software package [129] in Python using the following parameters: ‘nparts=8‘, ‘tpwgts=None‘, ‘ubvec=None‘, ‘recursive=False‘.
3. **Kernighan-Lin (K/L) Algorithm**: Based on its original paper [125], this algorithm was implemented in Python and run with parameters set as *C* = 0, ‘KL_modified=True‘, ‘random_labels=True‘, ‘unweighted=True‘, and *K* = 50.

Additionally, the Spectral Clustering algorithm, a commonly used clustering method, was applied using the ‘SpectralClustering()‘ function from the ‘sklearn‘ Python package with default parameters, apart from ‘n_clusters=8‘ and ‘assign_labels=’kmeans’‘. We stipulated eight clusters for each algorithm, as qualitatively, this resolution seemed to strike a good balance to summarize main biological effects.

For benchmarking purposes, the three graph partitioning techniques and the spectral clustering algorithm were evaluated by segmenting graph *G* into eight gene-pathway clusters using the respective algorithms. Spectral clustering was run over 1,000 initiations, while K/L and METIS were run over 50,000 iterations because their solutions were slightly more variable across runs. The deterministic bisection method was run only once. A randomized graph partitioning benchmark was also computed by permuting the eight cluster labels of approximately equivalent size for 1,000 initiations. Average losses were computed per algorithm on all initiations. The benchmarking process and source code are available at: GitHub Repository.

**Results.** Spectral clustering performed significantly better than all other algorithms based on the loss (Figure S7B). This was expected, as spectral clustering does not place a constraint on cluster size. Spectral clustering results were characterized by a single large cluster and many small clusters (Figure S7C), indicating that this clustering algorithm was highly susceptible to outliers and suggesting that graph partitioning, which imposes the constraint of equal partitioning, was a better approach to the problem of grouping genes and pathways into biologically informative groups. Indeed, all three graph partitioning algorithms divided the graph into more uniformly-sized groups (Figure S7C). Among the partitioning algorithms, K/L and METIS produced the most uniformly sized groups (Figure S7C) and also had significantly lower losses compared to the spectral bisection algorithm (Figure S7B). K/L and METIS solutions were very similar, with their respective best solutions (lowest loss) having an average Jaccard similarity index of 0.91 on the diagonal (Figure S7D,E). K/L and METIS solutions were also consistent across pairwise random initiations, both when comparing within K/L or METIS solutions (Rand Index=0.87 and 0.91, respectively) and when comparing all pairwise K/L and METIS solutions (Rand Index=0.88) (Figure S7F).

Overall, these results indicate the importance of non-redundant gene-pathway groupings to interpret biological effects. They also indicate that for some gene-pathway graphs, such as the one in this study, graph partitioning is a better approach than clustering.

#### Molecular Dynamics Simulations Results

**RMSD analysis of Ala1527 vs Gly1527 in open and closed states.** To evaluate conformational stability, we conducted root mean square deviation (RMSD) analyses on ABCA7 under different states and mutations (Figure 2G,H; Figure S9A,B; Table S3). RMSD values for the *C_α_* atoms were calculated over the course of a 300 ns simulation period, comparing closed and open conformations, each harboring either the G1527 or A1527 mutation.

For the open ABCA7 conformation, both the G1527 and A1527 mutants exhibited relatively minor RMSD fluctuations (Figure S9A-D), with RMSD values for the A1527 mutant significantly lower compared to those of the G1527 mutant (Figure S9E). Overall, both mutants showed narrow RMSD distributions in the open state (Figure S9C, D), indicating generally stable conformational behavior.

Differences in RMSD distributions between the two variants became more pronounced in the closed state: The RMSD profile of the closed conformation with the G1527 mutation exhibited substantial fluctuations throughout the simulation (Figure 2I), suggesting that the G1527 mutation significantly increases local structural flexibility. In contrast, the closed conformation harboring the A1527 mutation showed only minor RMSD fluctuations (Figure 2I), suggesting that the A1527 mutation confers greater structural stability and reduced flexibility in the closed ABCA7 conformation. Principal component analysis (PCA) further highlighted these differences visually; For the closed conformation, PCA projections of G1527 conformations over time were broad, indicating significant exploration of the conformational space (Figure 2J). Conversely, the PCA plot for the closed A1527 mutant displayed a tightly clustered distribution (Figure 2J), indicating limited conformational sampling over time and suggesting decreased conformational flexibility induced by the A1527 mutation.

**Dihedral angle analysis of Ala1527 vs Gly1527 in open and closed states.** To further explore the local structural variations induced by a p.Ala1527Gly mutation, we analyzed backbone dihedral angles (phi/psi; *ϕ/ψ*) for residues 1517-1537. In the open conformation, Gly1527 consistently occupied the *α*-helical region of the Ramachandran plot throughout the simulation. In contrast, Ala1527 showed two distinct populations within the *α*-helical region (Figure S10A), suggesting subtle local conformational differences, yet overall preservation of the *α*-helical structure. These findings align closely with the RMSD analysis, and suggest similar conformational behaviors between the variants in the open state.

However, significant structural differences emerged in the closed conformation: Ala1527 displayed two preferred conformations—one within the *α*-helical region and another shifted toward the *β*-structure region—while Gly1527 explored a broader range of dihedral angles, indicative of greater structural flexibility (Figure S10A,B). This observation is in line with the RMSD analysis, indicating structural differences specifically in the closed state, with Gly1527 exhibiting significantly greater conformational flexibility compared to Ala1527.

**Secondary structure analysis of Ala1527 vs Gly1527 in open and closed states.** To complement backbone angle analysis, we also evaluated secondary structure stability throughout the simulation. In the open state, secondary structure content was comparable between variants, maintaining similar *α*-helical character (Figure S10C). Upon transitioning to the closed state, both variants experienced a substantial loss of *α*-helical content across residues 1517-1537. This loss, however, was more pronounced in the Gly1527 variant compared to the Ala1527 variant, as residues 1520-1525 retained partial *α*-helical structure more robustly in the Ala1527 variant compared to Gly1527 (Figure S10C).

Finally, structural alignment of the closed-state Gly1527 ABCA7 structure (PDB 8EOP) with the closed-state structures of ABCA1 (PDB 7TBW) and ABCA4 (PDB 7LKZ) revealed that residues corresponding to Gly1527 in ABCA7 (V1646 in ABCA1; I1671 in ABCA4) adopt stable *α*-helical structures (Figure S10E,F). In contrast, the Gly1527 residue in ABCA7 exhibits significant flexibility and lacks defined *α*-helical structure. Interestingly, our simulations indicate that the Ala1527 variant partially restores this local *α*-helical conformation in ABCA7 (Figure S10C,D). These data suggest that the Gly1527 variant induces local structural changes that differentiate ABCA7 from its close homologs ABCA1 and ABCA4.

## Supplementary Figures

**Figure S1:**
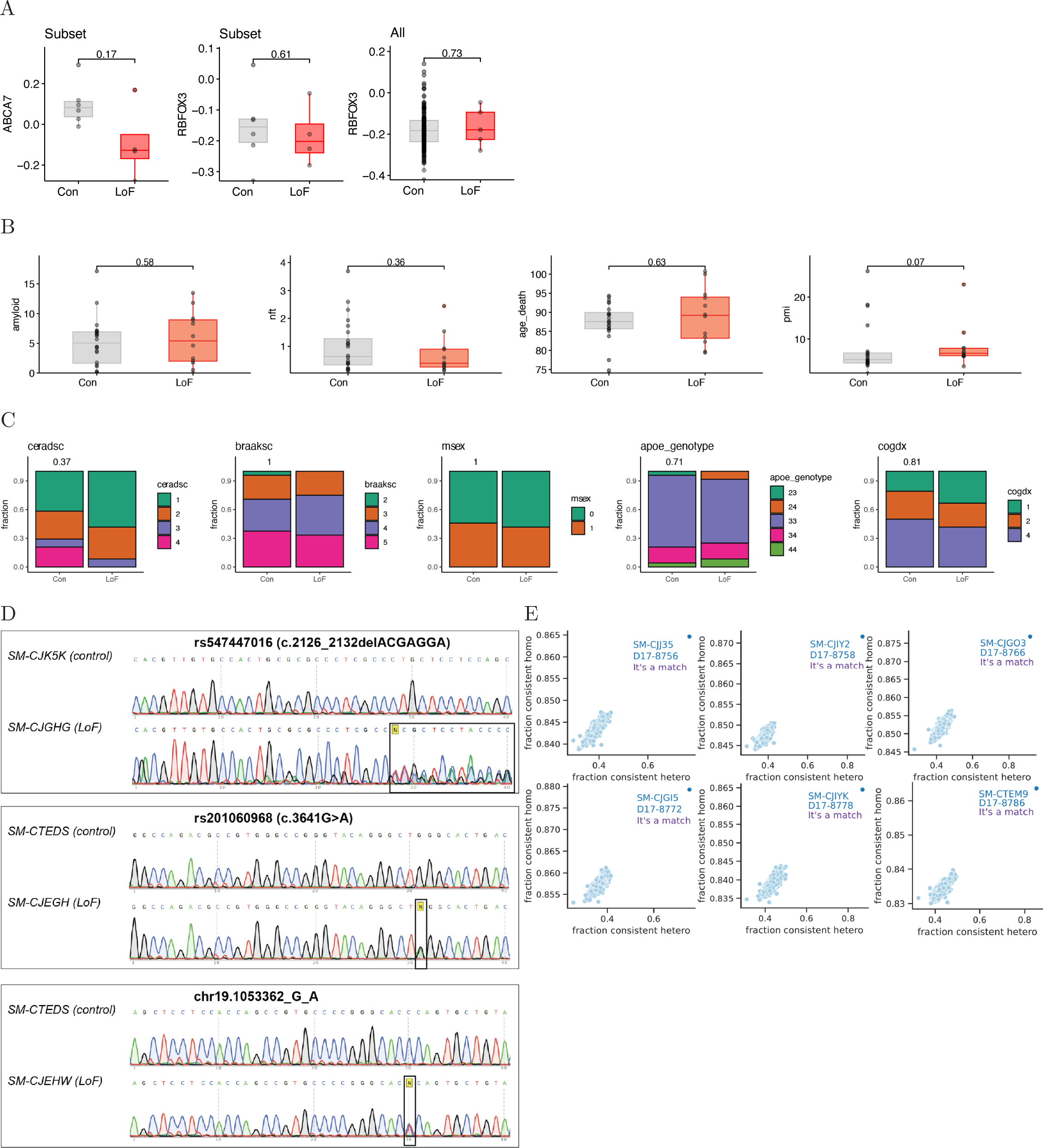
Overview of Human snRNA-Sequencing Cohort. (A) Protein levels from post-mortem human prefrontal cortex (see Table S6 for external dataset used) showing ABCA7 protein levels (left) and NeuN (RBFOX3) levels (middle) for a subset of individuals found to overlap with the snRNA-seq cohort (N=6 control and N=4 ABCA7 LoF carriers). The right panel shows NeuN (RBFOX3) protein levels by genotype in all available control samples (N=180) vs. all available ABCA7 LoF carriers with proteomic data (N=5). **(B)** Distributions of continuous metadata variables (see Supplementary Text for descriptions) for control individuals (N=24) vs. ABCA7 LoF carriers (N=12). For panels C and D, boxes indicate dataset quartiles per condition, and whiskers extend to the most extreme data points not considered outliers (i.e., within 1.5 times the interquartile range from the first or third quartile). **(C)** Distributions of discrete metadata variables for control individuals (N=24) vs. ABCA7 LoF carriers (N=12). Con=control, LoF=ABCA7 loss-of-function. P-values in panels A and B were computed by two-sided Wilcoxon rank sum test. P-values in panel C were computed by two-sided Fisher’s exact test. **(D)** Sanger sequencing of ABCA7 LoF variants in prefrontal cortex genomic DNA samples from 3 ABCA7 LoF carriers and 3 controls from the snRNA-seq cohort. Sequencing confirmed heterozygosity of the indicated variant in LoF samples, with variant location marked by a black box. Example plots validating matches between whole genome sequencing (WGS) and snRNA-seq libraries. Each plot shows the concordance of homo- and heterozygous SNP calls between WGS and snRNA-seq data for a single individual. Matches between WGS SNP calls and snRNA-seq BAM inferred SNP calls are indicated by extreme outliers. Expected (i.e., correct) matches are indicated in blue/purple.

**Figure S2:**
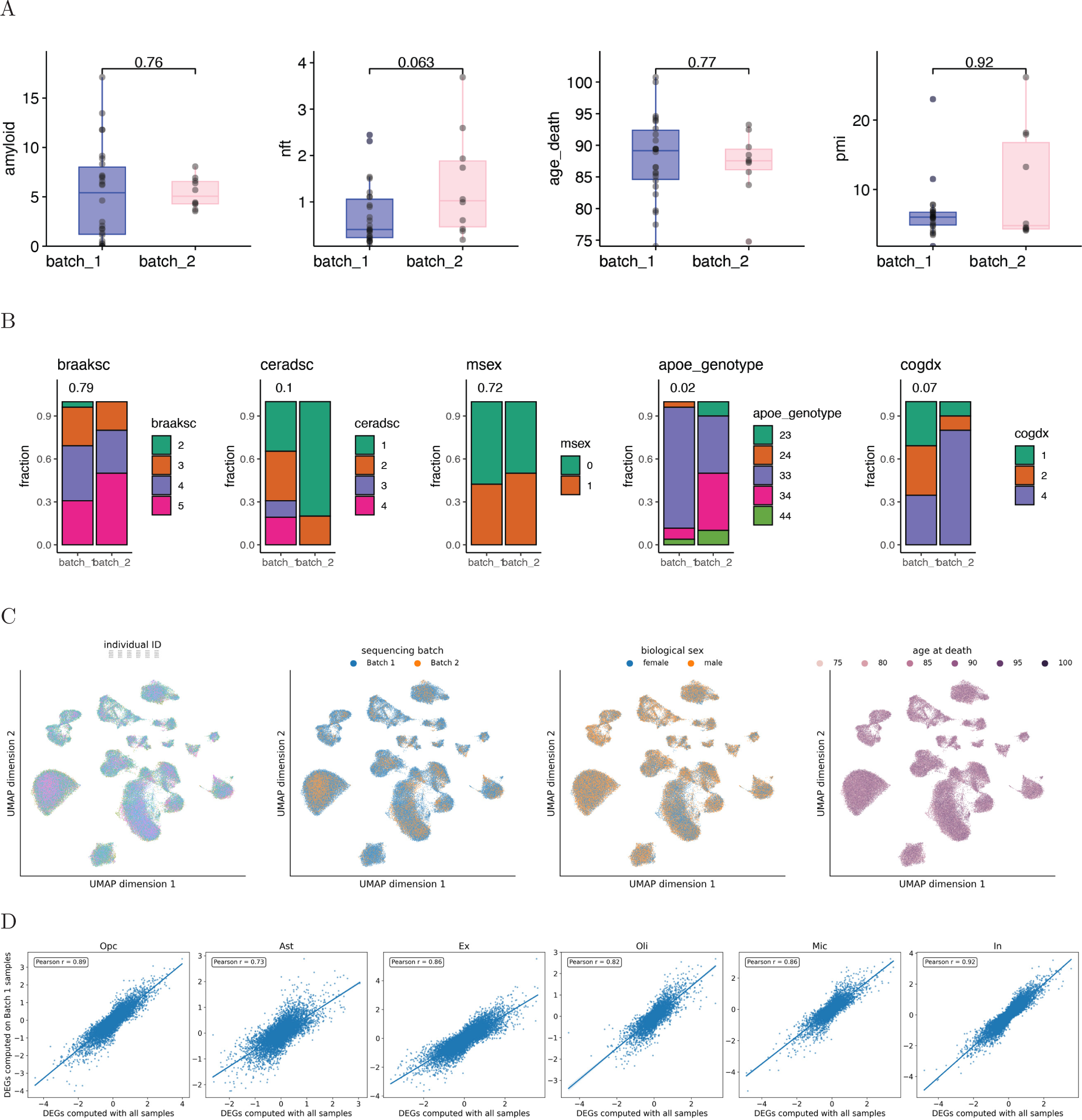
Overview of snRNA-sequencing Batch Correction and Data Quality. **(A)** Distribution of continuous metadata variables by sequencing batch. P-values in panels were computed by two-sided Wilcoxon rank sum test. **(B)** Distribution of discrete metadata variables by sequencing batch. P-values were computed by two-sided Fisher’s exact test. **(C)** 2D UMAP projection of snRNA-seq cells after quality control, colored by selected metadata variables. **(D)** Correlation of gene perturbation scores (*S* = − log_10_(*p*) × sign(log_2_(fold-change))) computed using all samples versus excluding batch 2 (v2 chemistry), demonstrating that results are robust and not driven by batch-specific effects.

**Figure S3:**
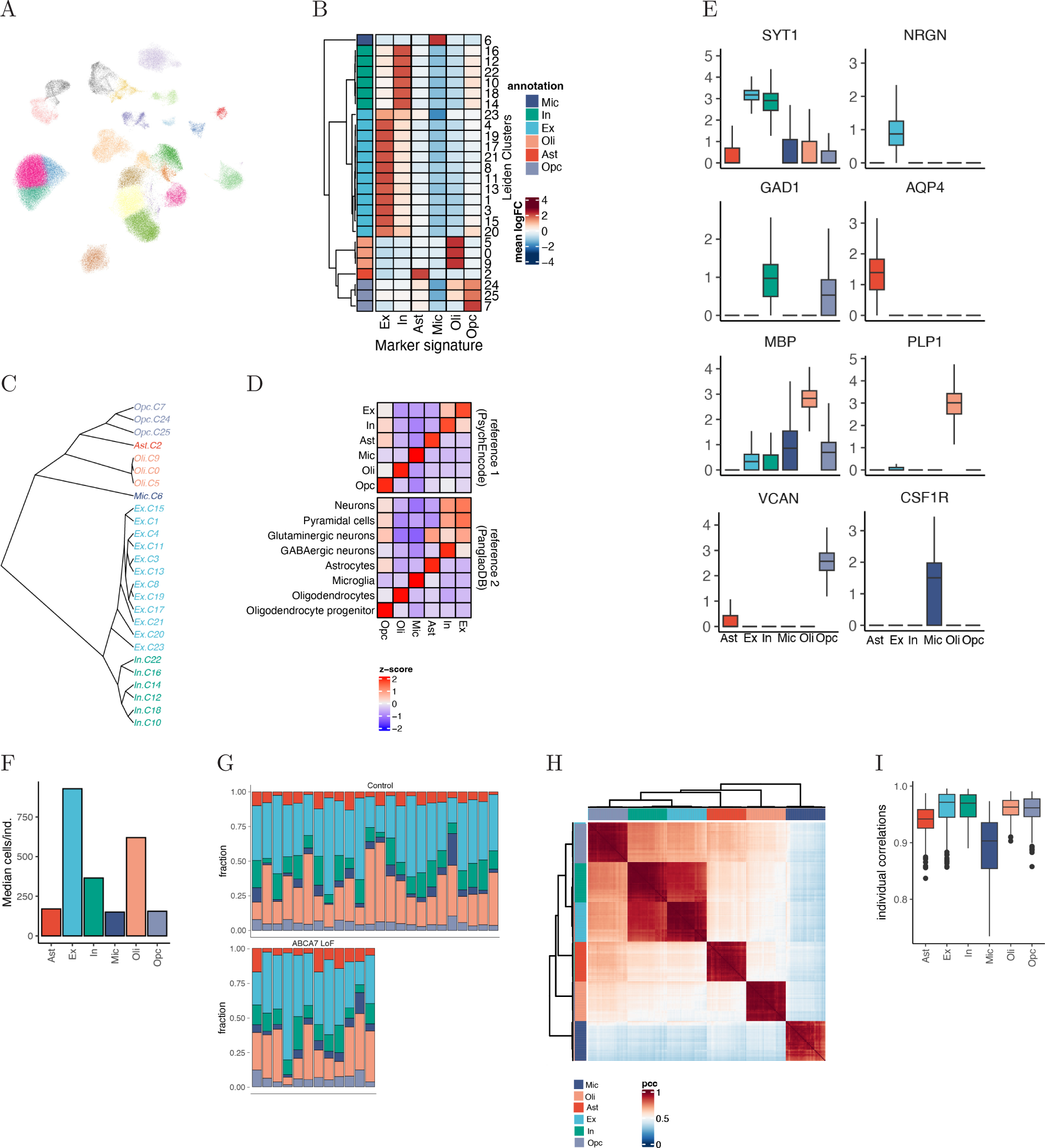
Overview of snRNA-sequencing Cell Type Annotations. **(A)** Two-dimensional UMAP projections of individual cells from gene expression space, colored by Leiden clusters. **(B)** Average marker gene expression (per-cluster mean log(fold-change)) for all marker genes for the cell type indicated along the x-axis. Log(fold-changes) are computed for the cluster of interest vs. all remaining clusters. Reference 1 (Table S6) marker genes were used. **(C)** Cladogram visualizing subcluster relationships based on pairwise distances between per-cluster gene expression profiles. **(D)** Average marker gene expression profiles (x-axis) per major cell type annotation (y-axis) for two marker gene references (Table S6). **(E)** Per-cell distribution of select marker gene expression by cell type. Y-axis indicates log-counts. **(F)** Median number of cells per cell type per individual. **(G)** Cell type fraction by individual. **(H)** Heatmap of individual-level gene expression correlations by cell type. Boxplot of individual-level gene expression correlations by cell type.

**Figure S4:**
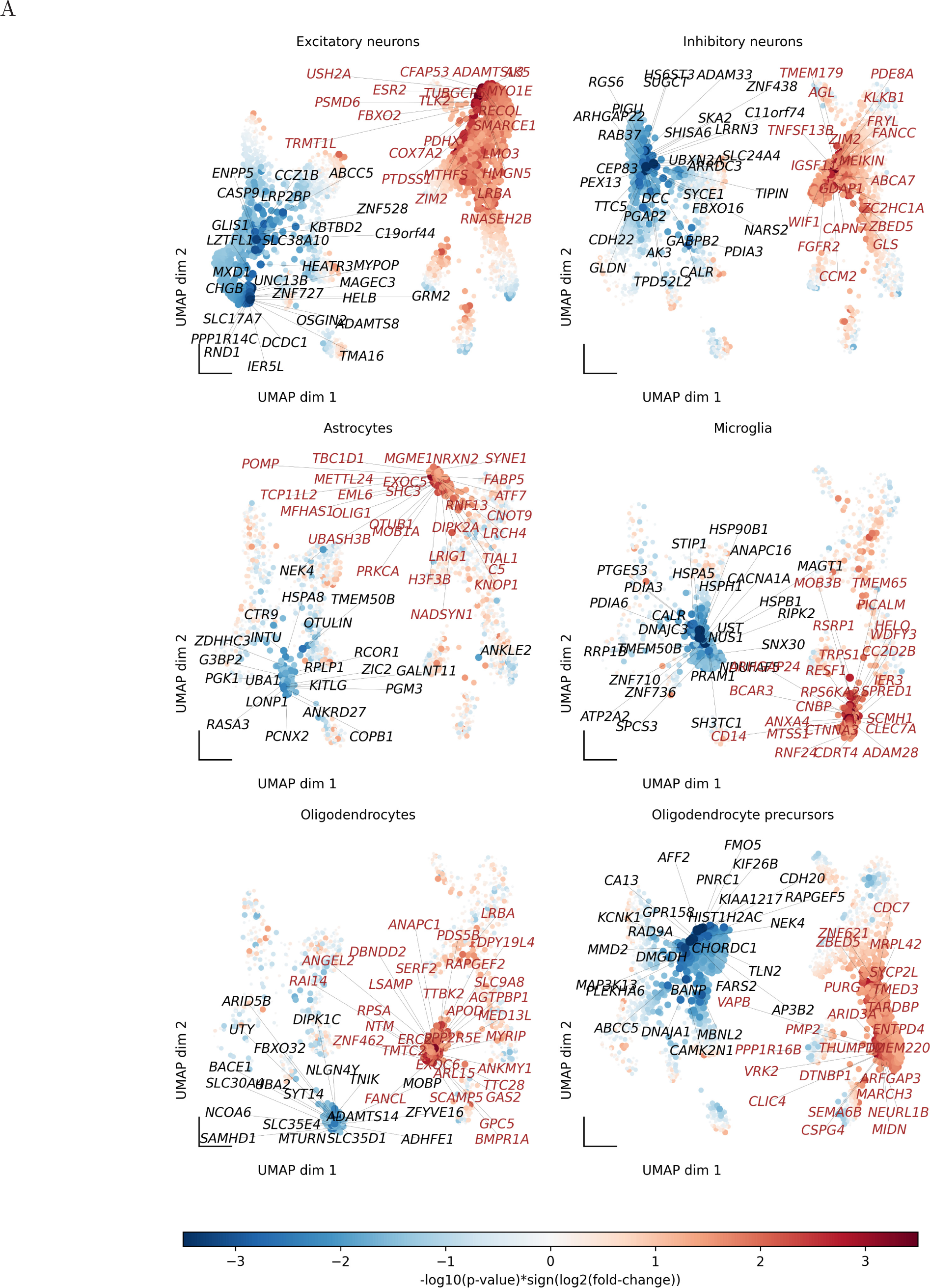
Annotated Projections of Gene Scores. Enlarged view of the UMAP projection of Figure 1E, showing the top 20 genes by absolute score (|*S*|) for each cell type.

**Figure S5:**
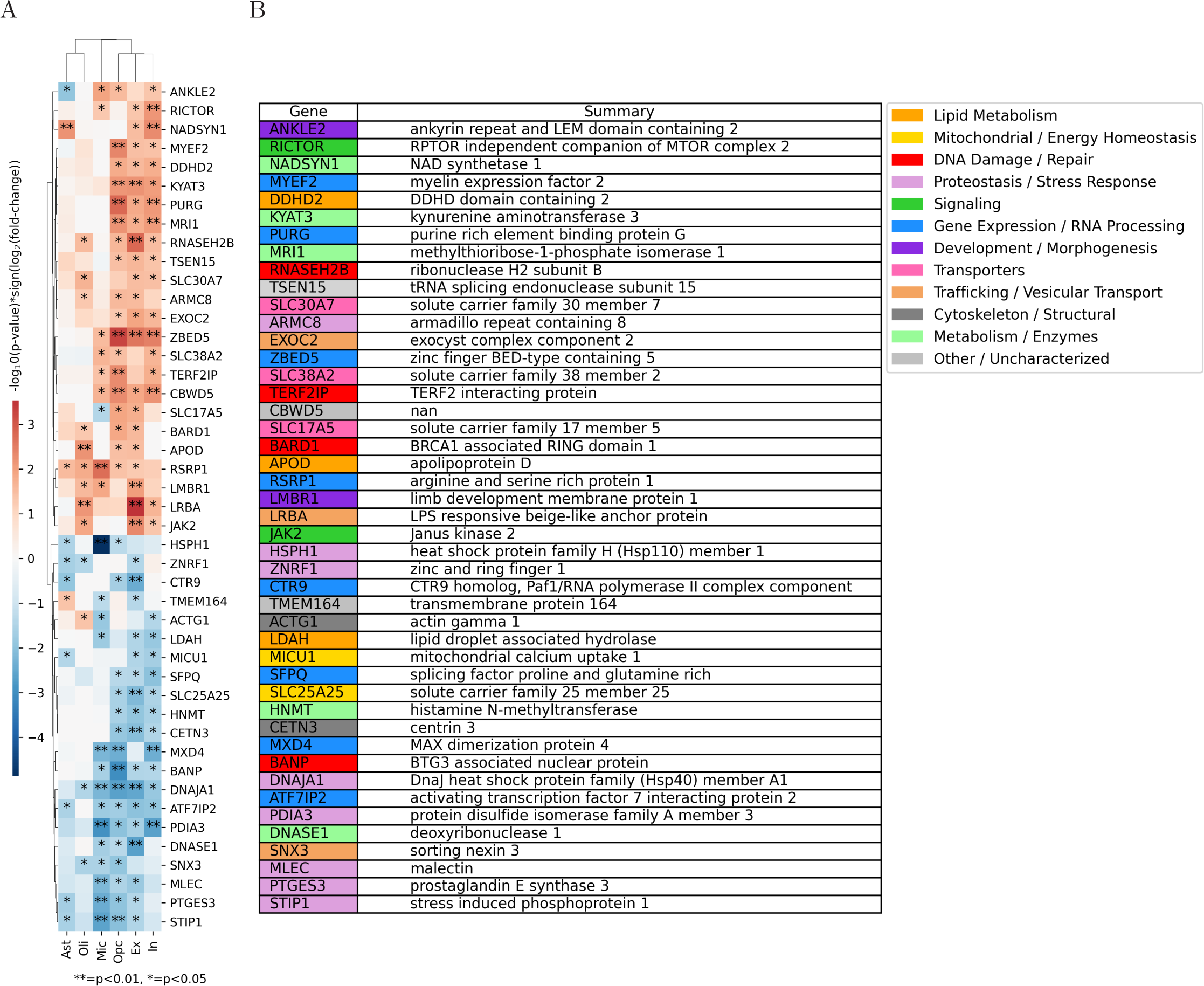
Shared Differentially Expressed Genes by Cell Type. **(A)** Heatmap indicating the overlap of differentially expressed genes between cell types (genes, where p-value < 0.05 in at least three cell types). Functional annotations of genes in the same order as in the heatmap in A.

**Figure S6:**
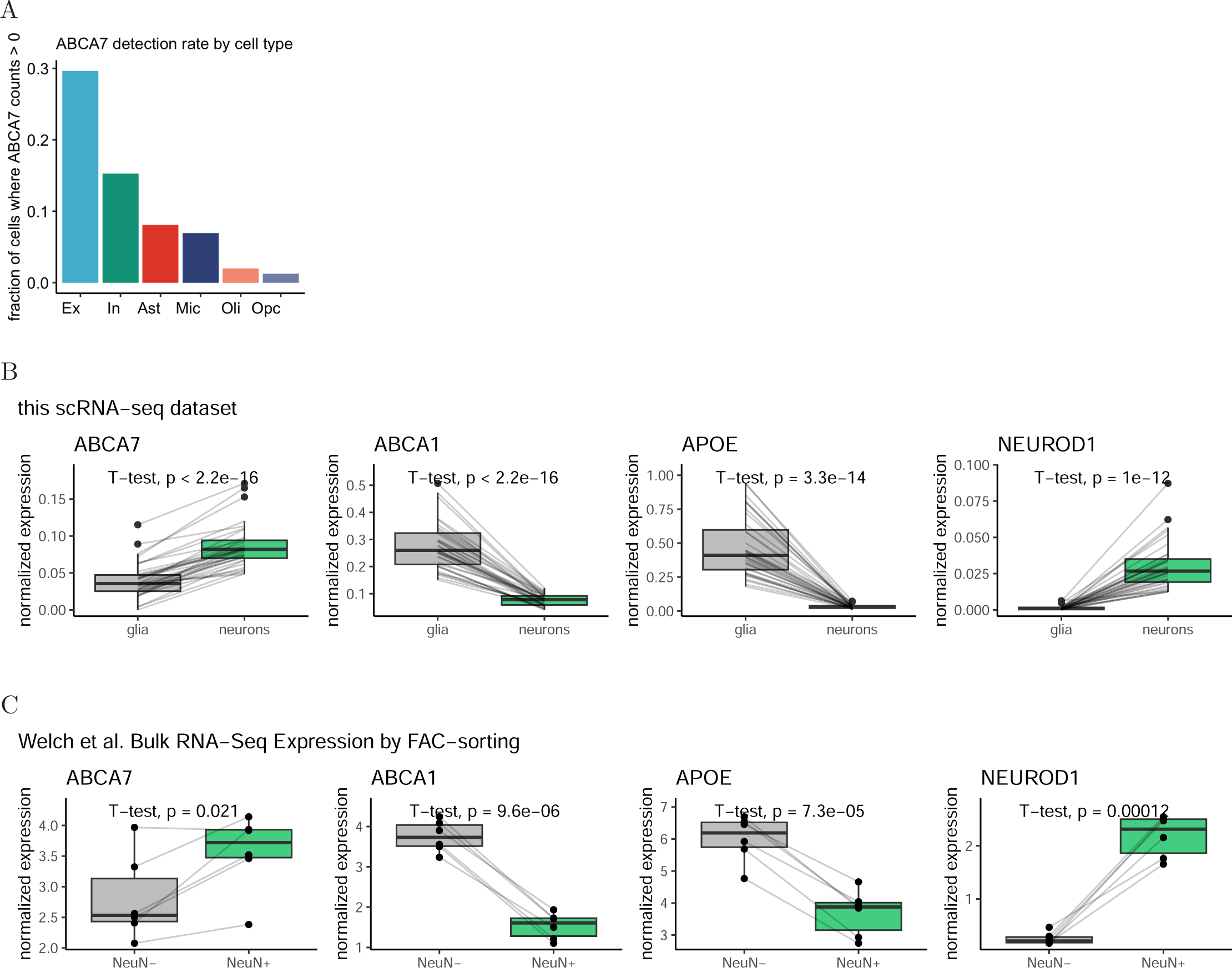
Neuronal Expression of ABCA7 in the Postmortem Human Brain. **(A)** Per cell type ABCA7 detection rate of major cell types in the *postmortem* PFC as quantified by snRNA-seq. **(B)** Normalized expression of indicated gene in glial cells (per-individual mean expression profiles across Oli, Opc, Ast, Mic) vs. neuronal cells (per-individual mean expression profiles across Ex and In) from *postmortem* snRNA-seq data. **(C)** Normalized expression of indicated genes in NeuN- vs. NeuN+ cells (N=6 individuals, from [51]; Table S6). All p-values are computed by paired two-sided t-test. Boxes indicate per-condition dataset quartiles, and whiskers extend to the most extreme data points not considered outliers (i.e., within 1.5 times the interquartile range from the first or third quartile).

**Figure S7:**
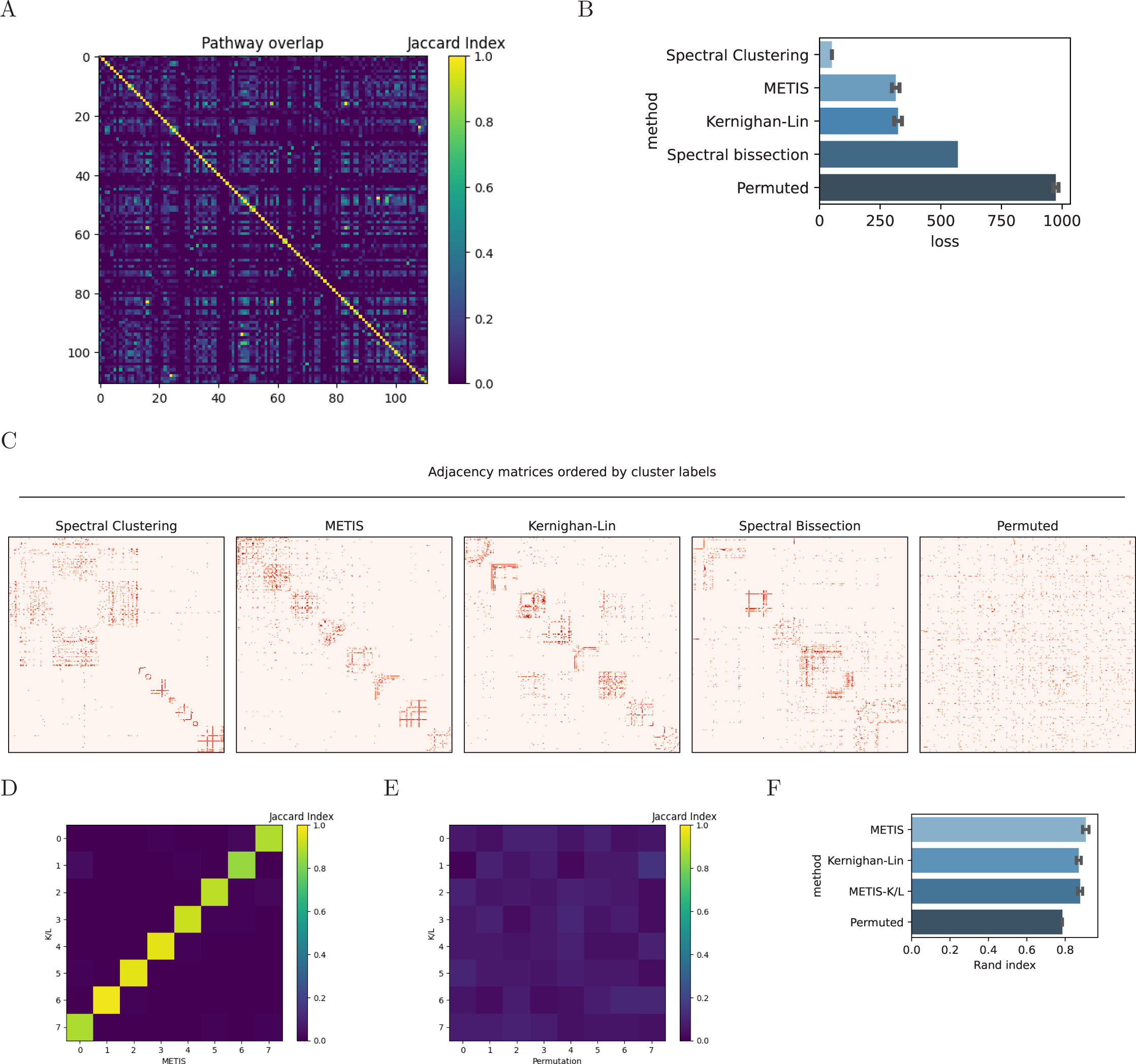
Benchmarking Partitioning and Clustering Algorithms for Gene-Pathway Grouping. **(A)** Jaccard indices quantifying overlap of genes for all 111 pathways in Figure 2B (see Methods; Supplementary Text). Consistency is quantified using the Jaccard Index (JI). JI = <u>^|^</u>*<u>^A^</u>*<u>^∩^</u>*<u>^B^</u>*<u>^|^</u> , where A and B are two sets (i.e., cluster A from initiation #1 and cluster B from initiation #2). **(B)** Average loss (total cut size; see Methods) associated with applying each algorithm (spectral clustering (SC), METIS, Kernighan-Lin (K/L), spectral bisection (SB), or random permutation) on the graph *G* (with 379 vertices; see Methods) over 1000 initiations (SC, random permutation) or 5 × 10^5^ initiations (METIS, K/L). The SB implementation is deterministic and was run only once. Error bars indicate the standard deviation. **(C)** Unweighted adjacency matrix for *G* sorted by labels assigned by the indicated algorithm. Red indicates the presence of an edge between two vertices. For each algorithm, labels corresponding to the best initiation (lowest loss) over 1000 initiations (SC, random permutation) or 5 × 10^5^ initiations (METIS, K/L) are shown. **(D)** Pairwise labeling consistency for the best K/L initiation and the best METIS initiation. Cluster labels corresponding to each are shown on the X- and Y-axes, respectively. Each color entry indicates the fraction of shared vertices per cluster across two initiations. **(E)** Same as (D), but comparing the best K/L initiation against the best random permutation initiation. Average Rand index (RI) for all pairwise initiations from (B). “METIS,” “Kernighan-Lin,” and “Permuted” labels on the Y-axis indicate the average RI (consistency across two sets of labels) for all combinations of initiations within the specified algorithm. “METIS-K/L” indicates the average RI for all combinations of initiations across the METIS and Kernighan-Lin algorithms. Error bars indicate standard deviations. (RI = number of agreeing vertex pairs ).

**Figure S8:**
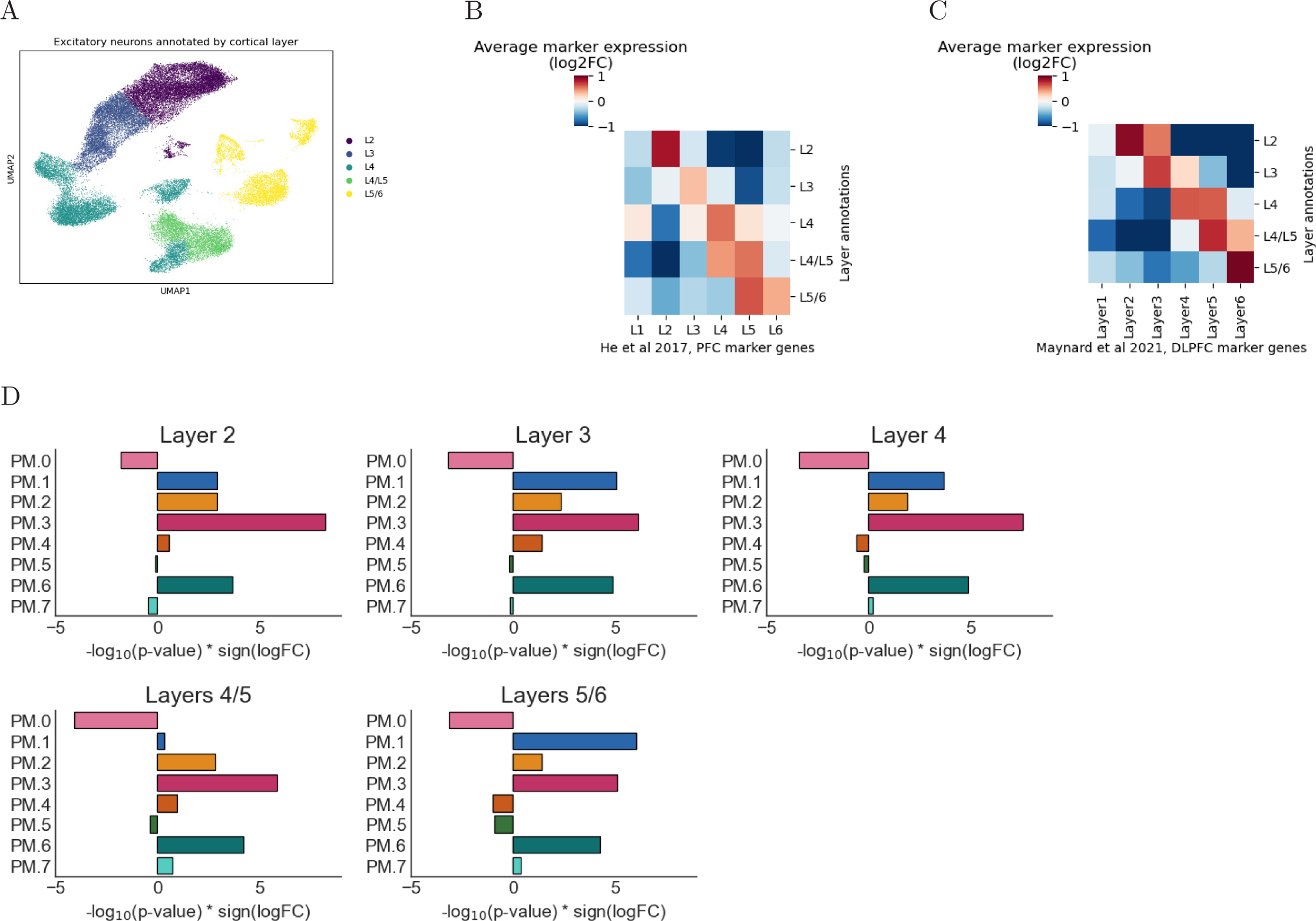
Annotation of Excitatory Neurons from *postmortem* snRNAseq Dataset by Cortical Layer. **(A)** UMAP visualization of excitatory neurons annotated by cortical layers based on Leiden clustering. **(B)** Heatmap showing enrichment of cortical layer-specific marker genes from [126] across annotated layers. Color indicates average marker gene expression (log2 fold change) for each layer marker gene set with respect to all other clusters. **(C)** Heatmap displaying validation of layer annotations using an independent set of cortical layer marker genes from [127]. Color represents average marker gene expression (log2 fold change) for each layer marker gene set with respect to all other clusters. ABCA7 LoF-associated perturbations of excitatory neuronal PM gene clusters by layer (computed by FGSEA, see Methods).

**Figure S9:**
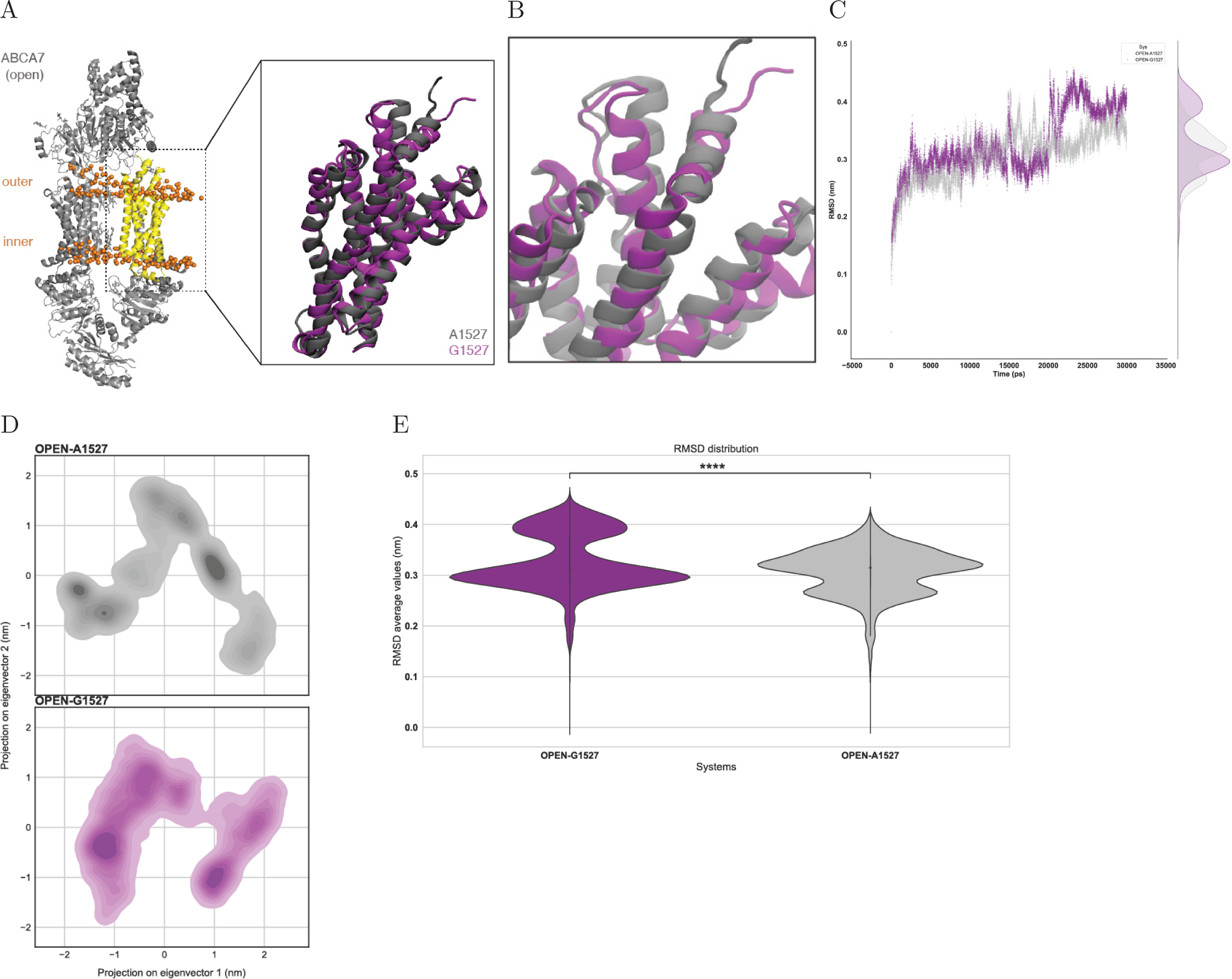
Molecular Dynamics Simulations of ABCA7 Open Conformations with p.Ala1527Gly Substitution. **(A)** Open conformation ABCA7 protein structure. ABCA7 domain between residues 1517 and 1756 used for simulations is shown in yellow. Expanded yellow domain (inset from left), with A1527 variant (light grey) and G1527 variant (purple). **(B)** Expanded inset from A. **(C)** Root mean squared deviations of open conformation domains from B with A1527 (light grey) or G1527 (purple) under simulation. Structural deviations over time were computed with respect to reference open structures from B. **(D)** Projection of *C_α_* atom positional fluctuations under simulation onto the first two principal components, for open conformation domain from B with A1527 (top, light grey) or G1527 (bottom, purple). Violin plot indicating average *C_α_* atom positional fluctuations over time.

**Figure S10:**
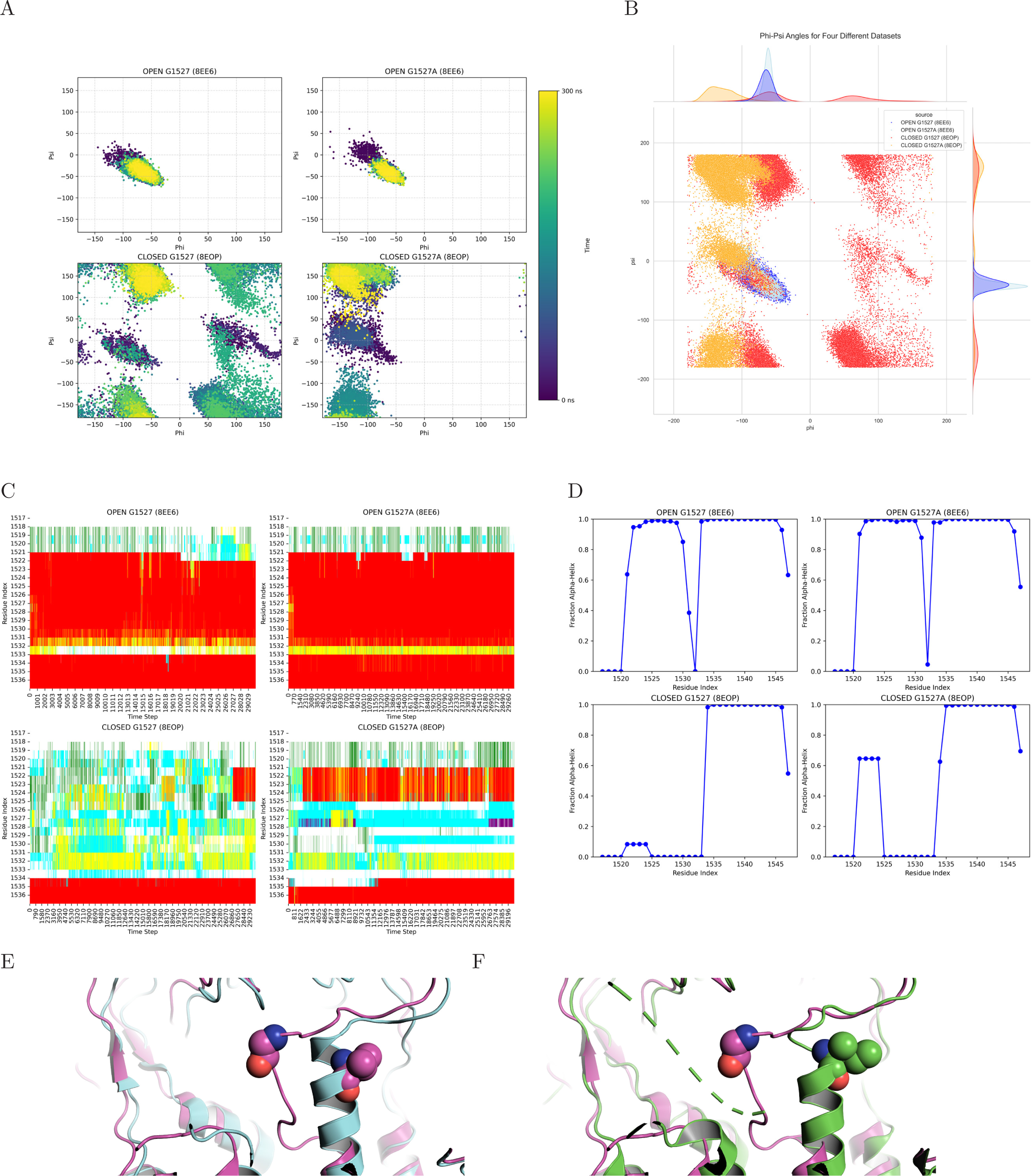
Analysis of Local Conformational Fluctuations and Secondary Structure Variations Induced by the p.Ala1527Gly Substitution in ABCA7 Open and Closed Conformations. **(A)** Phi vs. Psi dihedral angle distribution of residue 1527 as a function of simulation time for open and closed conformations. **(B)** Overall Phi vs. Psi dihedral angle distributions for residue 1527, comparing open and closed conformations throughout the entire simulation period. **(C)** Time-resolved secondary structure assignments for residues 1517–1537. Alpha-helical structures are highlighted in red; other colors represent distinct secondary structures. **(D)** Fraction of alpha-helical content observed for residues 1517–1537 during the simulations. A value of 1 indicates uninterrupted preservation of the alpha-helical structure throughout the simulation duration. **(E)** ABCA1 (cyan) closed structure; PDB ID: 7TBW. ABCA7 (purple) closed structure; PDB ID: 8EOP. Positions of Gly1527 in ABCA7 and V1646 in ABCA1 are indicated as spheres. **(F)** ABCA4 (green) closed structure; PDB ID: 7LKZ. ABCA7 (purple) closed structure; PDB ID: 8EOP. Positions of Gly1527 in ABCA7 and I1671 in ABCA4 are indicated as spheres.

**Figure S11:**
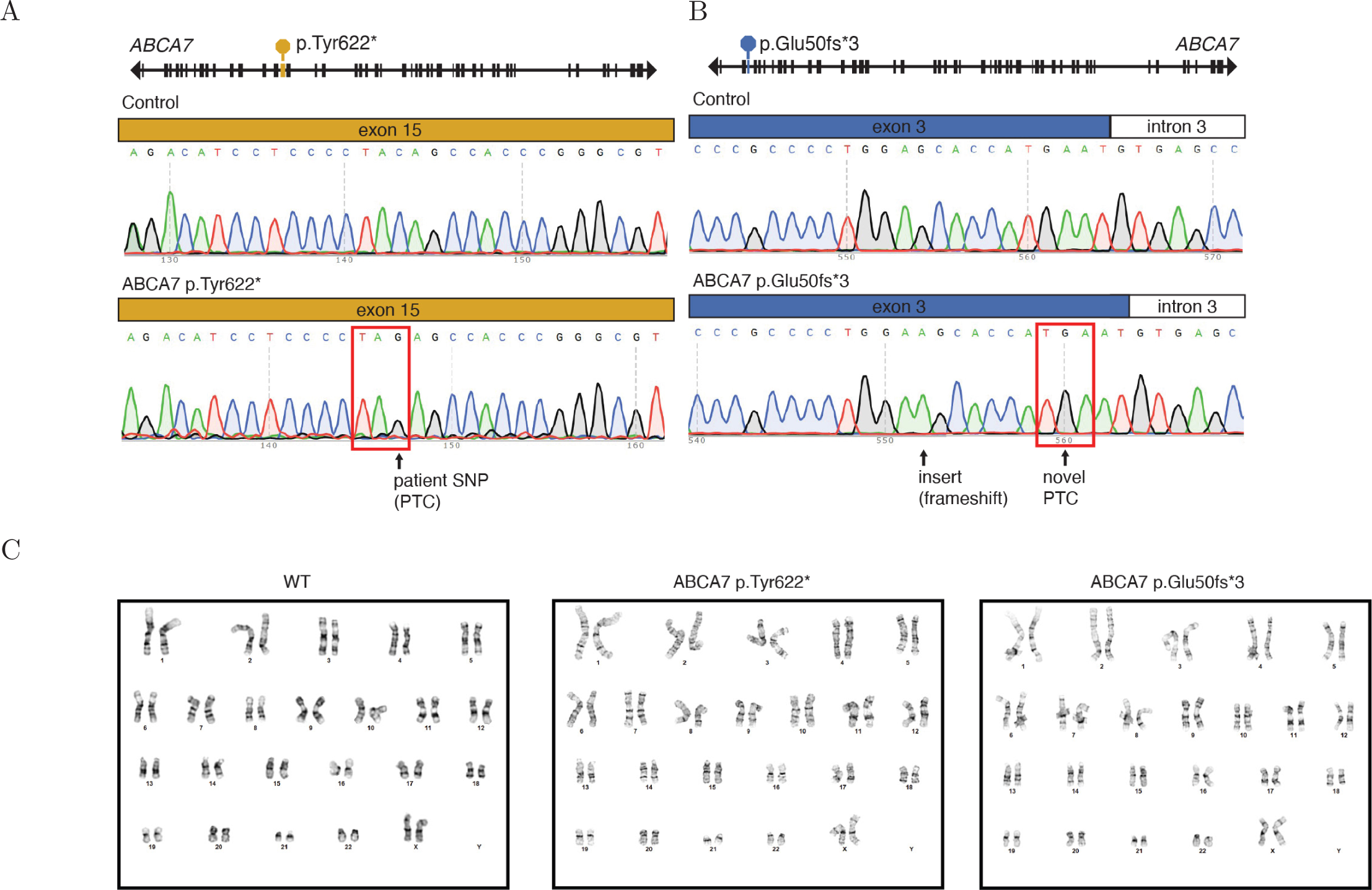
Generation of iPSC-Derived Cells Harboring ABCA7 PTC Variants. **(A)** Sanger sequencing chromatogram confirming single nucleotide insertion in ABCA7 exon 3 of the ABCA7 p.Glu50fs*3 isogenic iPSC line. **(B)** Sanger sequencing chromatogram confirming patient single nucleotide polymorphism in ABCA7 exon 15 of the ABCA7 p.Tyr622* isogenic iPSC line. **(C)** Normal karyotypes were observed for WT, ABCA7 p.Glu50fs*3, and ABCA7 p.Tyr622* isogenic iPSC lines.

**Figure S12:**
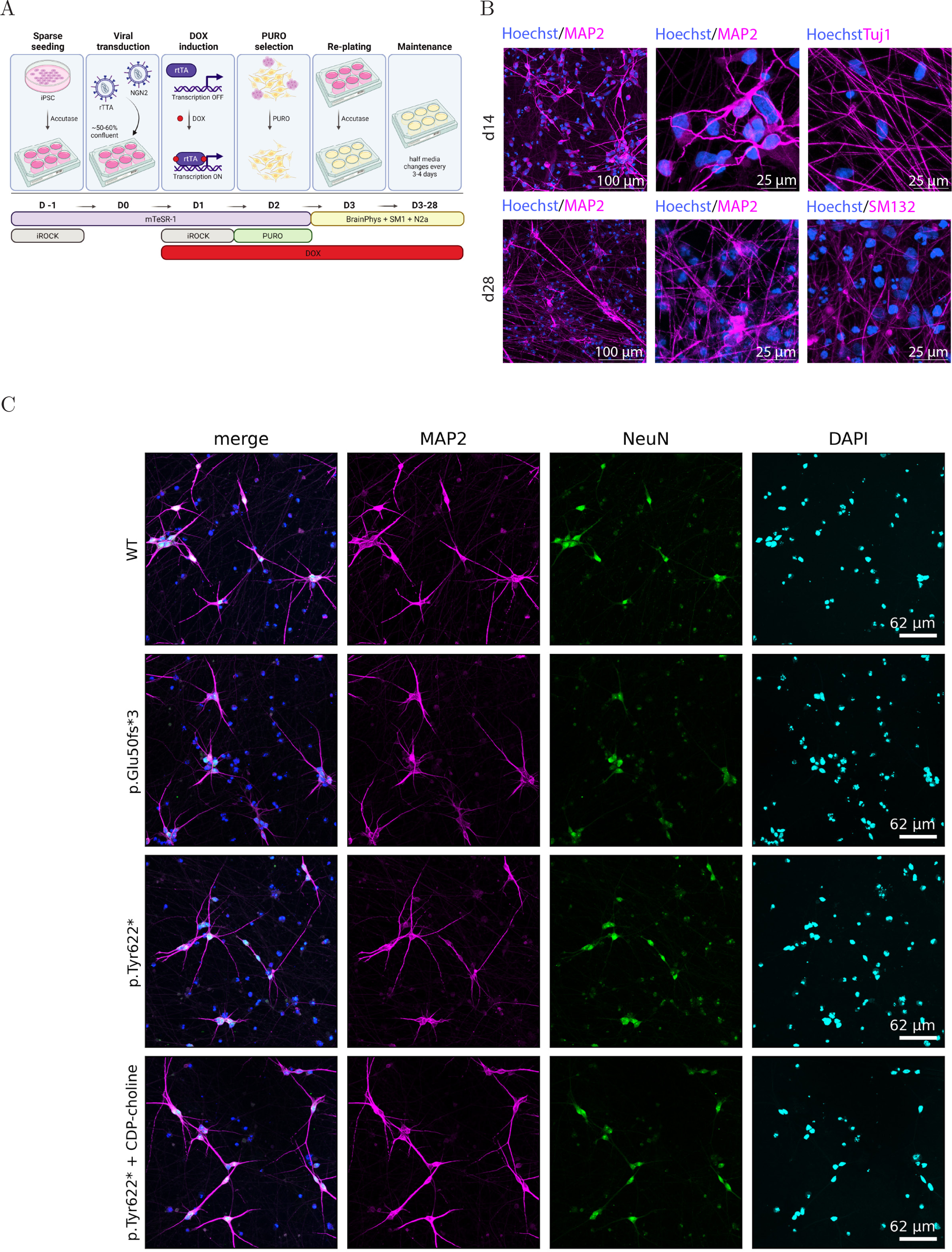
Differentiating iPSC-Derived Neurons Harboring ABCA7 PTC Variants. **(A)** iPSCs were plated at low density for NGN2 viral transduction. Expression of NGN2 was driven by doxycycline (DOX) induction with puromycin (PURO) selection, then cells were replated to match neuronal densities. Neurons were maintained for 4 weeks (DIV 28) before experimentation (Created with BioRender.com). **(B)** Neuronal marker expression in 2 and 4-week matured iNs. Neuronal marker expression in iNs matured for 4 weeks for indicated genotypes. CDP-choline treatment was applied for 2 weeks at 100 *µ*M.

**Figure S13:**
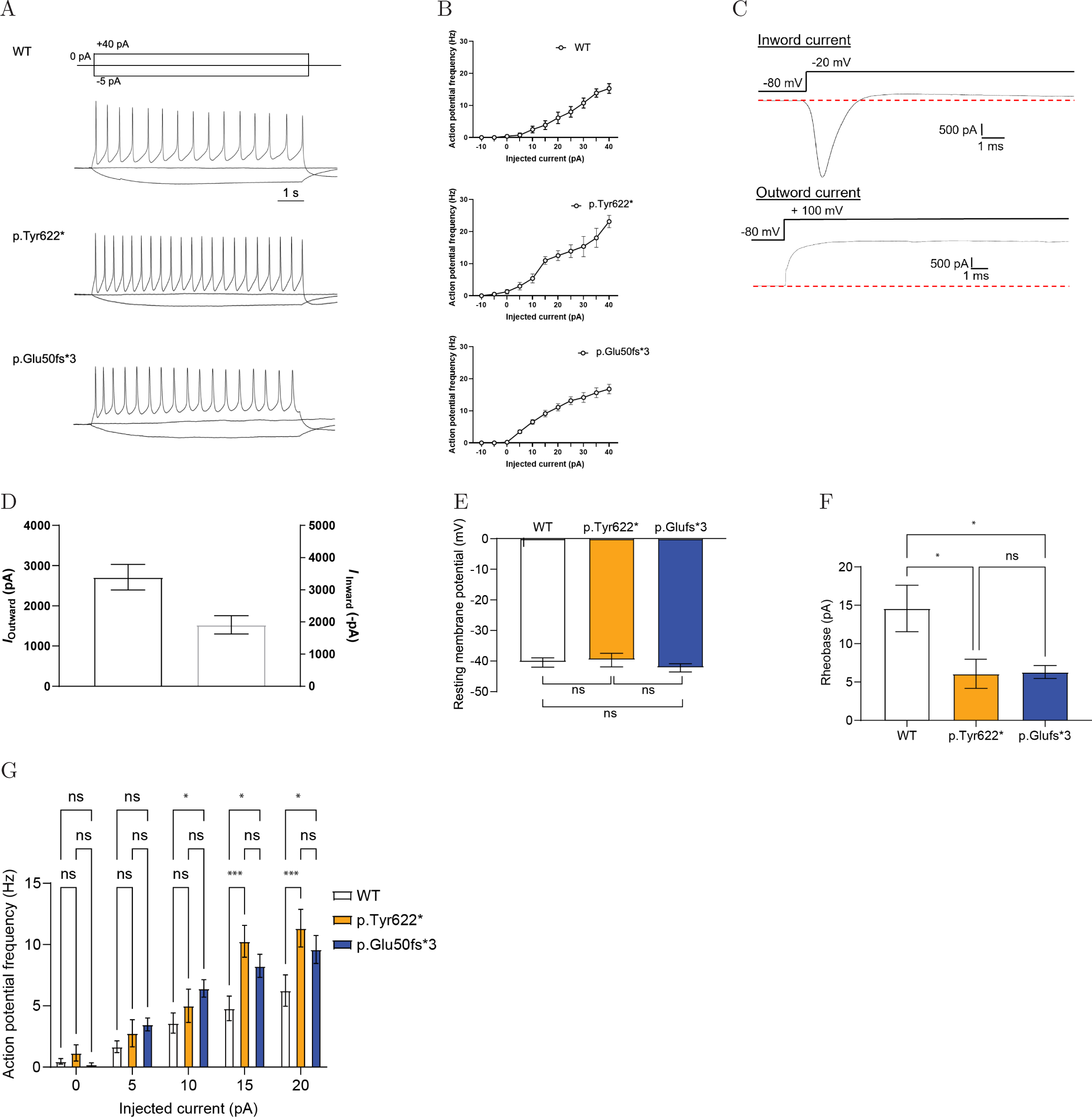
Measuring activity of iPSC-derived neurons harboring ABCA7 PTC variants. **(A)** Representative sweeps show action potentials elicited by 800 ms of current injections in patched 4-week-old iNs. **(B)** Summary of action potential frequency (means ± SEM) elicited with different amounts of injected current in 4-week-old iNs. **(C)** Representative sweeps of whole-cell current flow of inward (upper panel) and outward (lower panel) current recordings from WT 4-week-old neurons. **(D)** Quantification of (C). **(E)** Resting membrane potential (mV) of 4-week-old WT, ABCA7 p.Tyr622*, and ABCA7 p.Glufs*3 neurons. **(F)** Rheobase (pA) of 4-week-old WT, ABCA7 p.Tyr622*, and ABCA7 p.Glufs*3 neurons. **(G)** Action potential frequency of 4-week-old WT, ABCA7 p.Tyr622*, and ABCA7 p.Glufs*3 neurons with indicated current injections. For panels E-G: WT: *n* = 24; Y622: *n* = 13; G2: *n* = 23. For all panels: *P* ∗ *<* 0.05, *P* ∗ ∗∗ *<* 0.001. Graphs are mean ± SEM.

**Figure S14:**
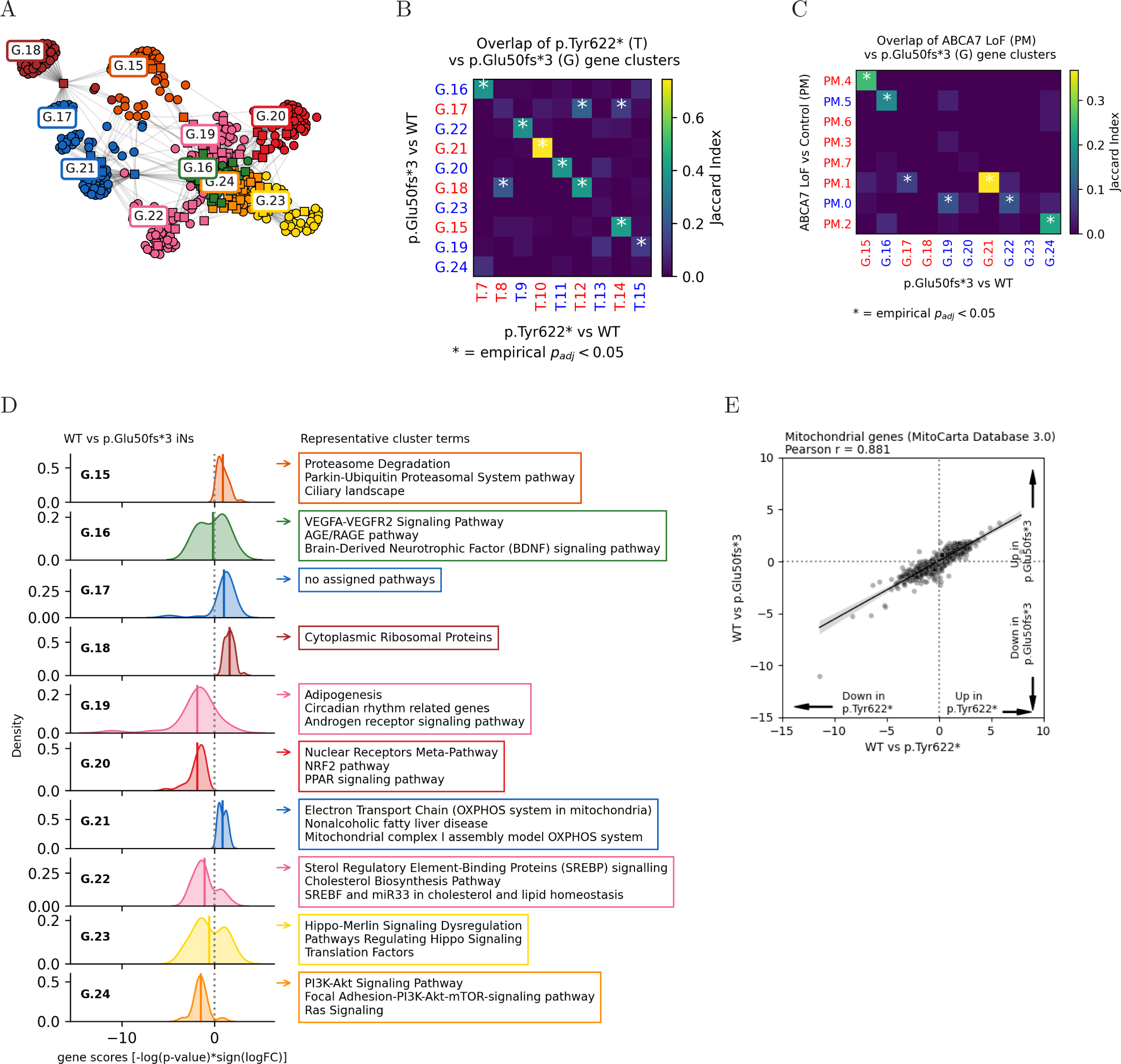
mRNA-seq analysis of p.Glu50fs* vs. WT iNs. (A) Kernighan-Lin (K/L) clustering of leading-edge genes from significantly perturbed pathways (Ben- jamini–Hochberg (BH) FDR-adjusted *p <* 0.05) in p.Glu50fs*3 vs. WT iNs. Colors represent distinct K/L clusters. **(B)** Heatmap of Jaccard index overlap between K/L gene clusters from p.Glu50fs*3 neurons and clusters identified in p.Tyr622* vs WT iNs. Red text denotes clusters with average score *S* upregulated in ABCA7 LoF; blue text denotes clusters with average *S* downregulated in ABCA7 LoF. **(B)** Heatmap showing Jaccard index overlap between K/L clusters identified in p.Glu50fs* vs. WT iNs and p.Tyr622* vs. WT iNs. **(C)** Heatmap of Jaccard index overlap between K/L gene clusters from p.Glu50fs*3 neurons and clusters identified in human postmortem excitatory neurons. **(D)** Gaussian kernel density plots of gene perturbation scores (*S*) within each cluster. Positive *S* indicates upregulation in p.Glu50fs*3. Solid lines denote cluster means. Top enriched pathways with highest intra-cluster connectivity indicated. **(E)** Correlation of per-gene perturbation scores (*S* = − log_10_(p-value) × sign(log_2_(fold change))) between p.Glu50fs*3 vs. WT and p.Tyr622* vs. WT iNs for MitoCarta genes.

**Figure S15:**
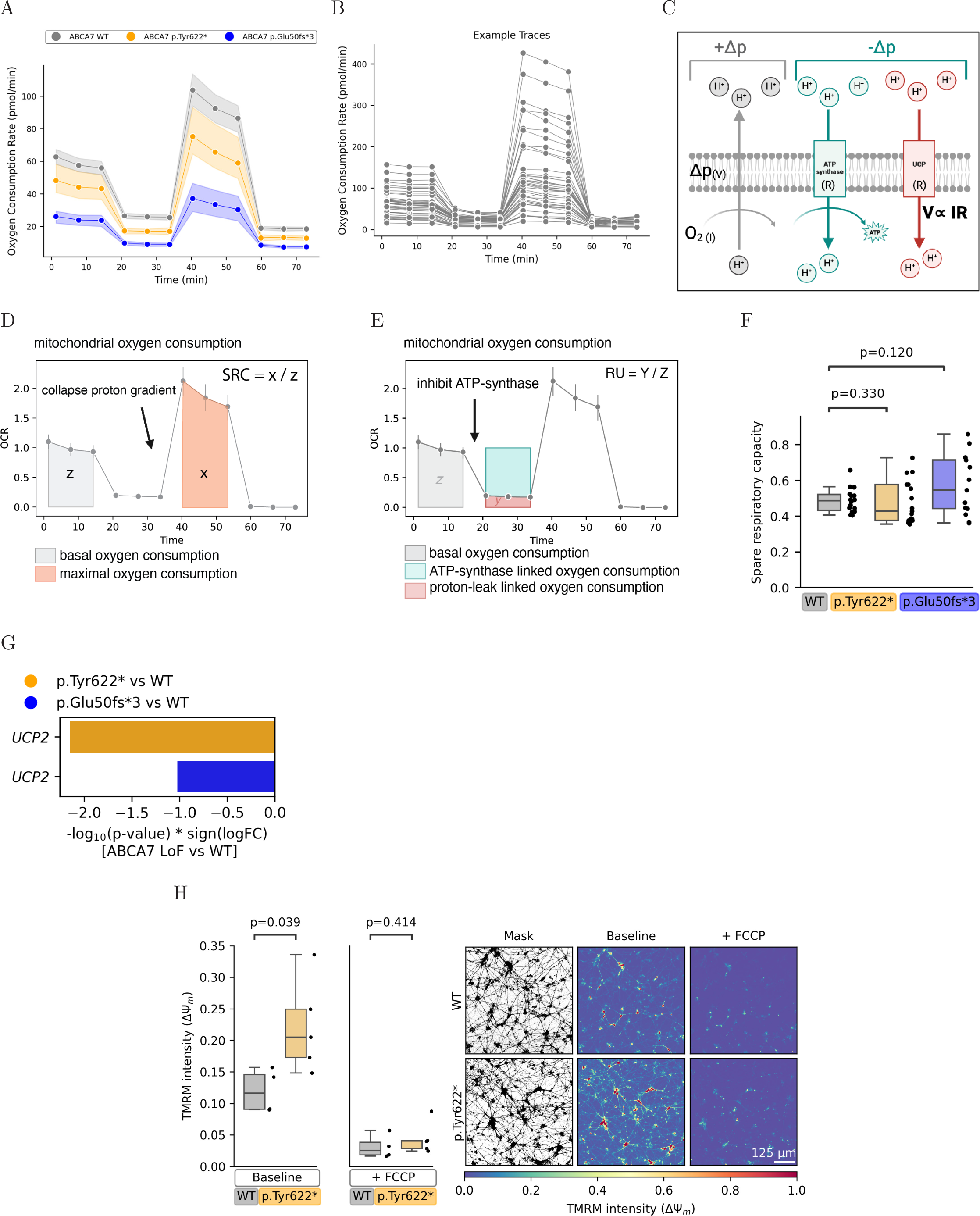
Analysis of Oxygen Consumption Rates in ABCA7 LoF vs. Control iNs. **(A)** Example oxygen consumption rate (OCR) curves from Batch 1 of the two differentiation batches used for analysis in Figure 3. The line plot indicates the per-condition mean estimator, and the error bars indicate the 95% confidence interval. **(B)** Representative per-well traces from (A). **(C)** Schematic indicating the relationship between oxygen consumption as a measure of proton current (I), which sustains the proton motive force (voltage, V). Regulation of ATP synthase and uncoupling protein (UCP) activity modifies resistance (R) and depletes the proton motive force. **(D)** Schematic indicating measurement of maximal and basal oxygen consumption to compute SRC. **(E)** Schematic indicating measurement of uncoupled oxygen consumption. **(F)** SRC computed for WT, ABCA7 p.Glu50fs*3, and ABCA7 p.Tyr622* iNs. P-values computed by independent sample t-test. *N* wells = 18 (WT), 17 (p.Tyr622*), 13 (p.Glu50fs*3) across two independent differentiation batches and Seahorse experiments. **(G)** UCP2 mRNA levels by genotype. **(H)** Mitochondrial membrane potential quantified by average TMRM fluorescence intensity per masked region (thresholded at 75th percentile) under baseline conditions and after addition of FCCP in ABCA7 LoF and WT iNs cultured for 4 weeks. Each datapoint represents average intensity per well. *N* = 4 (WT), 5 (p.Tyr622*) wells. Statistical comparison by independent-sample *t*-test. Same plot and images as shown under baseline conditions in Figure 3.

**Figure S16:**
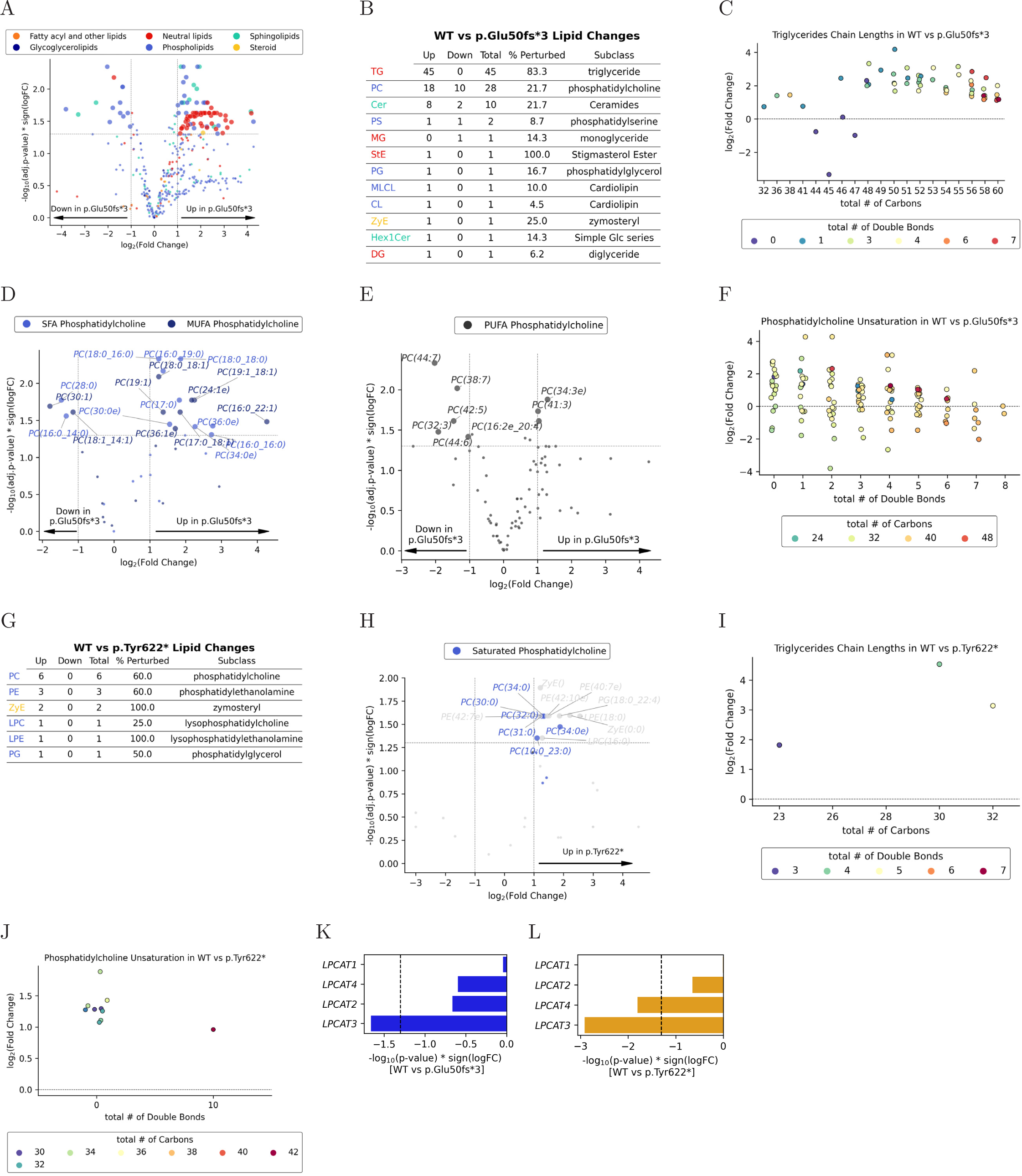
LC-MS Lipidomics in ABCA7 LoF iNs. (A) Volcano plot of significantly perturbed lipid species identified by LC-MS in p.Glu50fs*3 vs. WT iNs (BH FDR-adjusted *p <* 0.05, | log_2_(FC)| *>* 1), colored by lipid class. *N* = 6 wells per genotype. **(B)** Significantly perturbed lipid species from (A), summarized by lipid subclass. **(C)** Distribution of triglyceride fold changes grouped by fatty acid chain length and saturation for p.Glu50fs*3 vs. WT iNs. **(D)** Volcano plot highlighting significantly perturbed phosphatidylcholine species containing saturated or monounsaturated fatty acids (SFA/MUFA; BH FDR-adjusted *p <* 0.05, | log_2_(FC)| *>* 1) for p.Glu50fs*3 vs. WT iNs. **(E)** Volcano plot highlighting significantly perturbed phosphatidylcholine species containing polyunsaturated fatty acids (PUFA; BH FDR-adjusted *p <* 0.05, | log_2_(FC)| *>* 1) for p.Glu50fs*3 vs. WT iNs. **(F)** Distribution of phosphatidylcholine fold changes grouped by fatty acid chain length and saturation for p.Glu50fs*3 vs. WT iNs. **(G)** Table summarizing significantly perturbed lipid species by lipid subclass in p.Tyr622* vs. WT iNs (*N* = 10 WT, 8 p.Tyr622* wells; BH FDR-adjusted *p <* 0.05, | log_2_(FC)| *>* 1). **(H)** Volcano plot showing lipid species from (G), colored by lipid class; phosphatidylcholines highlighted in blue, for p.Tyr622* vs. WT iNs. **(I,J)** Same analysis as (C,F), but comparing p.Tyr622* vs. WT iNs. **(K,L)** Expression changes (mRNA) of LPCAT genes comparing p.Tyr622* vs. WT and p.Glu50fs*3 vs. WT iNs.

**Figure S17:**
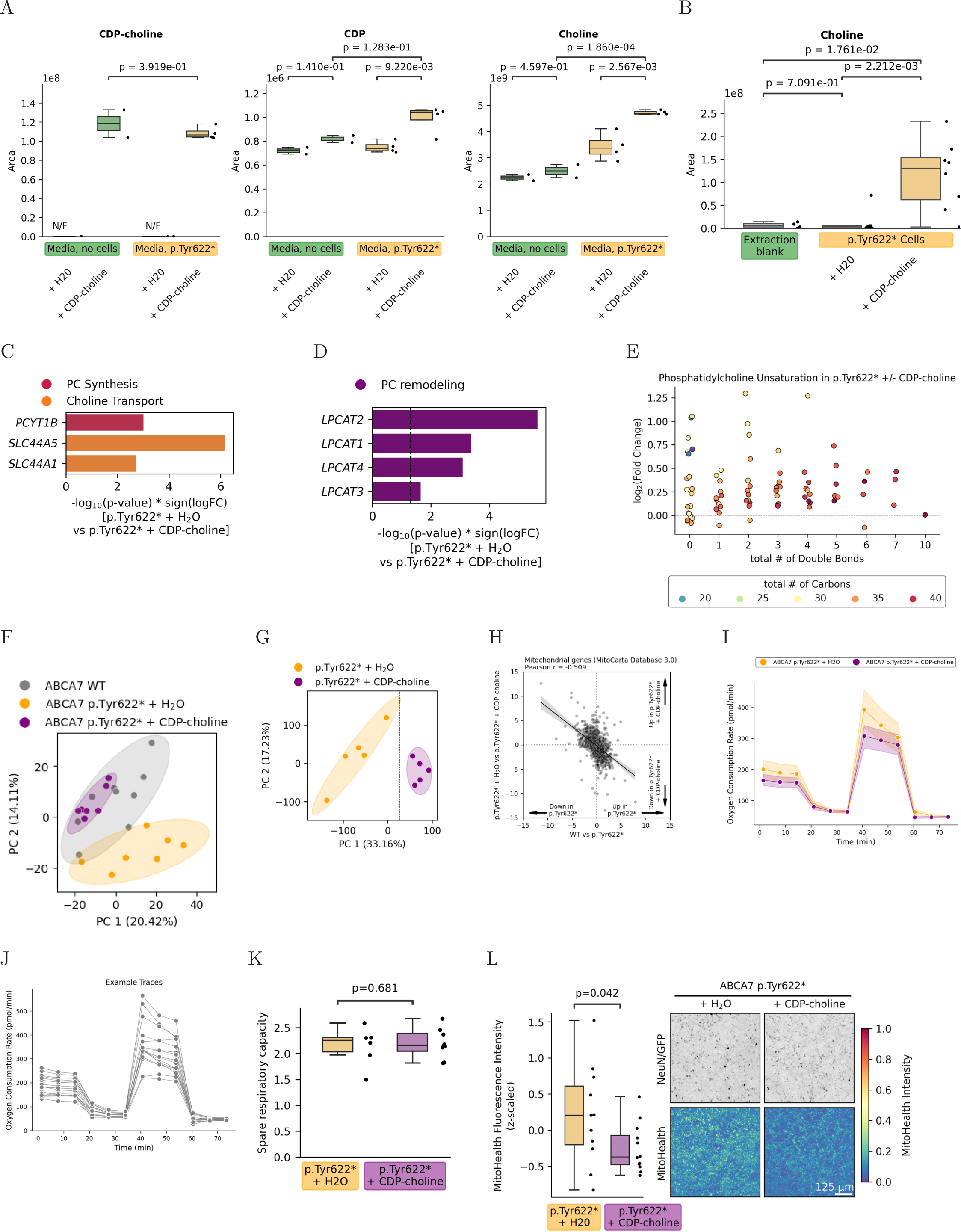
Effects of CDP-choline Treatment in p.Tyr622* iNs. **(A)** Choline metabolites detected in media by targeted LC-MS. *N* = 2 for media without cells, *N* = 4 for cell-conditioned media; N/F indicates not detected. **(B)** Choline metabolites detected intracellularly by targeted LC-MS. *N* = 8 per genotype, *N* = 4 blanks. **(C)** Selected choline synthesis and transport genes differentially expressed in p.Tyr622* ± CDP-choline iNs (mRNA-seq). **(D)** LPCAT gene expression changes in p.Tyr622* ± CDP-choline iNs (mRNA-seq). **(E)** Distribution of phosphatidylcholine species fold-changes by fatty acid chain length and saturation in p.Tyr622* ± CDP-choline iNs. **(F)** PCA plot of untargeted LC-MS metabolite profiles from p.Tyr622* ± CDP-choline and WT iNs. **(G)** PCA plot of mRNA-seq data from p.Tyr622* ± CDP-choline iNs. **(H)** Correlation of gene perturbation scores (*S*) for mitochondrial-localized genes comparing p.Tyr622* ± CDP-choline iNs versus p.Glu50fs*3 vs. WT iNs. **(I)** Example Seahorse oxygen consumption rate (OCR) curves. Lines represent per-condition means; error bars indicate 95% confidence intervals. **(J)** Representative per-well OCR traces from (I). **(K)** Quantification of spare respiratory capacity (SRC) from OCR curves in (I). Statistical comparisons by independent-sample *t*-tests; *N* = 6 wells (p.Tyr622* + H_2_O), *N* = 8 wells (p.Tyr622* + CDP-choline). Boxes indicate quartiles; whiskers extend to data within 1.5×IQR from quartiles. Quantification of mitochondrial membrane potential via neuronal HCS MitoHealth dye fluorescence intensity. Statistical comparisons by linear mixed-effects model on per-NeuN+ volume averages, with well-of-origin as random effect. *N* = 11 wells (p.Tyr622*), *N* = 9 wells (p.Glu50fs*3), *N* = 8 wells (WT); ∼ 3000 cells per condition from three differentiation batches. Individual data points represent per-well averages. Right: Representative images shown as mean-intensity projections of NeuN+ staining and corresponding MitoHealth and Hoechst signals within quantified NeuN+ volumes. Each NeuN/GFP image intensity was scaled relative to its maximum value, followed by gamma correction (*γ* = 0.5) for visualization.

**Figure S18:**
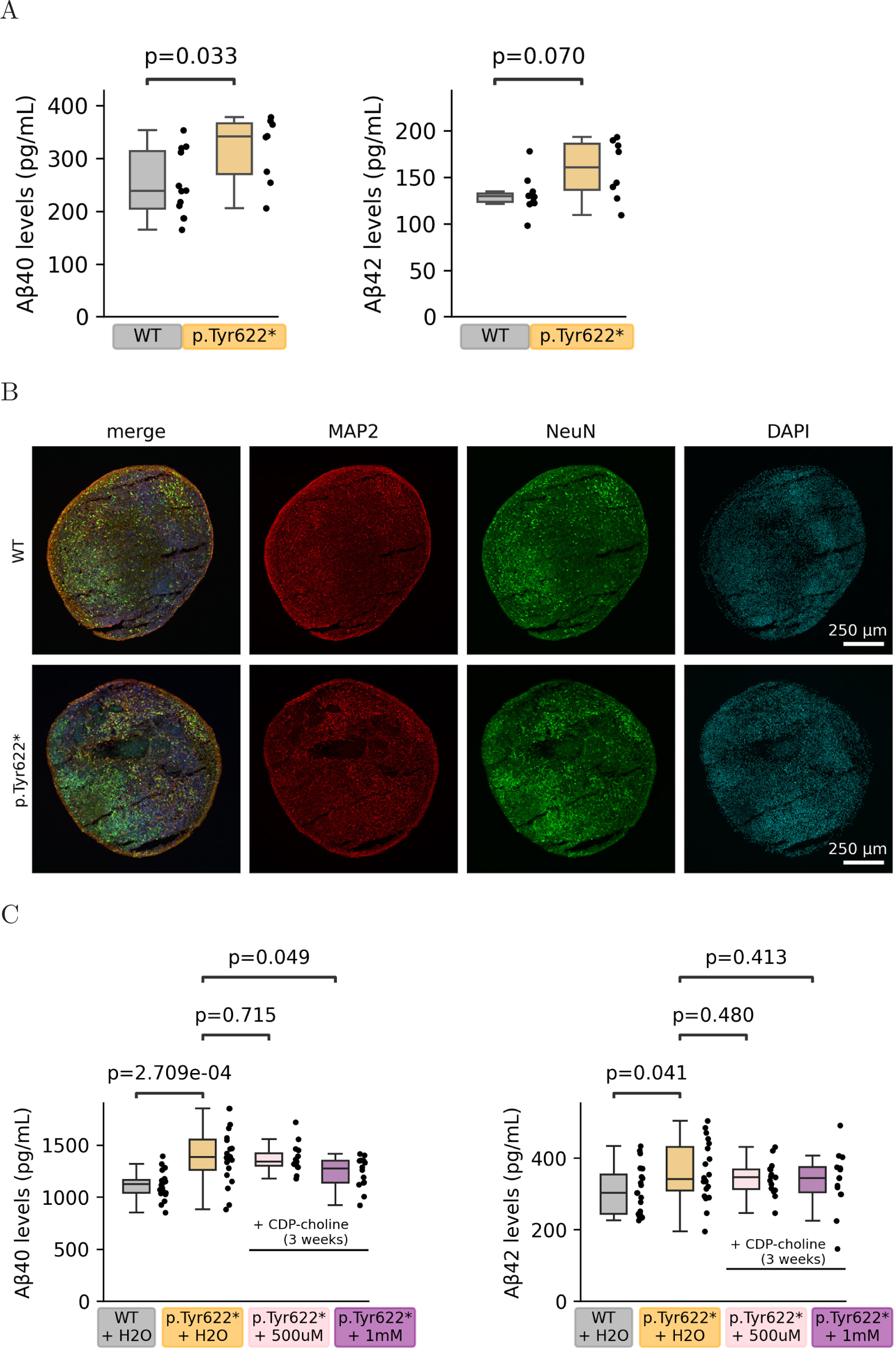
CDP-choline Treatment in Cortical Organoids. (A) Amyloid-*β* levels quantified by ELISA from media of 4-week-old iNs. **(B)** Representative images of cortical organoid slices from indicated genotypes. Amyloid-*β* levels quantified by ELISA from media of cortical organoids (176 days old), grouped by genotype and treated with 500 *µ*M or 1 mM CDP-choline for 3 weeks. Samples correspond to organoids in Figure 4K, analyzed one week prior to assays presented there.

## Supplementary Tables

**Table S1:** Annotation of ABCA7 loss of function variants used in this study.

| rsID | HGVS.c | HGVS.p | Annotation | AD associ-<br>ation | N in<br>co-<br>hort |
| --- | --- | --- | --- | --- | --- |
| rs113809142 | c.4416+2T>G | NA | splice donor<br>variant [130] | Steinberg et<br>al (2015),<br>Nature Ge-<br>netics, Table<br>1 [131] | 1 |
| rs200538373 | c.5570+5G>C | NA | splice region<br>variant [130,<br>131] | Steinberg et<br>al (2015),<br>Nature Ge-<br>netics, Table<br>1 [131] | 4 |
| rs538591288 | c.4208delT | p.Leu1403fs | frameshift<br>variant [130] | Steinberg et<br>al (2015),<br>Nature Ge-<br>netics, Table<br>1 [131] | 1 |
| rs547447016 | c.2126_2132delAGCAGGG | p.Glu709fs | frameshift<br>variant [130] | Steinberg et<br>al (2015),<br>Nature Ge-<br>netics, Table<br>1 [131] | 4 |
| rs201060968 | c.3641G>A | p.Trp1214* | stop gained | NA | 1 |
| 19_1053362_G_A | c.3255G>A | p.Trp1085* | stop gained | NA | 1 |

**Table S2:**
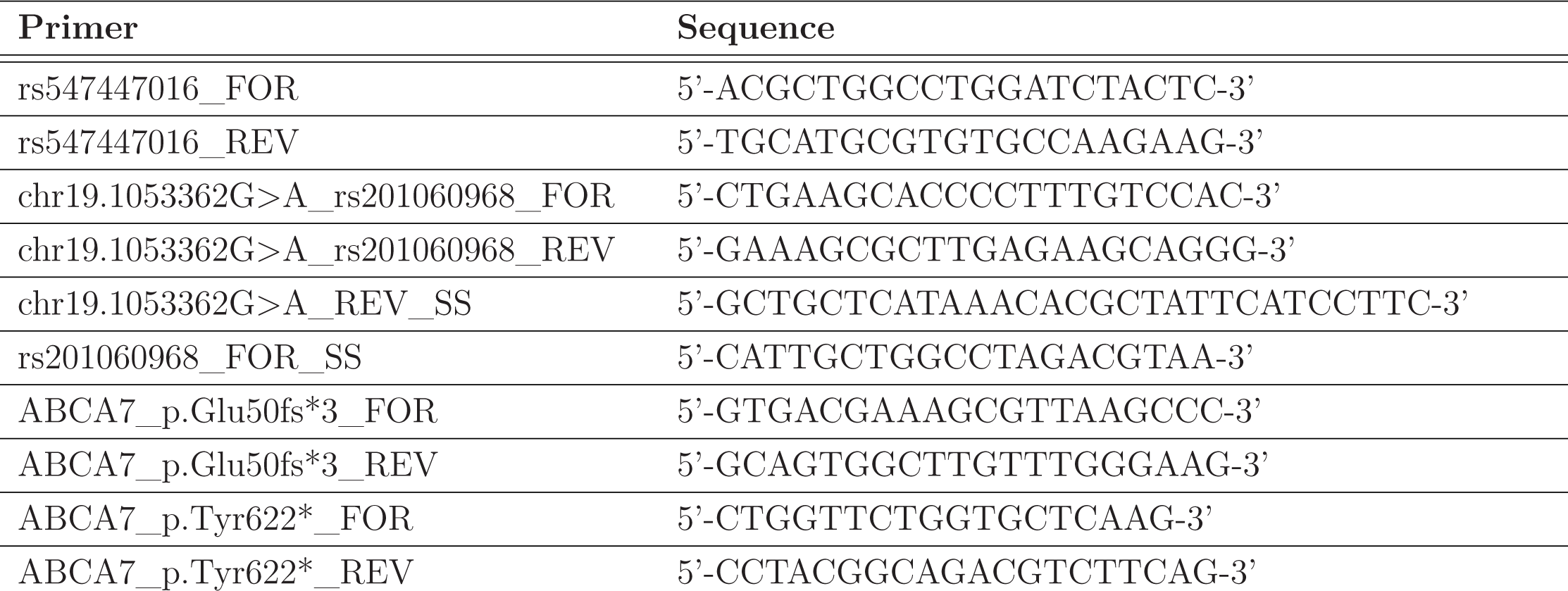
PCR/Sanger sequencing (SS) primers.

**Table S3:** Experimentally-determined 3D ABCA7 structures used in molecular dynamics simulations.

| **System** | **PDB ID** | **State** |
| --- | --- | --- |
| CLOSE-G1527 | 8EOP | HOLO |
| CLOSE-A1527 | 8EOP | HOLO |
| OPEN-G1527 | 8EE6 | APO |
| OPEN-A1527 | 8EE6 | APO |

**Table S4:**
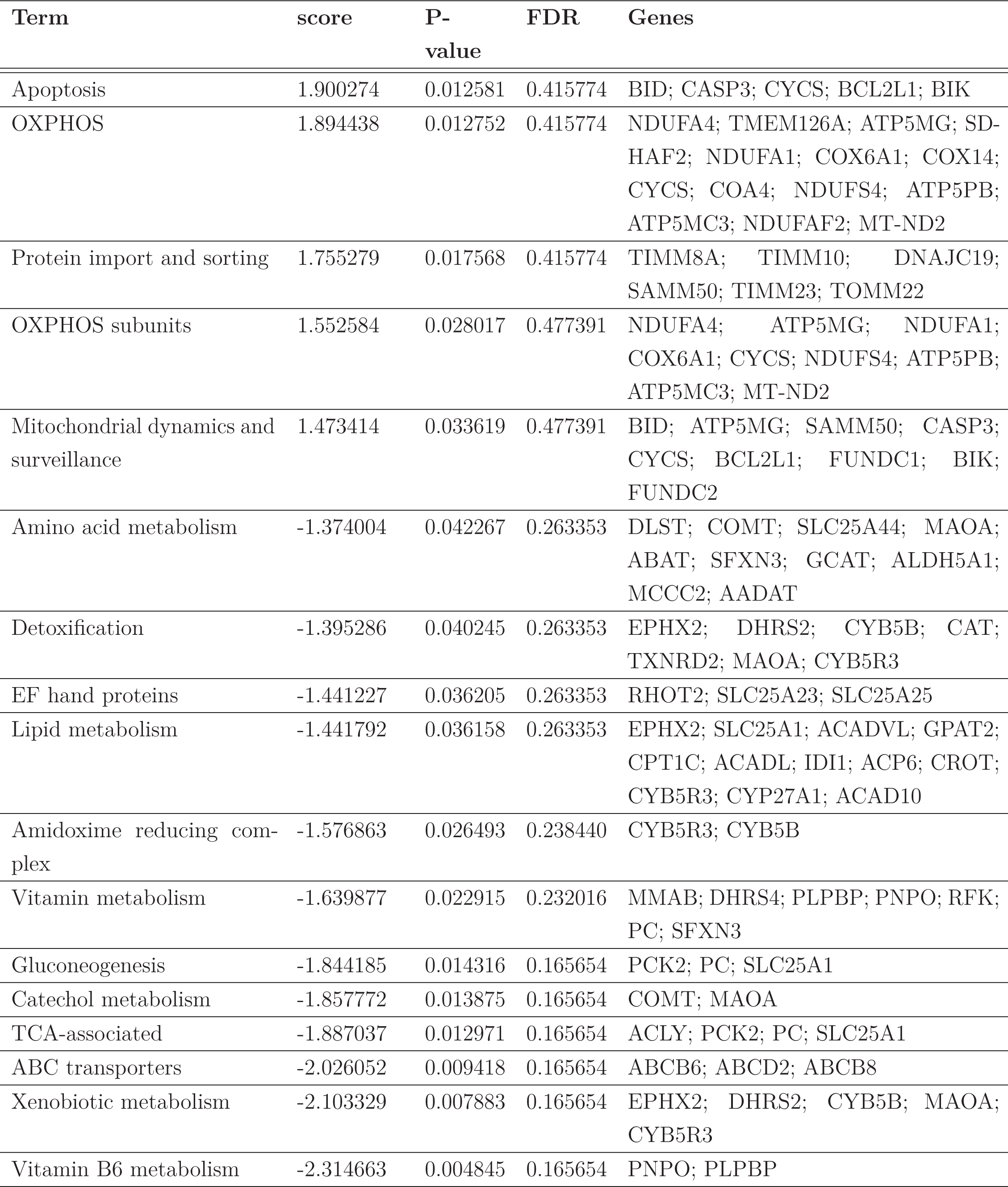
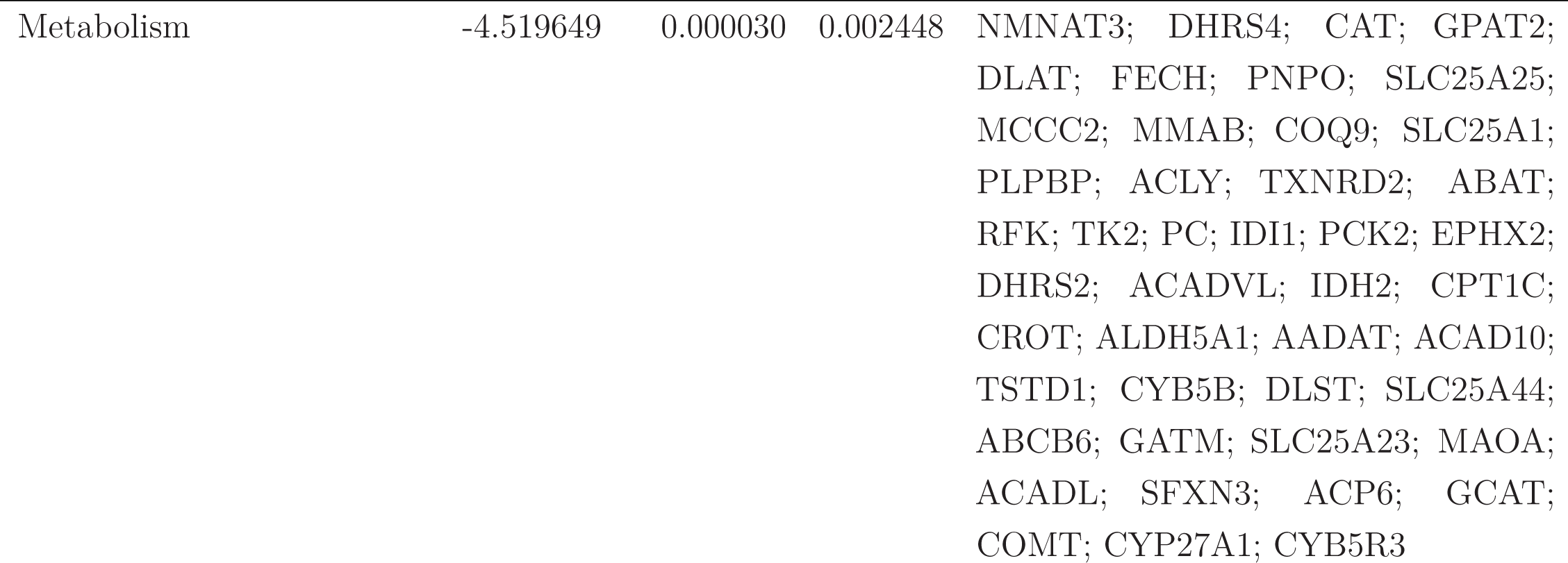
Enrichments of MitoCarta mitochondrial pathways in WT vs p.Tyr622* mRNA.

**Table S5:**
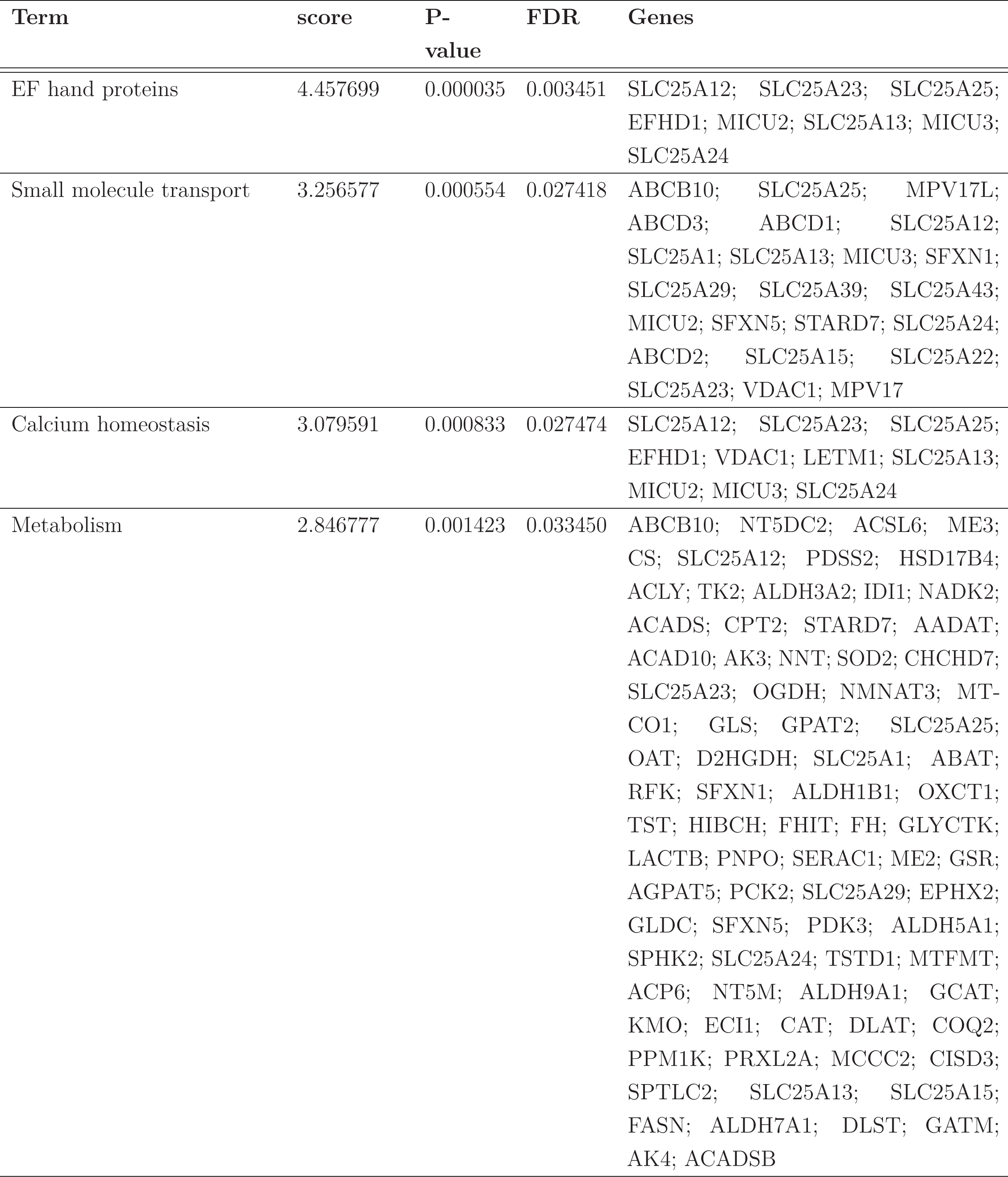
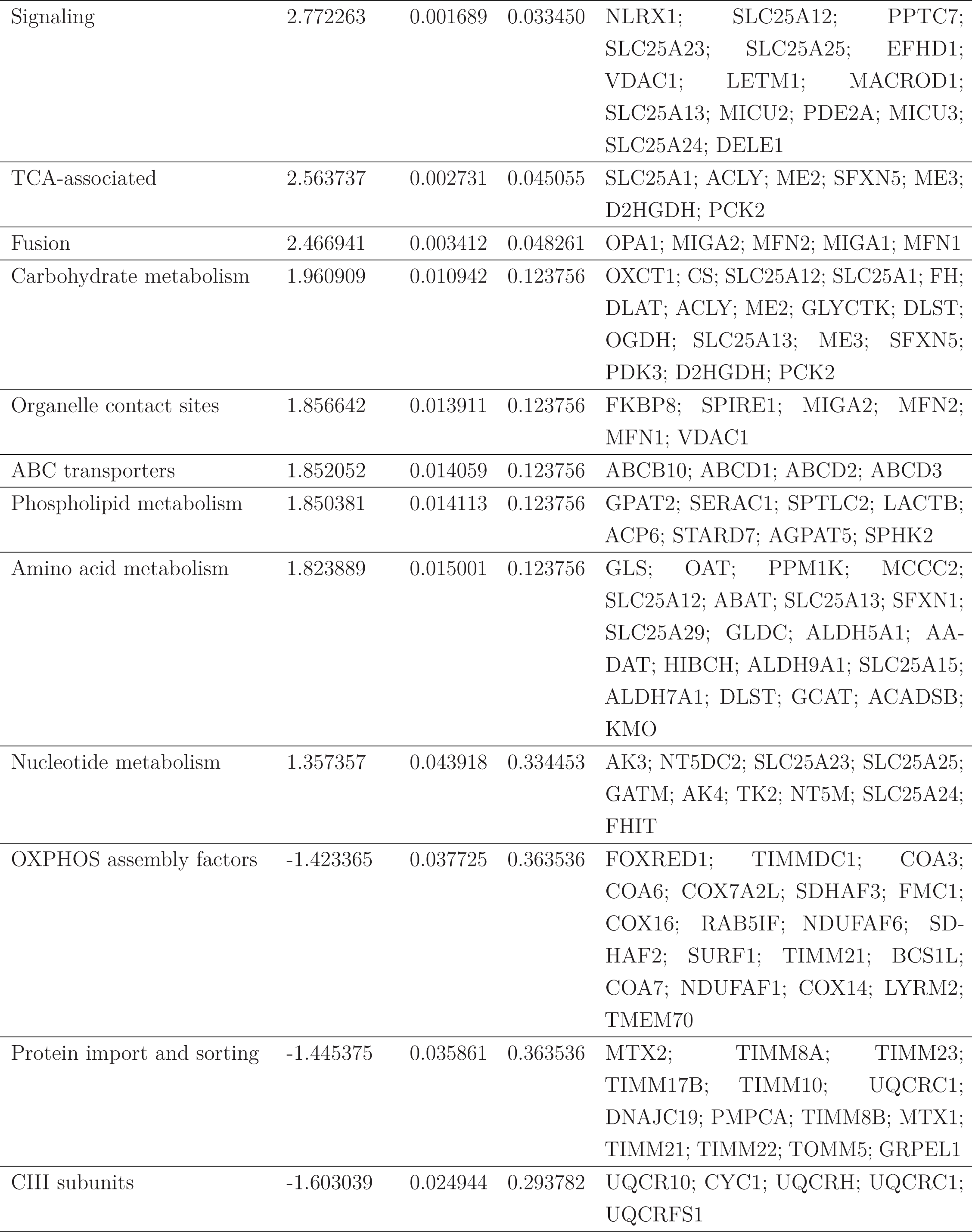
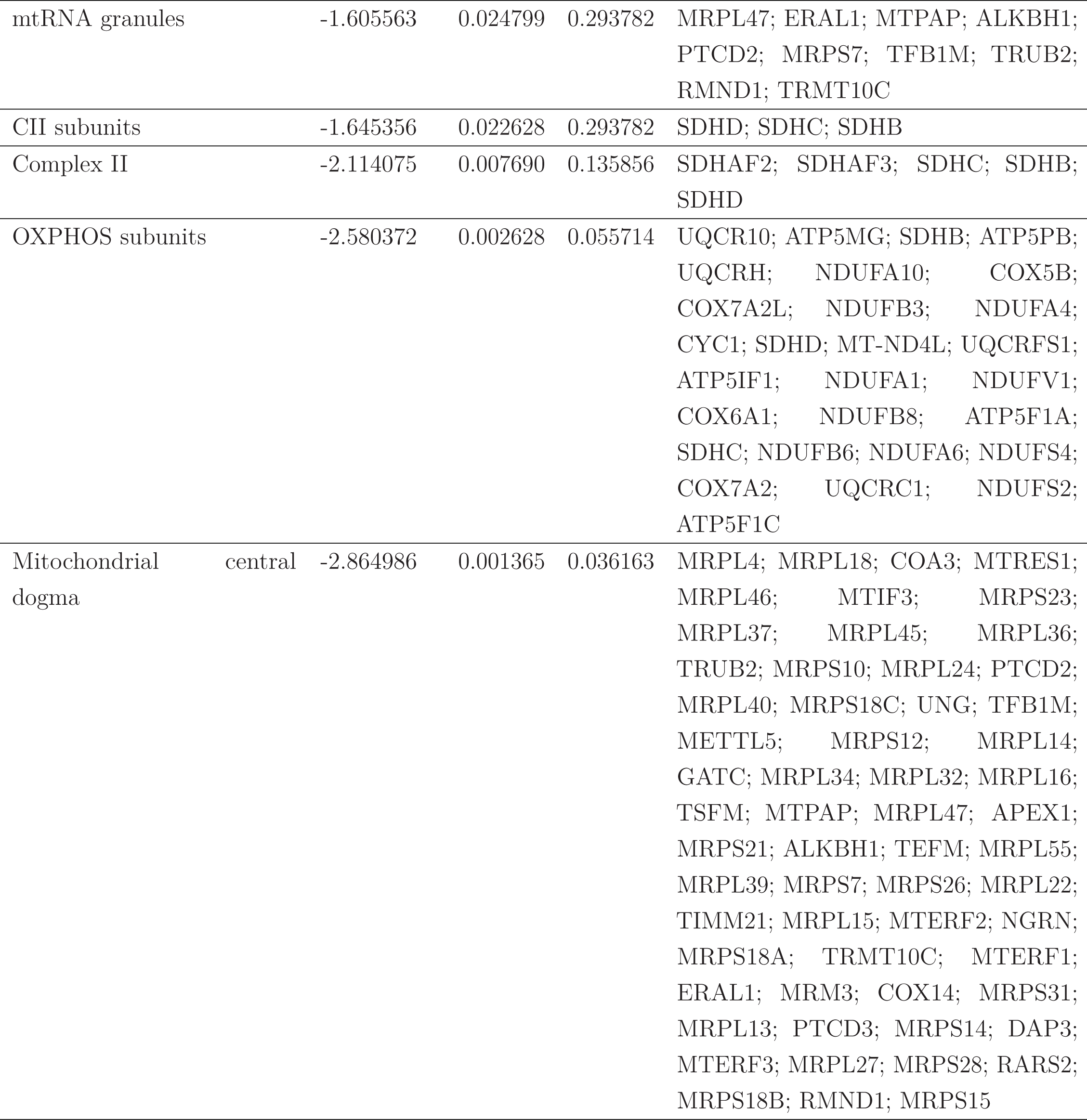
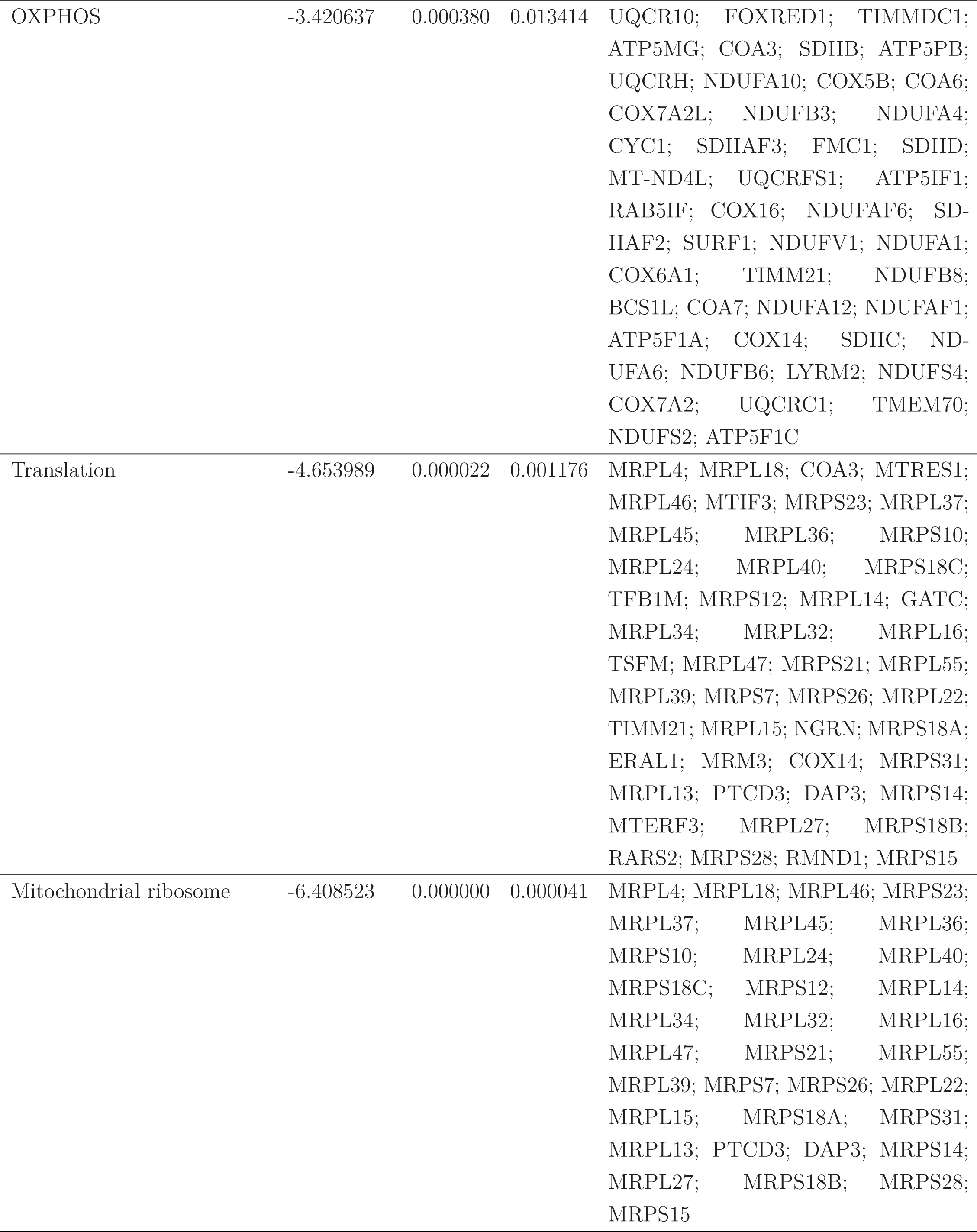
Enrichments of MitoCarta mitochondrial pathways in p.Tyr622* + H20 vs p.Tyr622* + CDP-choline mRNA.

**Table S6:**
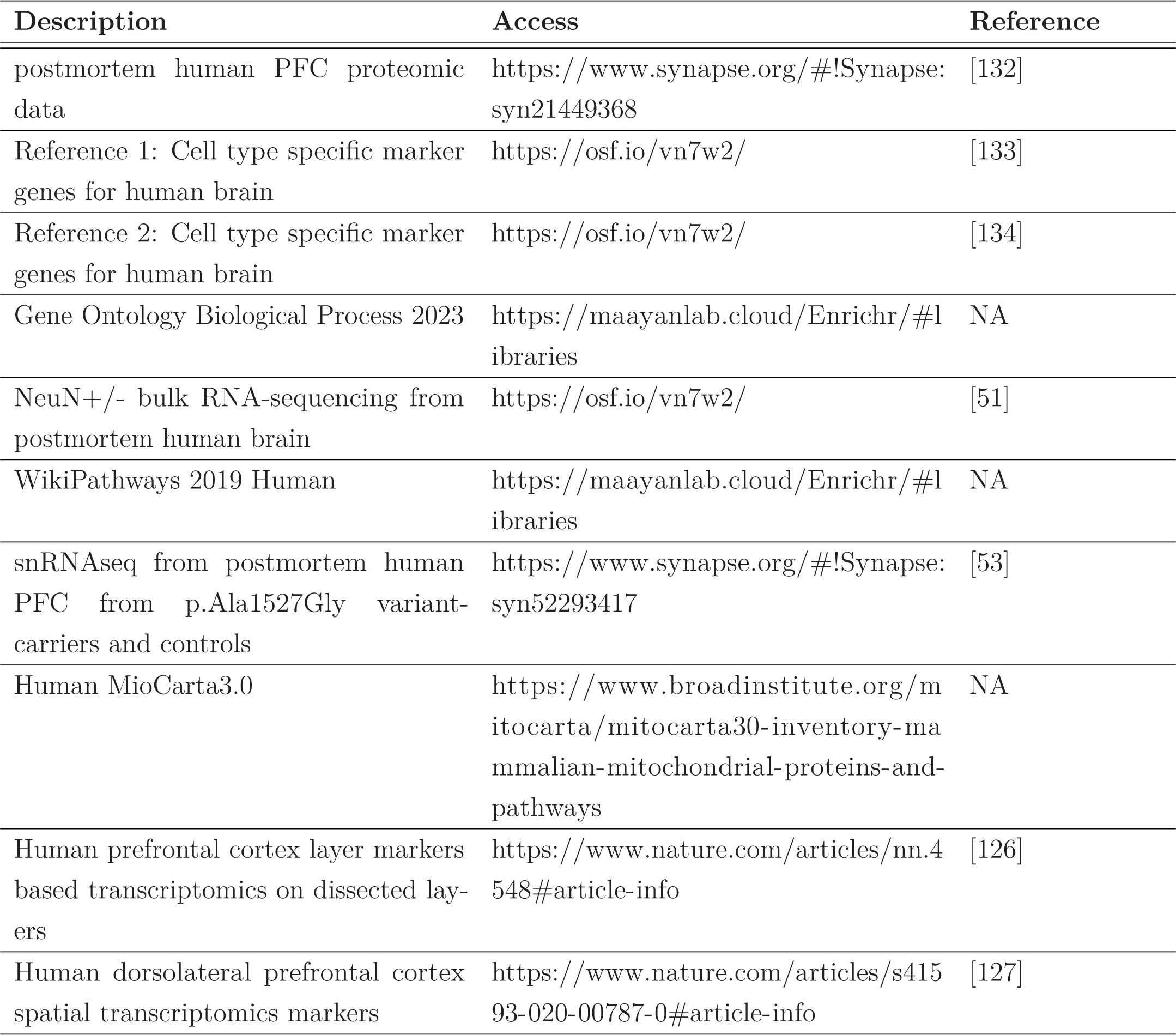
External datasets used.

**Table S7:** Antibodies used.

| **Antibody name** | **Company** | **Catalog No.** |
| --- | --- | --- |
| NeuN | Synaptic Systems | 266004 |
| Tuj1 | BioLegend | MMS-435P |
| SM312 (pan-axonal marker) | Biolegend | 837904 |
| MAP2 | Biolegend | 822501 |

## Extended Data

**Data S1.** ABCA7 PTC variants identified in the snRNA-seq cohort

**Data S2.** Metadata for selected individuals from the ROSMAP cohort used in this study

**Data S3.** snRNA-seq quality control metrics

**Data S4.** Differential gene expression statistics by cell type **Data S5.** Gene Ontology (GO) pathway enrichment results **Data S6.** WikiPathways enrichment results in excitatory neurons **Data S7.** Kernighan-Lin (K/L) gene cluster assignments

**Data S8.** RNA-seq differential expression statistics in NGN2-induced neurons

**Data S9.** Lipidomics differential abundance statistics in NGN2-induced neurons

**Data S10.** Targeted metabolomics on cells and media, with and without CDP-choline treatment

